# Distinct functions of three chromatin remodelers in activator binding and preinitiation complex assembly

**DOI:** 10.1101/2022.03.07.483278

**Authors:** Yashpal Rawal, Hongfang Qiu, Alan G. Hinnebusch

**Affiliations:** Division of Molecular and Cellular Biology, Eunice Kennedy Shriver National Institute of Child Health and Human Development, National Institutes of Health, Bethesda, Maryland 20892

**Author notes:** To whom correspondence should be addressed: YR: Dept. of Biochemistry and Structural Biology, University of Texas Health Science Center, San Antonio, TX 78229, USA; and. AGH: National Institutes of Health, Building 6, Room 230, Bethesda, MD 20892.; E- mail.

**Keywords:** RSC, Ino80, SWI/SNF, Gcn4, TBP, transcription, yeast

## Abstract

The nucleosome remodeling complexes (CRs) SWI/SNF, RSC, and Ino80C cooperate in evicting or repositioning nucleosomes to produce nucleosome depleted regions (NDRs) at the promoters of many yeast genes induced by amino acid starvation. We analyzed mutants depleted of the catalytic subunits of these CRs for binding of transcriptional activator Gcn4 and recruitment of TATA-binding protein (TBP) during preinitiation complex (PIC) assembly. RSC and Ino80 were found to enhance Gcn4 binding to both UAS elements in NDRs upstream of promoters and to unconventional binding sites within nucleosome-occupied coding sequences; and SWI/SNF contributes to UAS binding when RSC is depleted. All three CRs are actively recruited by Gcn4 to most UAS elements and appear to enhance Gcn4 binding by reducing nucleosome occupancies at the binding motifs, indicating a positive regulatory loop. SWI/SNF acts unexpectedly in WT cells to prevent excessive Gcn4 binding at many UAS elements, indicating a dual mode of action that is modulated by the presence of RSC. RSC and SWI/SNF collaborate to enhance TBP recruitment at Gcn4 target genes, together with Ino80C, in a manner associated with nucleosome eviction at the TBP binding sites. Cooperation among the CRs in TBP recruitment is also evident at the highly transcribed ribosomal protein genes, while RSC and Ino80C act more broadly than SWI/SNF at the majority of other constitutively expressed genes to stimulate this step in PIC assembly. Our findings indicate a complex interplay among the CRs in evicting promoter nucleosomes to regulate activator binding and stimulate PIC assembly.

**AUTHOR SUMMARY:** ATP-dependent chromatin remodelers (CRs), including SWI/SNF and RSC in budding yeast, are thought to stimulate transcription by repositioning or evicting promoter nucleosomes, and we recently implicated the CR Ino80C in this process as well. The relative importance of these CRs in stimulating activator binding and recruitment of TATA-binding protein (TBP) to promoters is incompletely understood. Examining mutants depleted of the catalytic subunits of these CRs, we determined that RSC and Ino80C stimulate binding of transcription factor Gcn4 to nucleosome-depleted regions, or linkers between genic nucleosomes, at multiple target genes activated by Gcn4 in amino acid-starved cells, frequently via evicting nucleosomes from the Gcn4 binding motifs. At some genes, SWI/SNF functionally complements RSC, while opposing RSC at others to limit Gcn4 binding. The CRs in turn are recruited by Gcn4 to establish positive or negative feedback loops that control Gcn4 binding. The three CRs also cooperate to enhance TBP recruitment, again involving nucleosome depletion, at both Gcn4 target and highly expressed ribosomal protein genes, whereas only RSC and Ino80C act broadly throughout the genome to enhance this key step in preinitiation complex assembly. Our findings illuminate functional cooperation among multiple CRs in regulating activator binding and promoter activation.

## INTRODUCTION

In the yeast *Saccharomyces cerevisiae*, most genes transcribed by RNA Polymerase II (Pol II) display a stereotypical pattern of nucleosome organization with a nucleosome-depleted region (NDR) of ∼120bp situated upstream of the coding sequences (CDS) and flanked by highly positioned “-1” and “+1” nucleosomes, with the transcription start site (TSS) generally located within the +1 nucleosome.

Promoter elements and upstream activation sequences (UAS elements) reside within the NDR and may extend upstream into the -1 nucleosome (-1_Nuc) (Jiang and Pugh 2009; Wang et al. 2011; Cui et al.

2012; Rando and Winston 2012). The exclusion of nucleosomes from the NDRs can facilitate efficient binding of transcriptional activator proteins at UAS elements (Devlin et al. 1991; Yu and Morse 1999) (Levo et al. 2017). The location of the TSS within the +1 nucleosome dictates that the latter is frequently evicted (Boeger et al. 2003; Reinke and Horz 2003), or shifted in the 3’ direction (Reja et al. 2015; Nocetti and Whitehouse 2016) (Rawal et al. 2018a), during assembly of the Pol II transcription preinitiation complex (PIC).

Promoter nucleosome organization arises in part from cooperative or antagonistic actions of ATP-dependent chromatin remodeling (CR) complexes (Krietenstein et al. 2016) (Kubik et al. 2019), which are recruited to NDRs or promoter-proximal nucleosomes of many yeast genes (Yen et al. 2012; Kubik et al. 2019). The CRs RSC and SWI/SNF generally move the -1 and +1 nucleosomes away from NDRs. RSC has been shown to function in this way at the majority of yeast genes (Badis et al. 2008; Parnell et al. 2008; Hartley and Madhani 2009) to maintain native NDR widths (Ganguli et al. 2014; Parnell et al. 2015). SWI/SNF, in contrast, acts in the promoters of a small fraction of genes that tend to have poor nucleosome phasing and wide NDRs (Ganguli et al. 2014; Kubik et al. 2019) and to be highly expressed in WT cells (Rawal et al. 2018a); and SWI/SNF and RSC exhibit functional redundancy at such genes (Rawal et al. 2018a; Kubik et al. 2019). The Ino80 complex (Ino80C) can have opposite effects on nucleosome positioning at different genes, and acts differently from RSC and SWI/SNF at certain genes to move the +1 nucleosome upstream to narrow, rather than widen, the NDR (van Bakel et al. 2013; Kubik et al. 2019). This Ino80C activity is much more pronounced on nuclear depletion of Isw2, catalytic subunit of the CR ISW2; and simultaneous depletion of both Ino80 and Isw2 leads to wider NDRs at a large number of genes and suppresses the widespread narrowing of NDRs conferred by depleting the RSC catalytic subunit Sth1 (Kubik et al. 2019). There is evidence that Ino80C also “edits” promoter nucleosomes to replace histone variant H2A.Z with conventional H2A (Brahma et al. 2017); however, there are conflicting findings regarding this activity (Jeronimo et al. 2015). Other evidence indicates that Ino80C can function in remodeling conventional nucleosomes containing H2A versus H2A.Z (Mizuguchi et al. 2004; Rosonina et al. 2014) (Qiu et al. 2020).

Transcriptional activation of yeast genes is mediated by binding of activator proteins to UAS elements that recruit an array of co-factors to facilitate binding of general transcription factors (GTFs) to the promoter, including TATA-binding protein (TBP), and in recruitment of Pol II for PIC assembly (Rando and Winston 2012). In addition to acting as mediators to bridge activators with GTFs or Pol II, co-factors can function to evict or re-position nucleosomes that occlude the promoter or TSS (Rhee and Pugh 2012). CRs can function in this way, and inactivation of RSC confers a widespread reduction in expression of many genes (Parnell et al. 2008), particularly those with intermediate to low expression levels in WT cells (Klein-Brill et al. 2019). RSC, in cooperation with general regulatory factors (GRFs) Reb1, Abf1 and Rap1, appears to slide the +1 nucleosome downstream to enhance TBP binding to the promoters of many genes (Kubik et al. 2018). SWI/SNF partners with RSC (Rawal et al. 2018a) and histone acetyltransferase Gcn5 (Qiu et al. 2016) in stimulating transcription of many highly transcribed, constitutively expressed genes. SWI/SNF was also implicated in transcriptional activation of condition- regulated genes such as *PHO*5 (Barbaric et al. 2007), *SUC2* (Schwabish and Struhl 2007), *RNR1* (Sharma et al. 2003), various genes involved in metabolic reprogramming (Nocetti and Whitehouse 2016), and genes activated by heat-shock (Shivaswamy and Iyer 2008) or amino acid starvation (Rawal et al. 2018a).

For many genes in this last group, mostly induced by transcriptional activator Gcn4, SWI/SNF partners with RSC to evict promoter nucleosomes and reposition the remaining -1 and +1 nucleosomes to widen the NDRs and promote transcription (Rawal et al. 2018a). There is evidence that downstream repositioning of the +1 nucleosome by SWI/SNF enhances TBP binding to the promoters at a subset of SWI/SNF-dependent genes (Kubik et al. 2019).

Ino80C is required for efficient transcriptional activation of genes induced in response to inositol depletion (Kubik et al. 2019), and it partners with SWI/SNF and RSC in nucleosome remodeling during induction of the *PHO5* gene (Barbaric et al. 2007; Musladin et al. 2014) and at a subset of genes induced by amino acid starvation (Qiu et al. 2020). For the latter, Ino80C promotes eviction of promoter nucleosomes and appears to stimulate transcription by enhancing TBP recruitment. Ino80C is also critical for expression of TORC1-responsive genes during the yeast metabolic cycle (YMC) and thus helps to coordinate respiration and cell division with periodic gene expression (Gowans et al. 2018). The frequent downstream repositioning of the +1 nucleosome on co-depletion of Ino80 and ISW2 mentioned above is associated with enhanced TBP binding and elevated transcription, often involving cryptic TSSs normally occluded by the +1 nucleosome (Kubik et al. 2019). There is also evidence that Ino80 acts to restore, rather than remove, promoter nucleosomes following their rapid eviction in response to osmotic stress, and thereby prevent prolonged transcriptional activation of certain stress-induced genes (Klopf et al. 2017). Another study indicates that Ino80C is recruited by TBP to both promoters and terminators of Pol II genes, in association with TBP-binding factors Mot1 and NC2, where they act to suppress cryptic promoters in intergenic regions and also some physiological promoters at transcriptionally inactive genes (Xue et al. 2017).

While there is evidence that CRs influence transcriptional activation by evicting or displacing promoter nucleosomes to control access of GTFs and attendant PIC assembly, much less is known about their importance in regulating the binding of transcriptional activators to the UAS elements of yeast genes. In vitro evidence from single-molecule analysis indicates that RSC promotes binding of transcriptional activator Ace1 to the *CUP1* promoter by increasing accessibility of the Ace1 binding site in chromatin to reduce the search time for Ace1 binding (Mehta et al. 2018). SWI/SNF remodeling activity is critical for efficient binding of activator Pho4 at the *PHO5* UAS (Adkins et al. 2007). On the other hand, in vitro single-molecule studies indicate that SWI/SNF can impede binding of the Gal4 DNA binding domain by sliding a nucleosome across its binding site (Li et al. 2015). Moreover, human SWI/SNF was shown in vitro to displace the glucocorticoid receptor from its binding site in a reconstituted nucleosome array, likely contributing to the transient nature of GR interactions with the promoter in chromatin (Nagaich et al. 2004). Thus, it seems that SWI/SNF can either enhance or impede activator binding in different settings.

Depriving yeast cells of an amino acid, including starvation for isoleucine and valine, achieved with the inhibitor sulfometuron methyl (SM), increases the transcription of hundreds of genes, most of which are dependent on activator Gcn4 for induction (Jia et al. 2000; Natarajan et al. 2001; Saint et al. 2014). Previously, by ChIP-Seq analysis of Pol II subunit Rpb3, we identified a group of ca. 200 genes exhibiting ≥2-fold induction of Pol II occupancies averaged across the CDS on SM treatment. Parallel ChIP-Seq analysis of histone H3 revealed a marked eviction of nucleosomes in the promoter intervals spanning the -1 and +1 nucleosomes and intervening NDRs at 70 of these SM-induced genes, and lesser increases in H3 occupancies at the remaining 134 induced genes (Qiu et al. 2016). As noted above, examining mutants lacking the catalytic subunits of SWI/SNF (*snf2Δ*) or Ino80C (*ino80Δ*), or conditionally depleted of the essential catalytic subunit of RSC by transcriptional shut-off (*PTET-STH1*), revealed overlapping roles for these three CRs in evicting promoter nucleosomes and inducing transcription of SM-induced genes (Qiu et al. 2016; Rawal et al. 2018a; Qiu et al. 2020). Here, we set out to examine their roles in binding of Gcn4 to UAS elements, which increases sharply on translational induction of Gcn4 protein by SM (Rawal et al. 2018b).

Interestingly, our previous ChIP-seq analysis of Gcn4 in SM-treated WT cells revealed that only ∼40% of the 546 sites of induced binding detected throughout the genome are located upstream of genes (5’ sites), with the majority occurring instead within the CDS (ORF sites). Roughly 70% of the genes with conventional 5’ sites show evidence of transcriptional activation in amino acid starved cells and are likely direct targets of Gcn4, whereas only ∼20% of the genes with only ORF binding sites are transcriptionally activated. Mutation of the internal Gcn4 binding sites at a number of the latter genes was shown to impair their transcriptional activation by SM, demonstrating that Gcn4 can activate 5’- positioned promoters by binding within the downstream ORFs. Gcn4 binding to the majority of 5’ sites occurs within NDRs; while the binding at ORF sites generally occurs within linkers between genic nucleosomes. Moreover, only ∼30% of all predicted Gcn4 binding motifs in the genome are occupied by Gcn4 in SM-induced cells, and the two most important features of bound motifs are (i) a strong match to the consensus Gcn4 motif and (ii) lower than average nucleosome occupancy (Rawal et al. 2018b). These findings suggest that Gcn4 binding is inefficient when its binding motif maps within a nucleosome, in agreement with previous findings that Gcn4 binding to the *HIS4* UAS is enhanced by Rap1 and its ability to exclude nucleosomes (Devlin et al. 1991; Yu and Morse 1999). Nucleosome occupancy is also a key determinant of binding by activator Leu3 in the yeast genome (Liu et al. 2006). Recent results indicating cooperation between different GRFs and RSC in downstream positioning of the +1 nucleosome for NDR formation (Kubik et al. 2018), led us to wonder whether Gcn4 binding to 5’ sites at many UASs might be stimulated by RSC.

The results of the current study indicate that Gcn4 binding at many 5’ sites, as well as ORF sites, is stimulated by RSC, by SWI/SNF in cells depleted of RSC, and by Ino80C. Surprisingly, when RSC is present, SWI/SNF acts primarily to limit rather than enhance Gcn4 occupancies at 5’ sites, suggesting a dual role for SWI/SNF in activator binding. We found that Gcn4 recruits all three CRs to 5’ sites, consistent with their direct roles, and indicating feedback loops that regulate Gcn4 binding at UAS elements. Other evidence indicates that the CRs enhance Gcn4 binding by reducing nucleosome occupancies of the Gcn4 binding motifs, supporting the notion that Gcn4 binding is impeded by nucleosomes. Examining the effects of deleting/depleting the CRs on TBP occupancies showed that RSC and SWI/SNF collaborate to enhance TBP recruitment at Gcn4 target genes, as does Ino80C, in a manner associated with nucleosome eviction at the TBP binding sites. Cooperation among the CRs is also evident at the highly transcribed ribosomal protein genes (RPGs), while RSC and Ino80C act more broadly than SWI/SNF to stimulate this step in PIC assembly throughout the rest of the genome.

## RESULTS

### RSC, SWI/SNF and Ino80C modulate Gcn4 binding within NDRs and coding regions

Our previous ChIP-seq analysis of Gcn4 in SM-induced WT cells identified 117 Gcn4 binding sites located 5’ of an annotated TSS in the canonical location of yeast UAS elements, which appear to mediate transcriptional activation of the adjacent genes during amino acid starvation (<u>5</u>’ Gcn4 peaks). An additional 62 Gcn4 peaks were identified within coding sequences that appear to activate transcription of the full-length transcript of the corresponding gene or an adjacent gene lacking a Gcn4 peak (ORF peaks) (Rawal et al. 2018b). As reported previously (Rawal et al. 2018b), averaging the Gcn4 occupancies measured by ChIP-seq over the 5’ or ORF peaks and aligning them to the Gcn4 binding motifs reveals that Gcn4 occupancies are highly induced by SM in WT cells, and that the summits of the occupancy peaks coincide precisely with the Gcn4 binding motifs (Fig. 1A(i) & 1B(i), yellow vs. blue).

**Figure 1.**
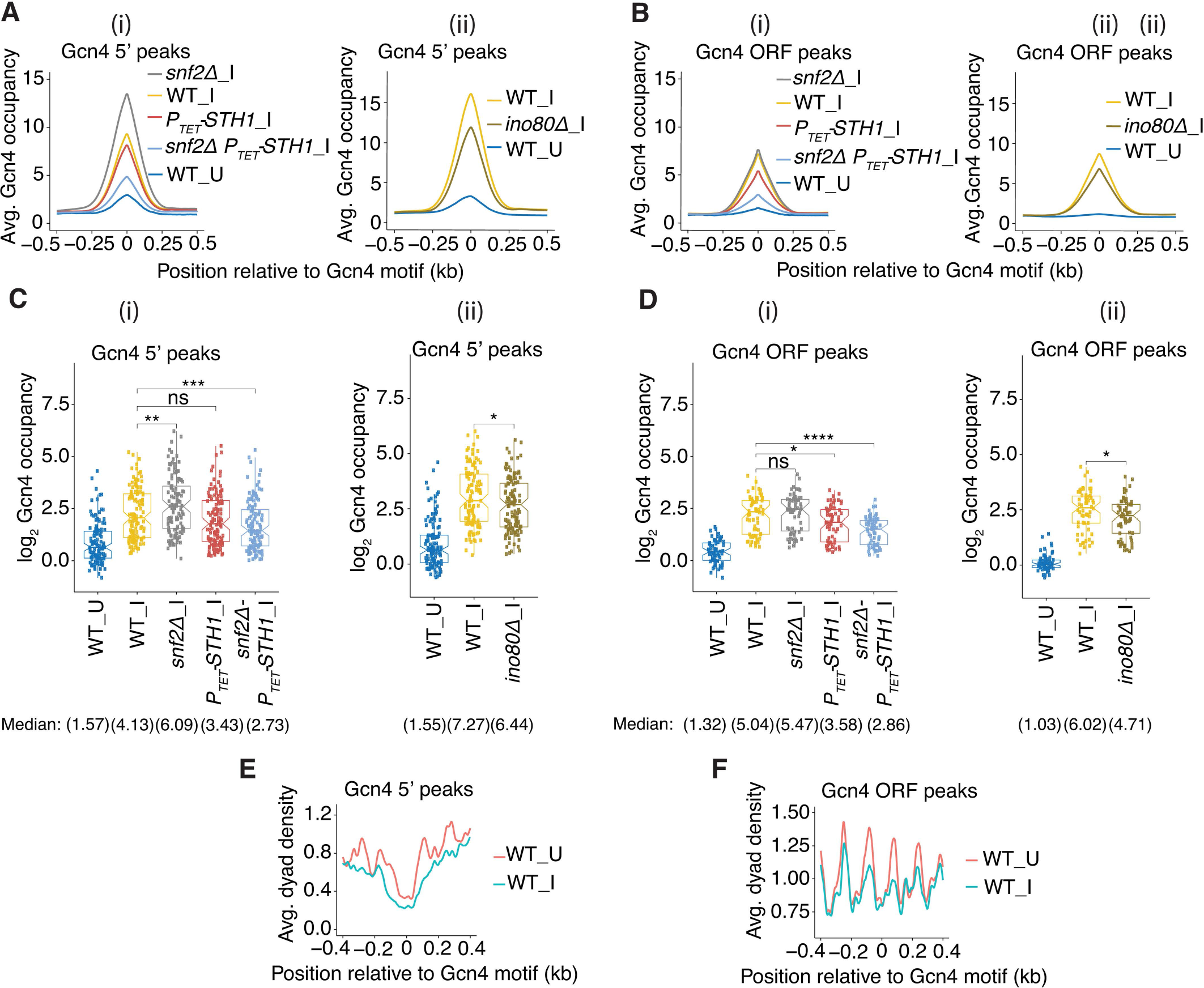
Gcn4 occupancy changes in CR mutants. **(A-B)** Gcn4 occupancies at each base pair surrounding the consensus binding motifs averaged over (A) all 117 5’ sites or (B) all 62 ORF Gcn4 peaks in WT_U (blue), WT_I (yellow), and SM-induced CR mutants for (i) *snf2ý*_I (gray), *PTET-STH1_*I (red), *snf2ý PTET- STH1_*I (cyan); and (ii) WT_I (yellow) and *ino80ý*_I (gold). In these and all similar plots below, yeast strain/condition labels adjacent to the tracings are listed in decreasing order of summit heights. (**C-D**) Notched box plots of log2 Gcn4 occupancies per nucleotide averaged over the peak coordinates assigned by MACS2 analysis of (C) 5’ and (D) ORF Gcn4 peaks for (i) WT_U, WT_I, and CR mutants *snf2ý*_I, *PTET-STH1_*I and *snf2ý PTET-STH1_*I; and (ii) WT_U, WT_I, and *ino80ý*_I samples, color-coded as in (A-B). For these and all subsequent box plots, each box depicts the interquartile range containing 50% of the data, intersected by the median; the notch indicates a 95% confidence interval (CI) around the median, and *p* values for the significance of differences in median values calculated by the Mann-Whitney-Wilcoxon test are indicated as follows: ****, ≤ 0.0001; ***, ≤ 0.001; **, ≤ 0.01; *, ≤ 0.05; ns, > 0.05. Unlogged median Gcn4 occupancies are indicated under the respective strain labels. Data analyzed in (A-D) were calculated from Gcn4 ChIP-seq analysis of sonicated chromatin from 2-4 biological replicates and normalized to the average occupancy per nucleotide on each chromosome for each data set. **(E-F)** Average dyad densities calculated from H3 MNase-ChIP-seq data aligned to the Gcn4 consensus motifs for (B) 5’ and (D) ORF Gcn4 peaks in WT cells either untreated (_U) or SM-treated (_I). Midpoints (dyads) of nucleosome-size sequences between 125 and 175 bp were mapped with respect to Gcn4 consensus motifs. Average profiles were smoothed using a moving average filter with a span of 31 bp. The data were normalized to the average occupancy per nucleotide on each chromosome for each data set.

Our previous ChIP-seq analysis of histone H3 using chromatin fragmented by micrococcal nuclease digestion, which preferentially cuts within linkers, allowed us to estimate the positions of nucleosome dyads from the midpoints of immunoprecipitated DNA fragments. Averaging dyad densities over all 5’ or ORF peaks reveals that Gcn4 binding motifs of 5’ sites are generally located in the centers of NDRs, whereas the motifs for ORF peaks reside in the linkers between the genic nucleosomes observed in untreated WT cells (Fig. 1E-F, red tracings). The dyad densities of the -1 and +1 nucleosomes and the intervening NDRs at the 5’ Gcn4 peaks are diminished, and the NDRs become wider, in response to SM treatment (Fig. 1E, blue vs. red), reflecting eviction and repositioning of the -1 and +1 nucleosomes (Rawal et al. 2018a). The dyad densities of the genic nucleosomes flanking ORF peaks are also diminished; however, the spacing between them appears unaltered by SM treatment (Fig. 1F, blue vs. red). The fact that Gcn4 binds preferentially to motifs located outside of nucleosomes, and our previous findings that eviction or repositioning of promoter nucleosomes induced by SM treatment involves SWI/SNF, RSC and Ino80C (Rawal et al. 2018a; Qiu et al. 2020), led us to examine whether Gcn4 binding is enhanced by one or more of these CRs.

To this end, we conducted Gcn4 ChIP-seq analysis on *snf2Δ* and *ino80Δ* mutants, lacking the catalytic subunits of SWI/SNF or Ino80C, respectively, and on a *PTET-STH1* strain in which the RSC catalytic subunit was transcriptionally repressed by addition of doxycycline (Dox) for 8h, in cells treated with SM to induce Gcn4. We similarly examined a *snf2Δ PTET-STH1* double mutant treated with Dox to examine the effects of depleting RSC in cells lacking SWI/SNF. We observed a slight reduction in the averaged Gcn4 occupancies for the group of 5’ sites on depleting Sth1 in *PTET-STH1* versus WT cells under inducing conditions (*PTET-STH1*_I vs. WT_I), but a more substantial reduction of ∼50% in the induced *snf2Δ PTET-STH1* double mutant versus WTI cells (Fig. 1A(i), red & cyan vs. yellow). Surprisingly, in the *snf2Δ* single mutant, the averaged Gcn4 occupancy was markedly increased compared to WT cells (Fig. 1A(i), grey vs. yellow). Calculating the mean Gcn4 occupancy per nucleotide across each 5’ site (with boundaries defined as described in Materials and Methods), revealed a significant reduction in median Gcn4 occupancy in *snf2Δ PTET-STH1*_I cells, but an increased median occupancy in *snf2Δ_*I cells, compared to WT_I cells (Fig. 1C(i), cols. 5 & 3 vs. 2). Similar analyses showed that the *ino80Δ* mutation conferred moderate reductions in both average and median Gcn4 5’ occupancies (Fig. 1A(ii) & 1C(ii), gold vs. yellow), which is less severe than that just described for the *snf2Δ PTET-STH1* double mutant. The ORF Gcn4 peaks also showed reductions in *PTET-STH1*_I and *snf2Δ PTET-STH1_*I cells, but no significant increase in *snf2Δ*_I cells (Fig. 1B(i) and 1D(i)). The *ino80Δ* mutation reduced Gcn4 occupancies at ORF peaks as well (Fig. 1B(ii) and 1D(ii)), to an extent comparable to that observed for the *PTET-STH1*_I single mutation (Fig. 1B(i) and 1D(i)).

To explore further the contributions of the CRs at individual Gcn4 motifs, we sorted the 5’ Gcn4 peaks according to their reductions in Gcn4 occupancies in the *snf2Δ PTET-STH1*_I double mutant compared to WT_I cells. A heat map showing the differences in Gcn4 occupancies between the double mutant and WT for this ordering of 5’ sites is shown in Fig. 2A(i), which was calculated from the corresponding occupancies displayed for these two strains as displayed in the heat maps of Fig. 2B (i)-(ii). Thus, the Gcn4 5’ sites in the top quartile of the maps (Set_1 in Fig. 2B) exhibit the greatest occupancy reductions in the double mutant versus WT cells, as indicated by the dark blue hues in the difference map of Fig. 2A(i). Similar difference maps were generated for the *PTET-STH1* or *snf2Δ* single mutants, shown in Fig. 2A(ii)- (iii), based on the Gcn4 occupancies measured in these strains displayed in Figs. 2B (iii) or (iv). The Set_1 peaks display relatively smaller occupancy decreases in the *PTET-STH1*_I single mutant vs. WT_I than was observed in the double mutant (lighter blue hues in the difference map of Fig. 2A(ii) vs. Fig. 2A(i) for the top quartile). In *snf2Δ*_I cells, the Set_1 peaks variously show small decreases, no change, or small increases in Gcn4 occupancy compared to WT_I cells (Fig.2A (iii), top quartile). The Gcn4 peaks in the middle two quartiles of the maps (dubbed Set_2 in Fig. 2B) show strong to moderate reductions in the double mutant, lesser reductions in the *PTET-STH1*_I single mutant, but have generally increased occupancies in *snf2Δ*_I versus WT_I cells (Fig. 2A(i)-(iii), middle quartiles). The genes in the bottom quartile of the maps (Set_3) tend to show similar, small occupancy increases or decreases in both the double and single *PTET-STH1*_I mutants (Fig. 2A(i) & (ii)), while showing marked increases in Gcn4 occupancies in *snf2Δ*_I cells (Fig. 2A(iii)).

**Figure 2.**
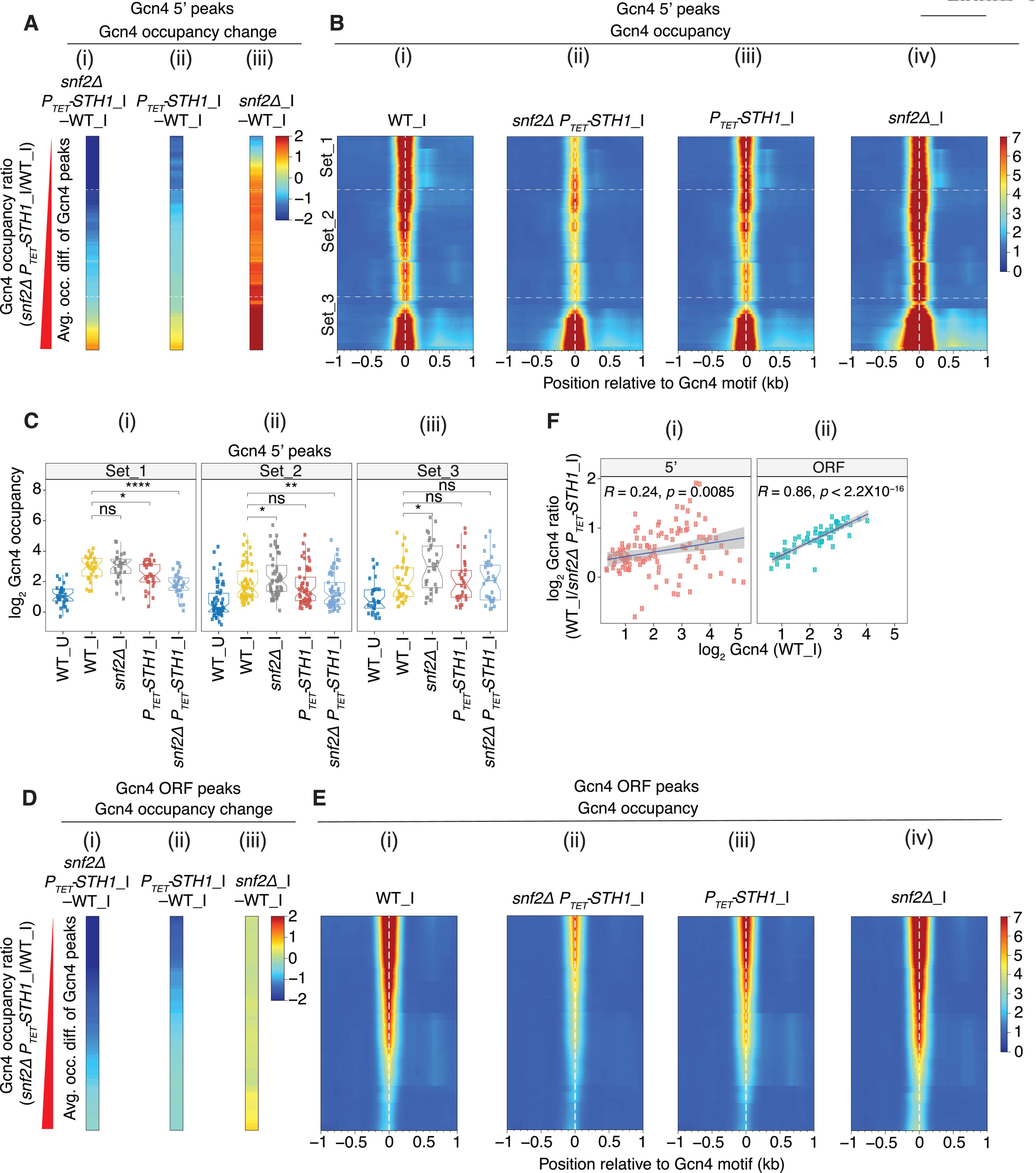
SWI/SNF and RSC have opposing effects on Gcn4 binding at 5’ sites. (A) **Heat maps of** differences in Gcn4 occupancies averaged across the coordinates of 5’ sites between the indicated mutant and WT_I samples for (i) *snf211 PTET-STH1_*I, (ii) *PTET-STH1_*I and (iii) *snf211*_I cells. Gcn4 5’ sites were sorted by increasing order of the ratio of Gcn4 occupancies in the double mutant *snf211 PTET-STH1_*I vs. WT_I. The peaks belonging to the first (Set_1, n=30), middle two (Set_2, n=57) and fourth (Set_3, n=30) quartiles of fold-changes are arranged from top to bottom and separated by white lines across the maps. In these and all heat maps below, color-coding of values in the heat maps are shown in a key to the right of the map(s). **(B)** Heat map depictions of Gcn4 occupancies surrounding the Gcn4 motifs of 5’ sites in (i) WT_I, (ii) *snf211 PTET-STH1_*I, (iii) *PTET-STH1_*I, and (iv) *snf211*_I cells, for the same ordering of 5’ sites as in (A). The locations of 5’ Gcn4 peaks in Sets_1 to 3 are indicated. **(C)** Notched box pots of log2 Gcn4 occupancies in WT_U, WT_I, or *snf211*_I, *PTET-STH1_*I and *snf211 PTET-STH1_*I cells in 3 sets of Gcn4 5’ sites comprised of the (i) first (Set_1, n=30), (ii) middle two (Set_2, n=57) and (iii) last (Set_3, n=30) quartiles of the fold-changes in Gcn4 occupancy in *snf211 PTET-STH1_*I vs. WT_I cells as depicted in Fig. 2B(i). **(D & E)** Same analyses shown in (A & B) except for Gcn4 ORF peaks. **(F)** Scatterplot of log2 ratios of Gcn4 occupancy changes in WT_I vs. *snf211 PTET-STH1_*I plotted against log2 Gcn4 occupancies in WT_I cells for (i) 5’ (left panel) and (ii) ORF (right) Gcn4 peaks. Pearson correlation coefficients (*R*) and associated *p* values are indicated.

The trends depicted in the heat maps of Fig. 2A-B were confirmed by quantifying the Gcn4_I occupancies in each peak and calculating the median values for the three sets of peaks. As expected, Set_1 displays a greater reduction in Gcn4 median occupancy in the double mutant versus the *PTET-STH1*_I single mutant, but no significant change in the *snf2Δ* single mutant (Fig. 2C(i), cols. 3-5 vs. 2). Set_2 also shows a greater reduction in median occupancy in the double mutant versus the *PTET-STH1*_I single mutant, even though Gcn4 occupancy is significantly elevated in the *snf2Δ* single mutant containing *STH1* (Fig. 2C (ii), cols. 3-5 vs. 2). Set_3 shows no significant decrease in median Gcn4 occupancy in either of the *PTET-STH1*_I mutants, but a strong increase in the *snf2Δ* single mutant (Fig. 2C(iii), cols. 3-5 vs. 2). The results in Figs. 2A-C were analyzed further by plotting the changes in Gcn4 occupancies at each 5’ site between WT and mutant cells. For Set 1 and Set 2, all peaks show moderate reductions in Gcn4 occupancies in *PTET-STH1*_I versus WT_I cells (Fig. S1A), which are exacerbated by deletion of *SNF2* in the double mutant compared to the *PTET-STH1* single mutant (Fig. S1B). The Set 3 peaks exhibit variable occupancy changes in both the *PTET-STH1* single and double mutant versus WT (Fig. S1A-B). In contrast, the *snf2Δ* single mutation confers variable changes for Set 1 sites, but moderate or strong increases in Gcn4 occupancies for Set 2 and Set 3 sites, respectively (Fig. S1C).

Together, these findings indicate that RSC enhances Gcn4 binding for Set_1 and Set_2 peaks, with a greater contribution for Set_1, but has weak and variable effects on Set_3 sites. In WT cells, SWI/SNF generally inhibits, rather than promotes, Gcn4 binding at the Set_2 and _3 sites, with the greatest inhibition for the Set_3 sites that are largely insensitive to RSC, but SWI/SNF has variable effects on the Set_1 sites that are most dependent on RSC. In cells depleted of RSC however, SWI/SNF promotes rather than inhibits binding at the 5’ sites in Set_1 and _2, thus compensating for reduced RSC function at the peaks most dependent on RSC; and SWI/SNF no longer inhibits Gcn4 binding to most peaks in Set_3. One way to explain these last findings for Set_3 would be to propose that the inhibitory effect of SWI/SNF on Gcn4 binding is offset by the stimulatory role that SWI/SNF exerts upon depletion of Sth1, with relatively little net change in Gcn4 occupancy found in the double mutant. The differing effects of the *PTET-STH1*_I and *snf2Δ* mutations on Gcn4 occupancies for the three Sets of 5’ sites described above are illustrated for two archetypal members of each Set in Figs. S2A-C ((i)-(ii)). It is worth noting that we previously obtained results fully consistent with those reported here by ChIP analysis of Gcn4 for particular genes, where occupancies were calculated as the percentage of input DNA recovered in the immunoprecipitates (Qiu et al. 2016). Thus, we reported that Gcn4 occupancy was elevated in *snf2Δ* versus WT cells by 2.1- and 1.6- fold in the UASs of Set3 genes *ARG1* and *ARG4*, respectively, elevated by 1.5-fold at the UAS of Set 2 gene *CPA2*, and did not increase at the Set 1 gene *HIS4*, as illustrated here for *ARG1* and *HIS4* in Figs. S2C & S2A, respectively.

The effects of the CR mutations on Gcn4 occupancies at binding sites in coding regions were less complex, as the majority of Gcn4 ORF peaks show relatively strong reductions in the *snf2Δ PTET-STH1* double mutant, moderate reductions in the *PTET-STH1*_I single mutant, and little change in the *snf2Δ*_I single mutant (see difference maps in Fig. 2D(i)-(iii), based on the heat maps in Fig. 2E(i)-(iv)). Thus, as noted for 5’ Gcn4 peaks, RSC generally stimulates Gcn4 binding and SWI/SNF partially compensates for RSC function on depletion of RSC, but there is no evidence for an inhibitory effect of SWI/SNF on Gcn4 binding at the ORF peaks. Interestingly, the magnitude of the reduction in Gcn4 occupancy at the ORF peaks in the *snf2Δ PTET-STH1* double mutant versus WT cells is strongly correlated with Gcn4 occupancy at these peaks in WT_I cells (Fig. 2F(ii)), suggesting an increasing requirement for RSC or SWI/SNF as the occupancy at ORF sites increases. A significant but considerably weaker correlation exists for 5’ Gcn4 peaks (Fig. 2F(i)), possibly reflecting the more complex interplay between RSC and SWI/SNF at the 5’ sites.

Analyzing the effects of *ino80Δ* for the same ordering of 5’ Gcn4 peaks described above reveals that the 5’ sites in Set_1 show reductions of lesser magnitude compared to those given by *snf2Δ PTET- STH1*; whereas comparable, smaller reductions are conferred by *ino80Δ* and *snf2Δ PTET-STH1* for Set_2 (Fig. S3A(i) vs. (ii) & S3B, Sets_1-2). In contrast, many of the Set_3 sites display reduced Gcn4 occupancies in *ino80Δ* cells but either little change or increased binding in *snf2Δ PTET-STH1* cells (Fig. S3A(i) vs. (ii) & S3B, Set_3), suggesting a greater dependence on Ino80C compared to RSC. For ORF peaks, *ino80Δ* confers a moderate reduction in Gcn4 binding across the spectrum of binding sites, similar to the effects of the *PTET-STH1* single mutation (Fig. S3C(i) vs. (ii) & S3D).

To identify all 5’ sites with a heightened dependence on Ino80C, we ordered them differently based on their decreased Gcn4 occupancies in *ino80Δ _*I versus WT_I cells, generating heat maps with peaks exhibiting the largest occupancy reductions conferred by *ino80Δ* at the top of the maps (Fig. S4A (i)-(iii)). As expected, *ino80Δ* substantially decreases the median occupancy for the top quartile of peaks (Set_I), but not for the lower quartiles in Sets_II or III (Fig. S4C(i)-(iii)). The corresponding difference maps for the *PTET-STH1* and *snf2Δ PTET-STH1* mutations for this same ordering of 5’ sites reveals more uniform reductions in Gcn4 occupancies across the sites compared to those given by *ino80Δ* (Fig. S4B(i)-(ii) vs. S4A(i)); and the *snf2Δ* single mutation generally confers increased Gcn4 occupancies for sites in Sets_I-II that show decreased occupancies in *ino80Δ _*I versus WT_I cells (Fig. S4B(iii) vs. S4A(i)). Thus, certain 5’ sites have a particularly strong dependence on Ino80C, but little requirement for RSC or SWI/SNF, for WT Gcn4 binding. The importance of Ino80C for WT Gcn4 binding is illustrated for the *YHR162W* and *HIS4* genes in Figs. S10E(i)-(ii).

We considered the possibility that the changes in Gcn4 occupancies in the mutants described above involve changes in Gcn4 expression. Quantifying the occupancies of Pol II across the coding sequences of *GCN4* from our previous ChIP-seq analysis of Rpb3 indicated similar reductions in *GCN4* transcription of ∼32-37% in the induced *snf2Δ PTET-STH1* and *PTET-STH1* mutants, but no change in *snf2Δ_*I or *ino80Δ_*I cells (Rawal et al. 2018a; Qiu et al. 2020) (Fig. S5A, cols. 3-6 vs. 2). Similarly, our Western analyses of Gcn4 protein levels indicated comparable reductions of ∼20% in the *PTET-STH1* and *snf2Δ PTET- STH1* mutants, but no change conferred by *snf2Δ* (Rawal et al. 2018a) (Fig. S5B) or *ino80Δ* (Qiu et al. 2020). The fact that the *snf2Δ PTET-STH1* and *PTET-STH1* mutations confer similar, moderate reductions in Gcn4 expression (Fig. S5A-B) suggests that the considerably larger reductions in Gcn4 occupancy observed for *snf2Δ PTET-STH1*_I versus *PTET-STH1*_I cells (Fig. 1A & 1C and Fig. 2A-E) do not result from decreased Gcn4 abundance.

To examine this issue further, we reasoned that if reduced Gcn4 occupancies arise primarily from reduced Gcn4 abundance, then the occupancy reductions should occur preferentially at binding sites of lower affinity or accessibility in chromatin. We showed previously that Gcn4 occupancies in WT cells are dictated primarily by their match to the consensus motif, with a secondary contribution from the distance of the motif from the dyad of the nearest nucleosome (Rawal et al. 2018b). Consistent with this, Gcn4 occupancies in WT_I cells are positively correlated with the match of the binding motif to the consensus sequence (quantified as “Find Individual Motif Occurrences” (FIMO) scores) for both 5’ and ORF peaks of Gcn4 occupancies (Fig. S5C-D). Accordingly, we used Gcn4 occupancies in WT_I cells as a proxy for affinity/accessibility of the binding motifs in chromatin. As shown above in Fig. 2C, the 5’ Gcn4 peaks in Set_1, showing the largest occupancy reductions in *snf2Δ PTET-STH1_I* cells have WT_I Gcn4 occupancies considerably greater than those in Set_2 and Set_3, which exhibit smaller reductions in binding in the double mutant (Fig. 2C(i) vs. (ii)-(iii), yellow). Consistent with this, the reductions in Gcn4 binding in the double mutant are positively correlated with WT_I Gcn4 occupancies for both the 5’ and ORF peaks (Fig. 2F(i)-(ii)). Moreover, the 5’ sites in Set_1 have higher median FIMO scores than those in Set_2 or _3 (Fig. S5E). The finding that Gcn4 sites of highest occupancy/affinity tend to show the largest occupancy decreases in the *snf2Δ PTET-STH1* mutant does not support the possibility that the reductions in Gcn4 abundance are important drivers of diminished Gcn4 occupancy in this mutant. The tendency for Gcn4 peaks with greater occupancies in WT to show larger binding reductions is also evident for the *ino80Δ* mutant, for both 5’ and ORF peaks (Fig. S4D(i)-(ii)). Because *ino80Δ* does not alter *GCN4* transcription (Fig. S5A) or Gcn4 protein abundance (Qiu et al. 2020), it appears that intrinsic properties of high-occupancy Gcn4 binding sites render them relatively more dependent on CR function for robust Gcn4 binding.

### Evidence that SWI/SNF, RSC, and Ino80C are recruited by Gcn4 to NDRs and act directly to regulate Gcn4 binding

To provide evidence that SWI/SNF, RSC and Ino80C function directly at Gcn4 motifs to regulate Gcn4 binding, we examined whether these CRs are recruited to Gcn4 binding sites. To this end, we conducted ChIP-seq analysis of the Myc epitope-tagged catalytic subunits, Snf2, Sth1, and Ino80, and corrected the occupancies for those obtained from ChIP-seq of an isogenic untagged strain. Heatmaps of the corrected CR occupancies sorted by the WT_I Gcn4 occupancies revealed that Snf2-myc occupancies are generally centered on the Gcn4 binding motifs, and that they tend to increase with increasing Gcn4 binding (Fig. 3A(i)), yielding a direct correlation between Snf2-myc_I and Gcn4_I occupancies (Fig. 3B(i)). A precise coincidence in the occupancies of Gcn4 and Snf2-myc is evident for several 5’ Gcn4 peaks associated with particularly high induced Snf2-myc occupancies (Fig. S6A-E). There is also evidence however that Snf2-myc occupancy is greatest upstream of the Gcn4 motif at a subset of 5’ sites (Fig. 3A(i)), which might indicate interaction of SWI/SNF with another transcription factor bound at these promoters in addition to Gcn4. The Sth1-myc occupancies are also generally centered on the Gcn4 motifs, but they are lower overall (Fig. 3A(ii)), and the correlation between Gcn4 and Sth1-myc occupancies is less pronounced (Fig. 3B(ii)); however, Snf2-myc and Sth1-myc occupancies at the 5’ Gcn4 peaks are positively correlated (Fig. S7A(i)). Ino80-myc occupancies also peak at, or somewhat upstream, of the Gcn4 motifs at most 5’ sites, and show a positive correlation with WT_I Gcn4 occupancies (Fig. 3B(iii)); although they are higher than expected at the 5’ sites of lowest Gcn4 occupancies found at the bottom of the heat map in Fig. 3A(iii). The Ino80-myc occupancies are additionally correlated with those for both Snf2-myc and Sth1-myc (Fig. S7A(ii)-(iii)). Importantly, the median occupancies of Snf2-myc, Sth1-myc, and Ino80-myc for all 5’ sites were significantly higher in cells induced by SM compared to uninduced cells (Fig. 3C(i)-(iii)). Together, these findings suggest that Gcn4 directly recruits SWI/SNF, Ino80C, and to a lesser degree RSC, to its 5’ binding sites.

**Figure 3.**
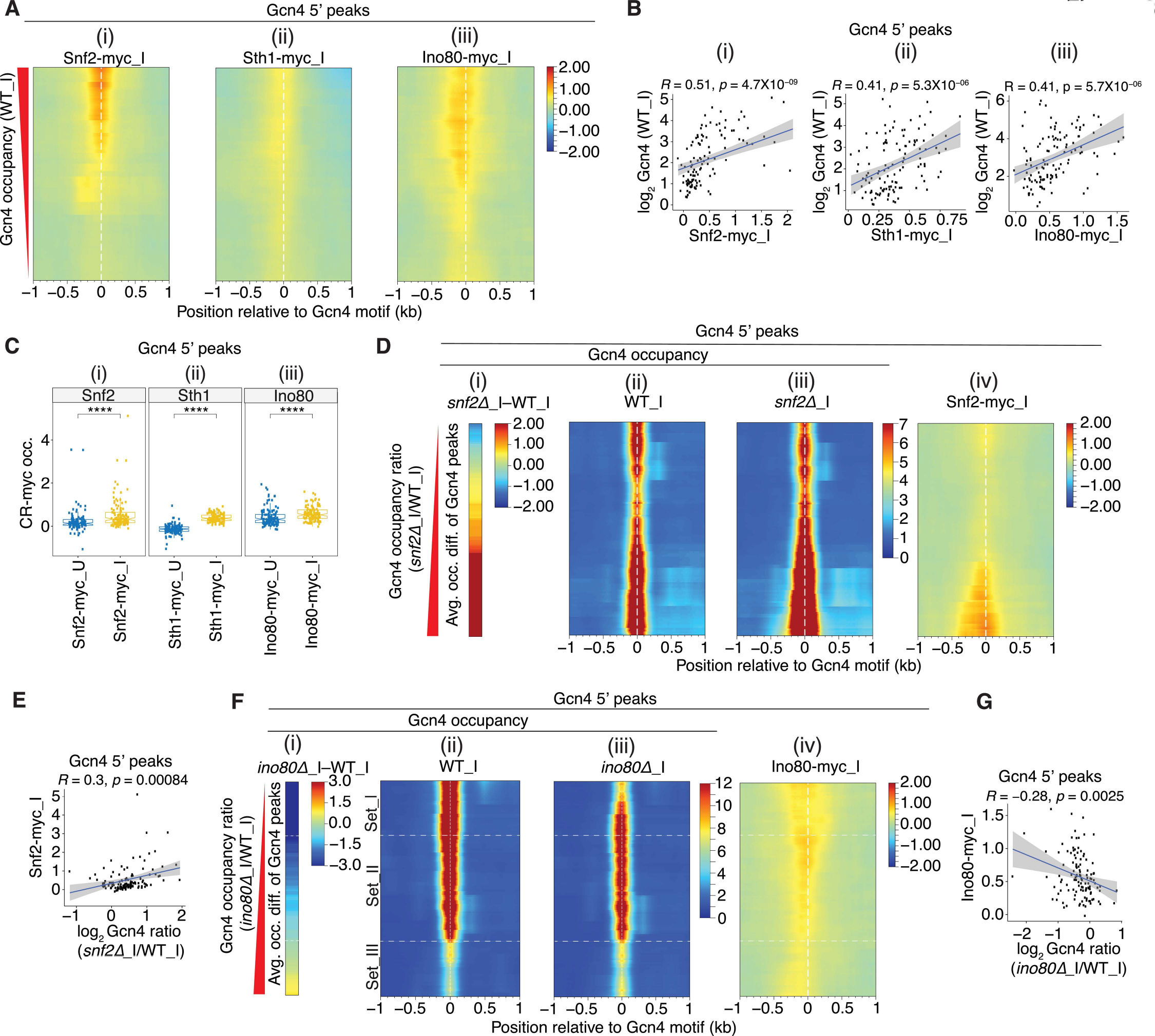
All three CRs are recruited near the Gcn4 motifs at 5’ sites. **(A)** Heat map depictions of corrected SM-induced occupancies for (i) Snf2-myc, (ii) Sth1-myc and (iii) Ino80-myc (corrected by subtracting the background myc-occupancies from untagged WT_I cells as described in Methods) at the 5’ Gcn4 peaks, sorted by decreasing Gcn4 occupancies in WT_I. Occupancies of the myc-tagged CR subunits were calculated from ChIP-seq data of mildly sonicated chromatin from 2 biological replicates for Snf2-myc_I and Sth1-myc_I, and 3 biological replicates for Ino80-myc_I . **(B)** Scatterplots of log2 Gcn4 occupancies in WT_I vs. corrected occupancies for (i) Snf2-myc_I, (ii) Sth1-myc_I, or (iii) Ino80-myc_I within ±100 bp windows surrounding the Gcn4 motifs of 5’ sites. Pearson correlation coefficients (*R*) and associated *p* values are indicated. **(C)** Notched box plots of (i) Snf2-myc, (ii) Sth1-myc, or (iii) Ino80-myc corrected occupancies per nucleotide within ±100 bp windows surrounding the Gcn4 motifs of 5’ sites in uninduced (_U) and SM induced (_I) conditions. **(D)** Heat maps depicting differences in Gcn4 occupancies between *snf211*_I and WT_I cells (i); Gcn4 occupancies surrounding the motifs of 5’ sites in (ii) WT_I or (iii) *snf211*_I cells; and (iv) Snf2-myc_I corrected occupancies surrounding the Gcn4 motifs of 5’ sites. Gcn4 5’ sites were sorted by increasing order of the ratio of Gcn4 occupancies in *snf211*_I vs. WT_I cells. **(E)** Scatterplot of corrected Snf2-myc occupancies within ±100 bp windows surrounding the Gcn4 motifs of 5’ sites vs. the log2 ratios of Gcn4 occupancies in *snf211_*I vs. WT_I cells at 5’ Gcn4 peaks. Pearson correlation coefficients (*R*) and associated *p* values are indicated. **(F)** Heat maps depicting differences in Gcn4 occupancies between *ino8011*_I and WT_I cells (i); Gcn4 occupancies surrounding the motifs of 5’ sites in (ii) WT_I or (iii) *ino8011*_I cells; and (iv) corrected Ino80-myc_I occupancies surrounding the Gcn4 motifs of 5’ sites. Gcn4 5’ sites were sorted by increasing order of the ratio of Gcn4 occupancies in *ino8011*_I vs. WT_I cells, and the first (Set_I, n=30), middle two (Set_II, n=57) and fourth (Set_III, n=30) quartiles of fold-changes are depicted. **(G)** Same analysis shown in (E) except for corrected Ino80-myc occupancies vs. the log2 ratios of Gcn4 occupancies in *ino8011_*I vs. WT_I cells.

Supporting this last conclusion, we showed previously that Gcn4 directly interacts with SWI/SNF and RSC in cells extracts in a manner dependent on key hydrophobic residues in the Gcn4 activation domain crucial for activation of Gcn4 target genes (Natarajan et al. 1999; Swanson et al. 2003), and that recruitment of both SWI/SNF and RSC to the *ARG1* UAS in SM-treated cells, as judged by ChIP analysis of Snf2-myc and Rsc8-myc, was diminished by deletion of *GCN4* (Swanson et al. 2003). Confirming our conclusion that Gcn4 also recruits Ino80C, we found here that Ino80-myc occupancies at the 5’ Gcn4 peaks were diminished by deletion of *GCN4* under both inducing and uninducing conditions (Fig. S7B-C). Together with our current findings that Snf2-myc, Sth1-myc, and Ino80-myc occupancies peak in the vicinity of the Gcn4 binding motifs at the majority of 5’ sites (Figs. 3A & S6), that occupancies of all three CRs are induced in parallel with Gcn4 induction by SM (Fig. 3C) and are positively correlated with Gcn4 occupancies for the group of 5’ genes (Fig. 3B), we consider it very likely that SWI/SNF, RSC, and Ino80C are all recruited directly by Gcn4 to the majority of 5’ genes.

We next sought evidence that SWI/SNF acts directly to dissociate Gcn4 from the 5’ binding sites where it negatively regulates Gcn4 binding. Sorting the 5’ motifs according to their increases in Gcn4 occupancy in *snf2Δ*_I versus WT_I cells (Fig. 3D (i)-(iii)), we found that Snf2-myc occupancies are generally higher at the 5’ motifs exhibiting the largest increases in Gcn4 occupancies conferred by the *snf2Δ* single mutation (and thus most negatively regulated by SWI/SNF), positioned at the bottom of the heat map in Fig. 3D (iv)); and there is a significant positive correlation between these two parameters for all 5’ sites (Fig. 3E). Interestingly, a small enrichment of Snf2-myc is also evident for the 5’ sites located at the top of the map in Fig. 3D(iv), containing the subset of sites where SWI/SNF stimulates rather than impedes Gcn4 binding in WT cells. Overall, these findings support the idea that in cells containing RSC, SWI/SNF directly regulates Gcn4 binding at 5’ sites, reducing binding at most sites but enhancing Gcn4 binding at a small subset of sites.

Carrying out the same analysis for Ino80C reveals that the 5’ sites showing the strongest reductions in Gcn4 binding conferred by *ino80Δ* (at the top of the heat maps in Fig. 3F(i)-(iii)) tend to have higher Ino80-myc occupancies (Fig. 3F(iv)), in this case producing a negative correlation between the two parameters (Fig. 3G). The correlation is likely weakened by the moderate enrichment for Ino80-myc at the bottom of the heat map for the 5’ sites that show moderately increased Gcn4 binding in *ino80Δ* cells (Fig. 3F(iv)). The elevated Ino80-myc occupancies at these latter sites could be explained by an inhibitory effect of Ino80C on Gcn4 binding, akin to our findings for SWI/SNF. Together, these results suggest that Ino80C also directly regulates Gcn4 binding at 5’ sites, functioning like RSC to enhance binding at most sites but mimicking SWI/SNF in reducing Gcn4 occupancy at a small subset of binding sites.

Our findings that RSC and Ino80C are recruited to 5’ sites on SM induction, and generally enhance Gcn4 occupancies, suggest a positive feedback loop; whereas the fact that SWI/SNF is highly recruited to 5’ sites where it generally reduces Gcn4 occupancies (in WT cells) indicates negative feedback control for this CR.

### Reduced Gcn4 occupancies are frequently associated with defective nucleosome eviction at Gcn4 binding motifs in mutants depleted of Sth1

The most likely mechanism for regulation of Gcn4 binding by CRs is the displacement of nucleosomes to either expose the motif and enhance binding or to occlude the motif and impede binding. ChIP-seq analysis of histone H3 in sonicated chromatin reveals reduced H3 occupancies within NDRs of genes harboring 5’ sites of Gcn4 occupancy on SM induction of WT cells (Fig. 4A, yellow vs. blue). These reductions in H3 seen in WT are diminished to similar extents in the *snf2Δ* and *PTET-STH1* single mutants, but to a greater extent in the *snf2Δ PTET-STH1* double mutant (Fig. 4A, grey, red, & cyan vs. yellow), indicating cooperation between RSC and SWI/SNF in evicting nucleosomes at 5’ Gcn4 sites. To examine the relationship between defects in H3 eviction and Gcn4 binding at particular 5’ sites, we first plotted the changes in H3 occupancies in *snf2Δ PTET-STH1*_I versus WT_I cells in the regions surrounding the 5’ Gcn4 motifs, sorted as above by their reductions in Gcn4 binding in this double mutant (Fig. 4B(i)-(ii)). The resulting difference map reveals an obvious tendency for the 5’ sites with the greatest reductions in Gcn4 binding, in Sets_1-2 (blue hues in Fig. 4B(i)), to also exhibit sizable increases in H3 occupancies at the Gcn4 motifs in the double mutant (orange hues in Fig. 4B(ii)), consistent with the idea that defective nucleosome eviction contributes to reduced Gcn4 binding in the absence of both RSC and SWI/SNF. Surprisingly however, 5’ sites at the bottom of the H3 difference map in Fig. 4B(ii) also show marked increases in H3 occupancies that are associated with increased, not decreased, Gcn4 binding in the double mutant (Set_3, orange hues in both Fig. 4B(i)-(ii)), a complexity we address below.

**Figure 4.**
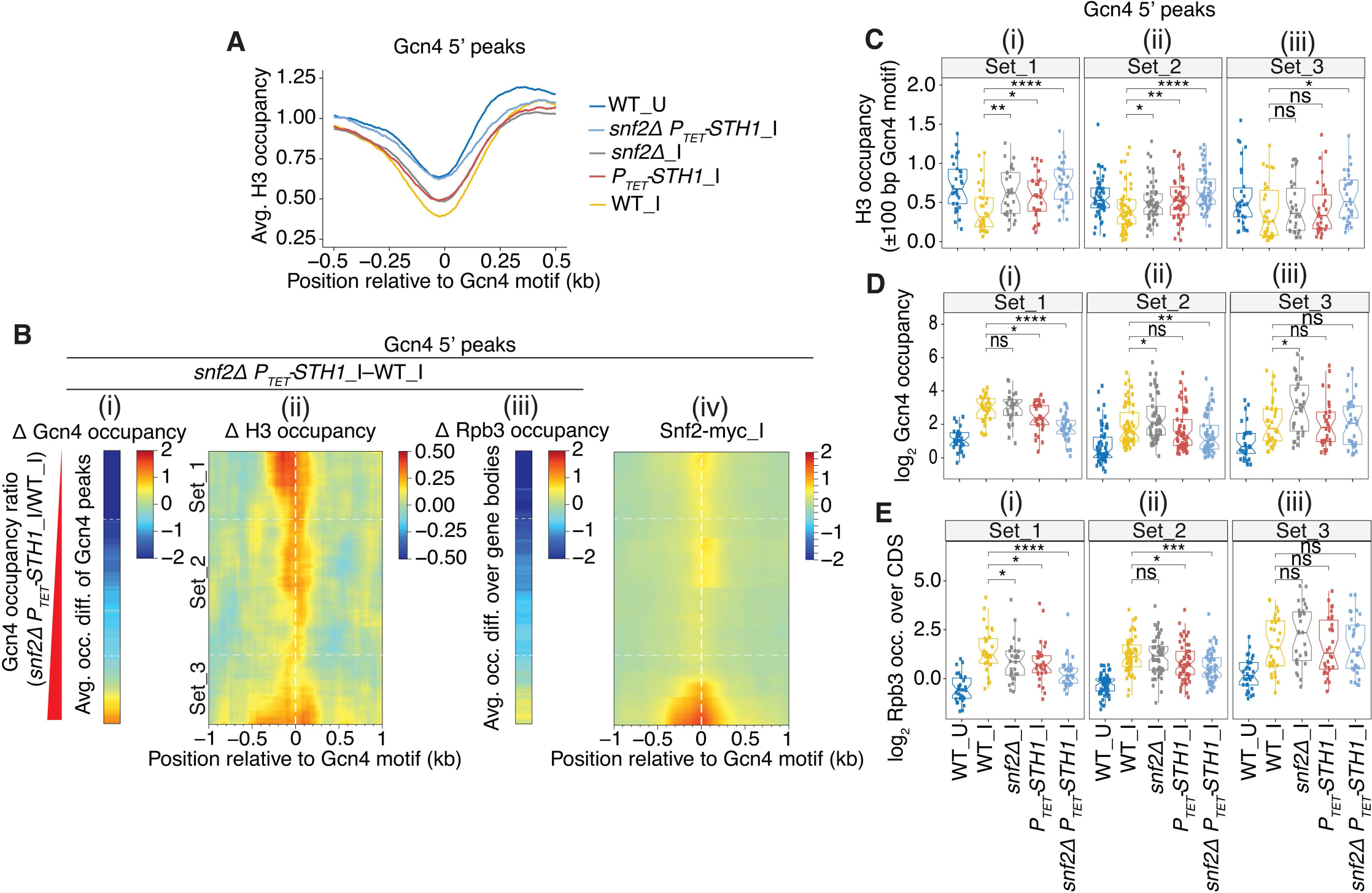
Defective eviction of nucleosomes surrounding 5’ Gcn4 motifs in SWI/SNF and RSC mutants. **(A)** Plots of H3 occupancies at each base pair surrounding the Gcn4 motifs averaged over all 5’ sites for the indicated strains/conditions, calculated from H3 ChIP-seq data. All of the average plots were found to converge at a position ∼1.0 kb upstream and downstream of the Gcn4 motifs. **(B)** (i)-(iii) Heat maps depicting differences between *snf211 PTET-STH1*_I and WT_I cells for (i) Gcn4 occupancies measured as in Fig 2A(i), (ii) H3 occupancies surrounding the Gcn4 motifs of 5’ sites from H3 ChIP-seq data, (iii) Rpb3 occupancies averaged over the CDS of 5’ genes, for Gcn4 5’ sites sorted by increasing order of fold-changes in Gcn4 occupancies in *snf211 PTET-STH1_*I vs. WT_I cells, as in Fig 2A-B. (iv) heat map of corrected Snf2-myc_I occupancies surrounding the Gcn4 motifs for the same order of 5’ sites as in (i)-(iii). **(C-E)** Notched box plots for the 3 sets of 5’ sites defined in Fig. 2A-B and indicated again in (B), depicting (C) H3 occupancies per base pair in the ±100 bp windows surrounding the Gcn4 motifs, (D) log2 Gcn4 occupancies taken from Fig 2C, and (E) log2 Rpb3 occupancies averaged over the CDS of genes with 5’ sites. H3 and Rpb3 occupancies were calculated from ChIP-seq data of sonicated chromatin from at least 3 biological replicates of WT_U, WT_I, or *snf211*_I, *PTET-STH1_*I and *snf211 PTET-STH1_*I cells.

The aforementioned trends in the heat maps were examined more closely by quantifying the changes in H3 occupancies for a 200bp window surrounding the Gcn4 motifs and comparing them to the corresponding changes in Gcn4 occupancies in the different mutants. In WT cells, SM-induction confers the expected reduced median H3 occupancies in parallel with increased Gcn4 occupancies for all three Sets of 5’ sites (cf. Figs. 4C & D, (i)-(iii), cols. 1-2). In the *PTET-STH1* and *snf2Δ PTET-STH1* mutants, the H3 occupancies are increased significantly for all three Sets (except Set_3 in the *PTET-STH1* single mutant), with larger increases in the double versus single mutant (Figs. 4C (i)-(iii), cols. 4-5 vs. 2); and the median Gcn4 occupancies are correspondingly reduced for Sets_1-2 in these two mutants (Figs. 4D (i)-(ii), cols. 4- 5 vs. 2). These results are illustrated differently in Fig. S8A by plotting changes in H3 against changes in Gcn4 occupancy in the double mutant vs. WT, revealing that nearly all of the 5’ sites in Sets_1-2 (red and green points) fall into the upper left-hand quadrant, exhibiting increased H3 and decreased Gcn4 occupancies in the double mutant. These results support our conclusion that RSC, and SWI/SNF upon Sth1 depletion, cooperate in evicting nucleosomes to facilitate Gcn4 binding at the 5’ sites in Sets_1-2. The association between reduced Gcn4 binding and increased H3 occupancies at such Gcn4 binding sites conferred by *PTET-STH1* in the single and double mutants is evident at the archetypal Set_1 and Set_2 genes depicted in Fig. S2A-B.

For the Set_3 sites, by contrast, increased H3 occupancies are generally not associated with decreased Gcn4 binding in the double mutant (cf. Figs. 4C(iii) and 4D(iii), col. 5 vs. 2); and Gcn4 occupancy is even increased for the subset of sites at the very bottom of the heat map (Fig. 4B(i)) despite their increased H3 occupancies in the double mutant (Fig. 4B(ii)). This uncoupling of Gcn4 and H3 occupancies for the Set_3 sites is also evident in the scatterplot of Fig. S8A, as a fraction of the Set_3 sites (blue points) fall into the upper right-hand quadrant indicating increased occupancies for both H3 and Gcn4 in the *snf2Δ PTET-STH1* double mutant vs. WT. This uncoupling is particularly noteworthy in both *PTET-STH1* single and double mutants for the Set_3 archetype *ARG1* shown in Fig. S2C. We concluded above that SWI/SNF exerts offsetting stimulatory and inhibitory effects on Gcn4 binding at the Set_3 sites in the double mutant. Hence, to account for the uncoupling of changes in H3 and Gcn4 occupancies at Set_3 sites, we suggest that elimination of SWI/SNF in the double mutant confers reduced nucleosome eviction at these sites, but the resulting increased H3 occupancies do not reduce Gcn4 binding owing to concurrent loss of a distinct SWI/SNF function that inhibits Gcn4 binding, which we speculate below might involve nucleosome sliding. Consistent with the proposed positive role of SWI/SNF in nucleosome eviction for the fraction of Set_3 sites at the very bottom of the heat-map in Fig. 4B(ii), these sites also show high-level SWI/SNF recruitment (Fig. 4B(iv)).

Uncoupling of changes in H3 and Gcn4 occupancy is also observed for the majority of 5’ sites in the *snf2Δ* single mutant. First, examining the scatterplot of H3 versus Gcn4 occupancy changes in Fig. S8B shows that *snf2Δ* increases the H3 occupancies for a large proportion of all 5’ sites, with the majority of points mapping above the x-axis, but most points fall into the upper right-hand quadrant indicating increased Gcn4 binding conferred by *snf2Δ*. Second, the *snf2Δ* mutation leads to increased median H3 occupancies for Sets_1-2 (Figs. 4C (i)-(ii), col. 3 vs. 2), consistent with nucleosome eviction by SWI/SNF; but *snf2Δ* has either no effect (Set_1) or confers increased rather than decreased median Gcn4 binding (Set_2) at these sites (Figs. 4D (i)-(ii), col. 3 vs. 2). For the sites in Set_3, the *snf2Δ* single mutation has no effect on median H3 occupancies and confers the largest increase in median Gcn4 binding observed among all three Sets (cf. Figs. 4C(iii) and 4D(iii), col. 3 vs. 2). Thus, the effect of *snf2Δ* in elevating Gcn4 occupancies increases progressively from Set_1 to Set_3 (Fig. 4D, (i)-(iii), cols. 3 vs. 2) and is associated with a progressively decreasing elevation of nucleosome occupancies across these sites (Fig. 4D, (i)- (iii), cols. 3 vs. 2). This trend is also illustrated in Fig. S9A where the only sites showing decreased Gcn4 binding in *snf2Δ* cells, located at the top of the heat-map in panel (i), also show a marked increase in H3 occupancies at these sites in panel (ii). Increased H3 occupancy that is unassociated with decreased Gcn4 binding in the *snf2Δ* single mutant is observed for the Set_3 archetypal gene *ARG1* in Fig. S2C, and is especially pronounced for the Set_1 archetype *HIS4* (Fig. S2A). The finding that increased H3 occupancies is frequently associated with increased rather than decreased Gcn4 binding in *snf2Δ* cells can be explained, as suggested above, if the reduced nucleosome eviction conferred by *snf2Δ* is offset by loss of the proposed SWI/SNF function that impedes Gcn4 binding.

Finally, to evaluate more completely the minority fraction of 5’ sites where SWI/SNF stimulates rather than impedes Gcn4 binding in WT cells, we sorted all of the 5’ sites by their reductions in Gcn4 occupancies in *snf2Δ* versus WT cells, placing those sites at the top of heat map in Fig. S9B(i). Importantly, these sites also exhibit marked increases in H3 occupancies at the Gcn4 motifs (Fig. S9B(ii) vs. (i)). Hence, at this particular subset of 5’ sites, SWI/SNF plays only the conventional role observed for RSC of evicting nucleosomes to promote Gcn4 binding.

As described fully in Supplementary Figure S10, analysis of the *ino80Δ* mutant provides evidence that Ino80C enhances Gcn4 binding at a subset of 5’ sites in Set_I by evicting nucleosomes surrounding the relevant binding motifs, which is highly similar to our findings above for the *PTET-STH1* single mutant. The association between reduced Gcn4 binding and increased H3 occupancies at the Gcn4 binding sites conferred by *ino80Δ* is illustrated for *YHR162W* and *HIS4* in Fig. S10E.

In summary, the reduced Gcn4 binding conferred by the mutations that impair RSC or Ino80C can be attributed to the increased H3 occupancies found at the binding sites in these mutants, whereas the effects of *snf2Δ* are more complex at the 5’ sites owing to the opposing roles that SWI/SNF can play in (i) stimulating Gcn4 binding via nucleosome eviction or (ii) impeding Gcn4 binding by a mechanism unrelated to nucleosome eviction. An intriguing possibility is that the latter inhibitory function of SWI/SNF might involve sliding of the remaining, non-evicted nucleosomes over the Gcn4 binding motifs, in a manner demonstrated previously for yeast SWI/SNF and the Gal4 DNA binding domain in vitro (Li et al. 2015).

### A combination of impaired Gcn4 binding and nucleosome eviction is associated with reduced transcription in the CR mutants

We asked next whether the reductions in Gcn4 binding at 5’ sites in the *PTET-STH1* and *snf2Δ PTET-STH1* mutants confer reduced transcription of the associated genes. Indeed, for genes in Set_1 and Set_2, decreased Gcn4 binding is generally associated with reduced Rpb3 occupancies averaged over the downstream CDS in *snf2Δ PTET-STH1*_I versus WT_I cells. This trend is evident in both the heat-maps of Fig. 4B (panel (iii) vs. (i), Sets_1-2) and in box-plot comparisons of median occupancies of Rpb3 (Fig. 4E(i)-(ii), col. 5 vs. 2) versus Gcn4 (Fig. 4D(i)-(ii), col. 5 vs. 2) for the Set_1 and Set_2 genes. A similar association between reduced Rpb3 and diminished Gcn4 occupancies was found for the Set_1 and Set_2 genes in the *PTET-STH1* single mutant (Fig. 4E(i)-(ii), col. 4 vs. 2) & Fig. S9C(iii) vs. (i)). Note however that these genes also exhibit increased promoter H3 occupancies in the *snf2Δ PTET-STH1* and *PTET-STH1* mutants (Fig. 4C(i)-(ii) (cols. 4-5 vs. 2), which might contribute to their reduced transcription owing to impaired PIC assembly.

The genes in Set_3 exhibit increased promoter H3 levels in the double mutant (Fig. 4C(iii), col. 5 vs. 2), which appear to be insufficient to impair transcription (Fig. 4E(iii), col. 5 vs. 2). Given the nearly WT Gcn4 occupancies of the Set_3 genes in this mutant (Fig. 4D(iii), col. 5 vs. 2), one possibility is that a reduction in Gcn4 binding is required in addition to defective nucleosome eviction for impaired transcription, as observed above for genes in Sets_1 and 2 in the double mutant, perhaps owing to diminished recruitment of other coactivators by Gcn4 at its reduced occupancies. This inference can also account for the finding that the *snf2Δ* single mutation reduces transcription of only the Set_1 genes at the top of the difference heat maps in Figs. S9A(i)-(iii), which exhibit both reduced Gcn4 binding and marked increases in promoter H3 occupancies. Analysis of the *ino80Δ* mutant by the same approaches, described fully in Fig. S10, likewise provides evidence that a combination of defects in Gcn4 binding and impaired promoter nucleosome eviction confer reduced transcription of a subset of genes with 5’ Gcn4 binding sites by *ino80Δ*.

### Evidence that the CRs stimulate TBP recruitment at SM-induced genes by evicting or displacing the +1 nucleosome

We examined next the contributions of SWI/SNF and RSC to PIC assembly at genes with 5’ Gcn4 binding sites by conducting ChIP-seq of native TBP in the same CR mutants described above. The averaged occupancy plots revealed strong induction of TBP binding on SM treatment of WT cells, peaking ∼130 bp downstream of the 5’ motifs (Fig. 5A(i), yellow vs. blue). Lower-level TBP binding also was induced ∼200 bp upstream of the motifs, consistent with activation of bidirectional promoters by Gcn4. Essentially identical results were obtained in WT cells treated with Dox prior to SM treatment (Fig. 5A(ii), black vs. blue), which served as the WT control for the *PTET-STH1,* and *snf2Δ PTET-STH1* mutants. The averaged TBP occupancies at both locations were substantially reduced in the SM-treated *snf2Δ PTET-STH1* double mutant and *ino80Δ* strain (Fig. 5A(ii), cyan vs. black; Fig. 5A(i), gold vs. yellow), but were either unaffected or slightly increased, respectively, in the *PTET-STH1* and *snf2Δ* single mutants (Fig. 5A(ii), red vs. black; Fig. 5A(i), grey vs. yellow). Interestingly, while not reducing TBP occupancies, the *snf2Δ* mutation shifted the position of TBP binding upstream towards the Gcn4 binding site (Fig. 5A(i) and 5C(ii)). The latter might indicate that TBP is recruited by Gcn4 to the UAS but is not efficiently delivered to the core promoter downstream, but additional work is required to understand the underlying mechanism. For Gcn4 motifs located within ORFs, induction of TBP occupancy peaks in WT cells was again observed both upstream and downstream of the motifs (Fig. S11A-B), consistent with our identification of bidirectional antisense and subgenic sense transcripts induced by internal Gcn4 binding (Rawal et al. 2018b). As above, the averaged TBP occupancies within ORFs was markedly reduced only in the *snf2Δ PTET-STH1* double mutant and the *ino80Δ* strain (Fig. S11A-B).

**Figure 5.**
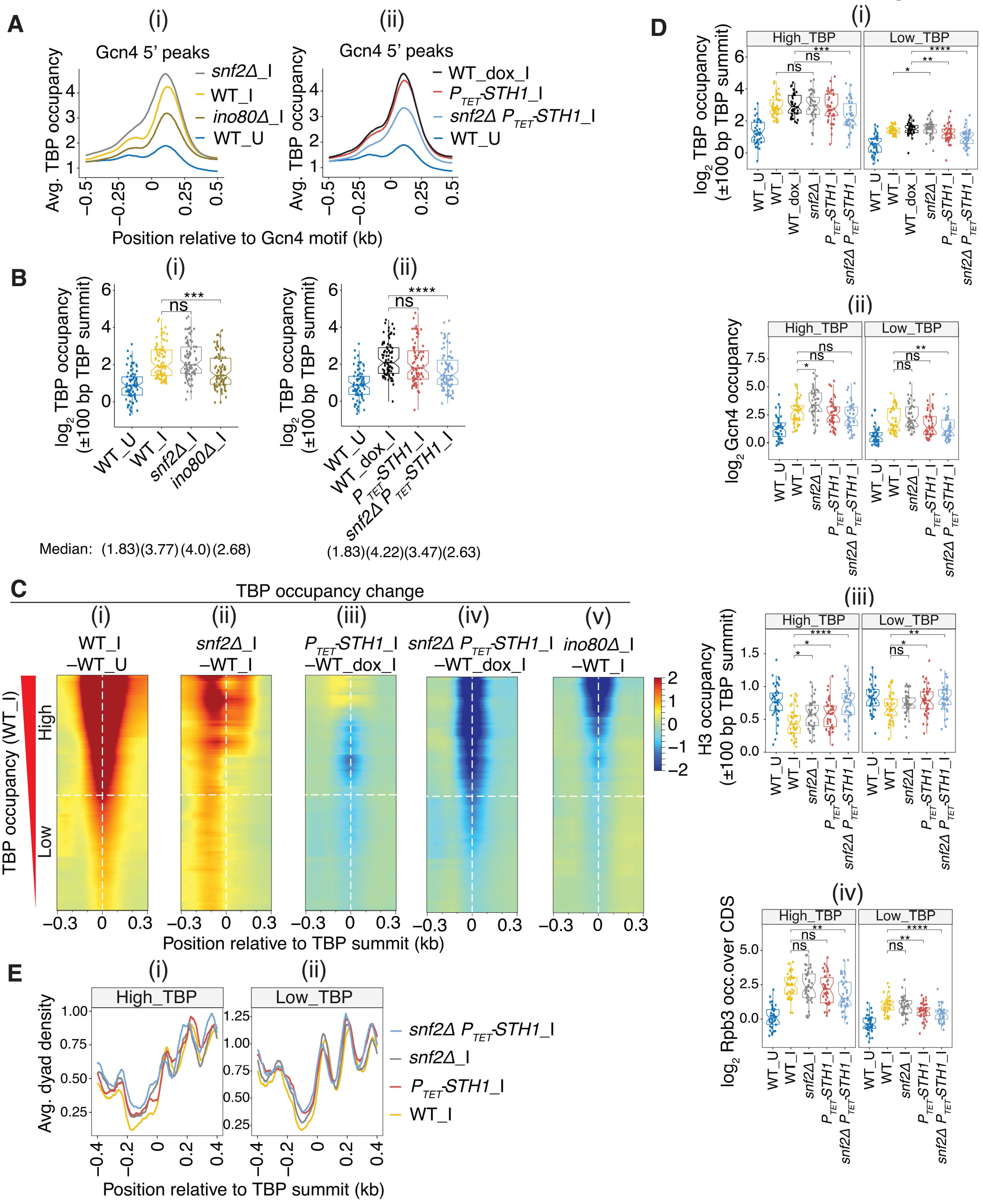
Reduced TBP recruitment in SWI/SNF and RSC mutants is frequently associated with increased +1 nucleosome occupancies in promoters of genes with 5’ Gcn4 binding sites. (A) (i)-(ii) Averaged TBP occupancies surrounding Gcn4 5’ motifs from ChIP-seq data of sonicated chromatin from at least 2 biological replicates of the indicated strains, with labels along the plots are arranged in decreasing order of summit heights. (B) (i)-(ii) Notched box plots of log2 TBP occupancies measured within ±100 bp of TBP peak summits identified by MACS2 peak analysis in the promoters of 83 of the 5’ Gcn4 target genes in the indicated yeast strains, color-coded as in (A). (C) Heat map depictions of differences in TBP occupancies surrounding TBP peak summits in the 83 5’ genes from (B) sorted in decreasing order of WT_I TBP levels in (i) WT_I vs. WT_U, (ii) *snf211*_I vs. WT_I, (iii) *PTET-STH1_*I vs. WT_I*, (iv) snf2Δ PTET-STH1_*I vs. WT_I, and (v) *ino8011*_I vs. WT_I, color-coded as indicated to the right. Two equal bins based on WT_I TBP of High_TBP or Low_TBP occupancies are indicated. (D) Notched box plots of factor occupancies in the 83 5’ gene promoters for the two bins defined in (C) for (i) log2 TBP within ±100 bp of TBP peak summits, (ii) log2 Gcn4 at the respective Gcn4 peaks, (iii) H3 within ±100 bp of the TBP peak summits, and (iv) log2 Rpb3 within the CDS of target genes. (E) Average dyad density from H3 MNase-ChIP-seq data aligned to the TBP summits of the 83 5’ genes binned as in (C). Midpoints (dyads) of nucleosomal size sequences between 125 and 175 bp were averaged and plotted relative to the TBP summits. Profiles were smoothed using a moving average filter with a span of 31 bp. Yeast strain labels are arranged in decreasing order of dyad densities at the +1 nucleosome spanning the TBP summits.

To quantify the effects of the CR mutations on TBP recruitment at individual genes with 5’ Gcn4 peaks, we determined the TBP occupancies in a window of ±100 bp surrounding the summits of each TBP peak that we identified at 83 of the 117 genes with 5’ Gcn4 sites. Consistent with the averaged TBP data in Fig. 5A, the results revealed significant reductions in median TBP occupancies only in the *snf2Δ PTET- STH1* and *ino80Δ* mutants compared to WT_I cells (Fig. 5B(i)-(ii)). We also displayed the TBP occupancy changes gene-by-gene using TBP difference heat maps in which the 5’ genes were sorted by their TBP occupancies in WT_I cells. As might be expected, the genes with highest TBP occupancies in WT_I cells also display the largest inductions of TBP binding on SM-treatment of WT cells (Fig. 5C(i), High vs. Low group). Most of the genes in the High_TBP group (upper half of each map) show reductions in TBP binding in both the *snf2Δ PTET-STH1* and *ino80Δ* mutants compared to WT induced cells (Fig. 5C(iv)-(v)); whereas only about one-half of these genes exhibit even moderately reduced TBP binding in the *PTET-STH1* single mutant (Fig. 5C(iii)), and none show reduced TBP binding in *snf2Δ*_I vs. WT_I cells (Fig. 5C(ii)). Together, the results in Fig. 5A-C indicate overlapping functions of RSC and SWI/SNF in promoting TBP recruitment on Gcn4 binding at 5’ sites, wherein a strong recruitment defect occurs only when both Sth1 and Snf2 are removed simultaneously in the *snf2Δ PTET-STH1* double mutant. In contrast, eliminating Ino80 confers a marked reduction in TBP recruitment in otherwise WT cells at most of the genes exhibiting the highest levels of TBP recruitment by Gcn4 in WT cells.

Recently, we provided evidence that reductions in Pol II occupancies in *ino80Δ* cells are associated with both decreased TBP occupancies and increased promoter H3 occupancies at the SM-induced genes, consistent with the idea that Ino80C promotes Gcn4-activated PIC assembly and transcription by evicting promoter nucleosomes (Qiu et al. 2020). To extend this model to include RSC and SWI/SNF, we quantified the H3 occupancies surrounding the summits of the TBP peaks for the High_TBP and Low_TBP groups of 5’ genes defined above. As expected, these H3 occupancies are reduced on SM-treatment of WT cells (Fig. 5D(iii), High_ & Low_TBP, cols 1-2), consistent with eviction of promoter nucleosomes to facilitate TBP binding. Importantly, we observed increased H3 occupancies in the *snf2Δ PTET-STH1* double mutant that are greater in magnitude than observed in the *snf2Δ* or *PTET-STH1* single mutants under inducing conditions, particularly for the High_TBP group (Fig. 5D(iii), High_ & Low_TBP, cols 3-5 vs. 2). We also examined the positions of nucleosome dyads measured by H3 MNase ChIP-Seq analysis, which revealed a shift upstream in the averaged dyad densities of +1 nucleosomes that results in greater dyad densities immediately 5’ of the TBP summits in the mutants, which was most pronounced in the double mutant for the High_TBP group (Fig. 5E(i)-(ii), cyan vs. grey & red vs. yellow). The fact that combining the *snf2Δ* and *PTET-STH1* mutations in the double mutant confers additive increases in nucleosome occupancies at TBP binding sites that are coupled with additive reductions in TBP recruitment at many of these sites (noted above in Fig. 5D(i), cols. 6 vs. 4-5) supports the model that RSC and SWI/SNF functionally cooperate to enhance Gcn4-activated TBP binding by eviction and repositioning of +1 nucleosomes. The coupling of reduced TBP binding and elevated H3 occupancies in the *snf2Δ PTET-STH1* double mutant is illustrated for six archetypal genes with 5’ Gcn4 sites in Fig. S12A-B(i)-(iii). The unexpected increase in TBP recruitment in the *snf2Δ* strain evident in Fig. 5C(ii) might arise from the elevated Gcn4 binding found at most 5’ sites in this mutant (Fig 5D(ii), cols. 3 vs. 2), which could increase recruitment of the cofactors serving as TBP adaptors, SAGA and TFIID, to elevate TBP recruitment despite somewhat higher +1 nucleosome occupancies.

Consistent with our previous findings (Qiu et al. 2020), reduced TBP recruitment conferred by *ino80Δ* at genes with 5’ Gcn4 sites also is frequently associated with elevated H3 occupancies at the TBP binding sites (Fig. S11C (i)-(ii)), as illustrated for four archetypal genes in Fig. S11D-E.

Finally, as might be expected, the Pol II (Rpb3) occupancies in WT_I cells are greater for the High_TBP versus Low_TBP group of genes containing 5’ Gcn4 sites (Fig. 5D(iv), High_ vs. Low_, col. 2). Importantly, we observed marked reductions in Rpb3 occupancies at these genes only in the *snf2Δ PTET- STH1* double mutant, with smaller reductions in the *PTET-STH1* single mutant and no significant changes in the *snf2Δ* single mutant (Fig. 5D(iv), cols. 3-5 vs. 2). The fact that the changes in Poll II occupancies generally parallel the changes in TBP occupancies in these mutants (noted above in Fig. 5D(i) & Fig. 5C(ii)- (iv)) supports the model that RSC and SWI/SNF functionally cooperate to enhance Gcn4-activated transcription of these genes by stimulating PIC assembly.

### SWI/SNF functions with RSC and Ino80C in TBP recruitment only at highly expressed genes in non- stressed cells

Having found that SWI/SNF, RSC, and Ino80C all participate in TBP recruitment at genes with 5’ Gcn4 binding sites in starved cells (Figs. 5A-B), we next examined their relative importance for this step of PIC assembly at genes expressed at high levels in non-starved cells (ie. not treated with SM). In the absence of stress, ∼50% of Pol II and the transcriptional machinery is devoted to transcription of ribosomal protein genes (RPGs) (Kuang et al. 2014). Accordingly, we examined the RPGs as a group and divided the remaining constitutively expressed genes into ten deciles according to their Rpb3 occupancies. Interestingly, all four of the CR mutations conferred significantly decreased TBP occupancies at the RPGs, with the greatest reductions in the *snf2Δ PTET-STH1* double mutant (Fig. 6A), comparable marked reductions in the *PTET-STH1* and *ino80Δ* single mutants, and somewhat smaller reductions in the *snf2Δ* single mutant (Fig. 6A). These findings indicate functional cooperation between RSC and SWI/SNF, and a substantial contribution from Ino80C, in TBP recruitment at the RPGs. All of the mutants also displayed reduced Rpb3 occupancies at the RPGs, but here *snf2Δ* conferred reductions comparable to those observed in the *snf2Δ PTET-STH1* double mutant and the *ino80Δ* mutant, with lesser reductions in the *PTET- STH1* single mutant (Fig. 6B). These last results might indicate that SWI/SNF makes an additional contribution to transcription beyond TBP recruitment at these genes.

**Figure 6.**
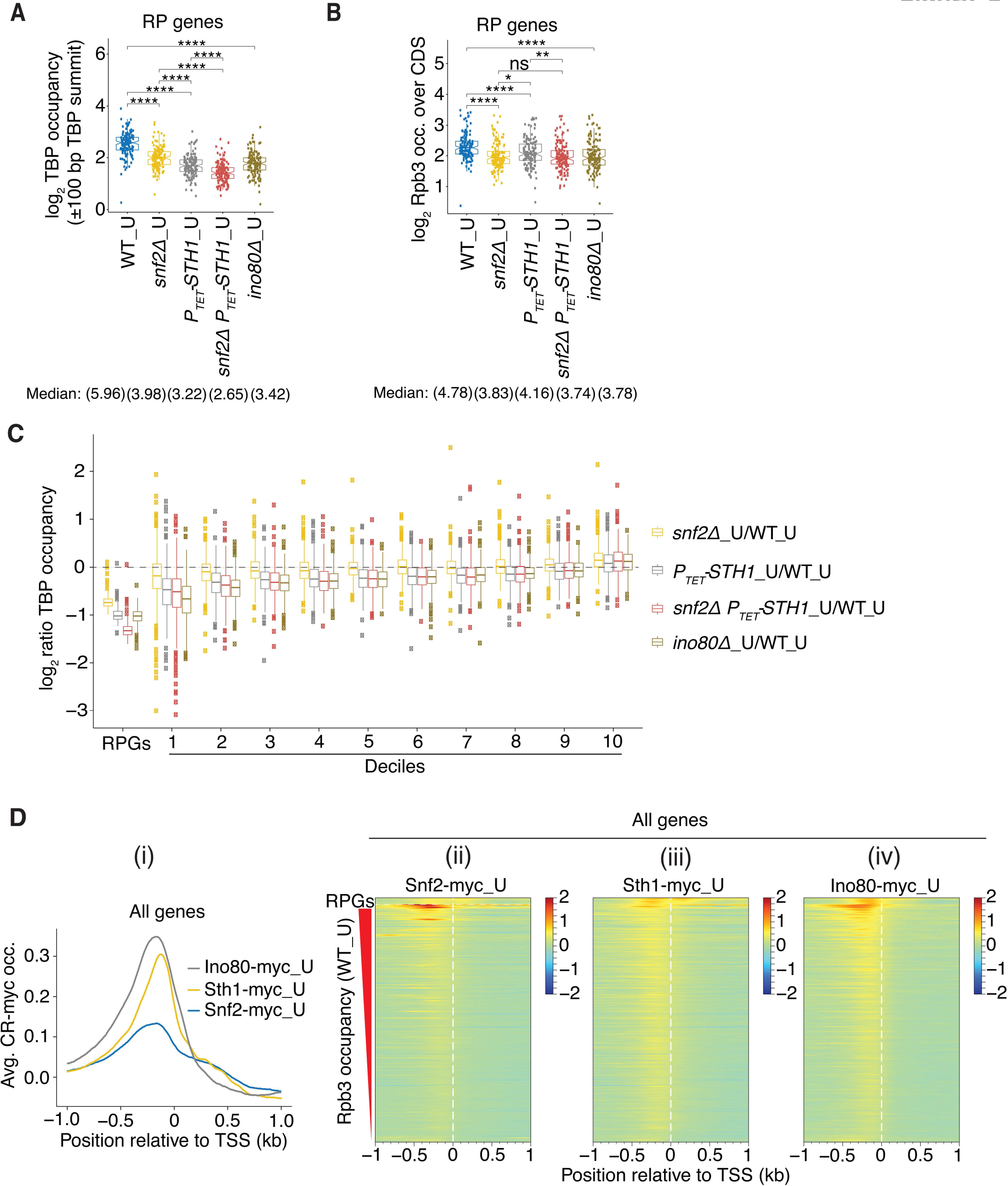
RSC and Ino80C act broadly whereas SWI/SNF functions mainly at highly expressed genes to promote TBP recruitment. **(A-B)** Notched box pots of log2 TBP occupancies within ±100 bp of TBP peak summits (A) and log2 CDS Rpb3 occupancies (B) at all RP genes in WT or the indicated mutants, in uninduced cells. **(C)** Notched box plots of log2 ratios of TBP occupancies in the indicated mutants vs. WT in uninduced cells for RP genes and Deciles 1-10 of all genes arranged in descending order of Rpb3 occupancies in WT_U cells. **(D)** (i) Averaged corrected Ino80-myc_U, Snf2-myc_U, and Sth1-myc_U occupancies aligned to the TSSs of all genes. (ii)-(iv) Heat map depictions of corrected occupancies of (ii) Snf2-myc_U, (iii) Sth1-myc_U, and Ino80-myc_U (iv) at RP genes (at the top) and all other genes sorted in decreasing order of WT_U Rpb3 levels, color-coded as indicated on the right.

Examining changes in TBP occupancies at the remaining genes reveals reduced TBP recruitment for the ∼40% most highly expressed genes in Deciles 1-4 in all four CR mutants, albeit of smaller magnitude compared to the RPGs (Fig. 6C). By comparison to the RPGs, SWI/SNF makes a smaller contribution than RSC for the genes in Deciles 1-4, such that reductions in TBP occupancies are smaller in the *snf2Δ* versus *PTET-STH1* single mutant, and similar in magnitude between the *PTET-STH1* single mutant and *snf2Δ PTET- STH1* double mutant (Fig. 6C, cols. 1-3 for Deciles 1-4). The effect of *ino80Δ* is relatively greater in comparison to the other mutants for Deciles 1-4 compared to that observed for the RPGs, being comparable to the *snf2Δ PTET-STH1* double mutant in reducing TBP recruitment for the former genes (Fig. 6C, Deciles 1-4, cols. 3-4). For most of the remaining genes in Deciles 5-9, it appears that RSC and Ino80C make comparable, modest contributions to TBP recruitment with little involvement of SWI/SNF, as we observe similar reductions in TBP occupancies in the *PTET-STH1, snf2Δ PTET-STH1,* and *ino80Δ* mutants, but no significant change in *snf2Δ* cells (Fig. 6C, Deciles 5-9, cols. 1-4). All of the mutants show somewhat elevated TBP occupancies for Decile 10 (Fig. 6C, Decile 10, cols. 1-4), which might result from reduced competition with the more highly expressed genes whose ability to recruit TBP is impaired in the CR mutants. Overall, the results indicate that RSC and Ino80C function broadly to enhance TBP recruitment at constitutive genes regardless of gene expression levels, whereas SWI/SNF acts primarily at highly expressed genes, and particularly at the RPGs, to help stimulate this step of PIC assembly.

Examining averaged occupancies of Snf2-myc, Sth1-myc, and Ino80-myc at all genes in uninduced cells reveals enrichment of all three factors upstream of the TSS, with relatively higher averaged occupancies for Ino80-myc and Sth1-myc versus Snf2-myc (Fig. 6D(i)). Interrogating individual genes using heat maps, sorting genes by Rpb3_I occupancies, shows that Sth1-myc and Ino80-myc are moderately enriched upstream of the TSS at nearly all genes, whereas Snf2-myc is more restricted to highly expressed genes near the top of the map (Fig. 6D(ii)-(iv)). Thus, the relative occupancies of RSC, SWI/SNF, and Ino80C generally parallel their relative importance for TBP recruitment at these gene groups, which is consistent with direct contributions of the three CRs to this step of PIC assembly throughout the genome.

Finally, we compared the dependence on the three remodelers for TBP recruitment between genes that rely primarily on TFIID or utilize SAGA in addition to TFIID for this step of PIC assembly. The 655 genes designated as coactivator-redundant, which exhibit overlapping functions for SAGA or TFIID (Donczew et al. 2020), show greater reductions in TBP in the *PTET-STH1, snf2Δ PTET-STH1,* and *ino80Δ* mutants than do the 4245 genes that rely primarily on TFIID (Fig. S13). Consistent with this, the most highly transcribed decile of the 4680 non-RPG genes (Decile 1), shown above to exhibit the greatest TBP reductions in all four remodeler mutants among all deciles (Fig. 6C), are highly enriched for coactivator-redundant genes (53%) compared to Deciles 2-10 (5-19% coactivator-redundant genes) and the non-RPG genes as a group (∼14%). Moreover, the genes with 5’ Gcn4 sites, which show redundant contributions by SWI/SNF and RSC and a marked requirement for Ino80C for TBP recruitment (Fig. 5B(i)-(ii)), are comprised of 45% coactivator-redundant genes. These findings suggest a tendency for coactivator-redundant genes to show a greater requirement than TFIID-dependent genes for RSC and Ino80C, and also for SWI/SNF among the most highly expressed members, for robust TBP recruitment. Interestingly, the RPGs are dramatic exceptions to this tendency, as they exhibit the strongest reductions in TBP recruitment in all four CR mutants of any gene group we analyzed, but are almost exclusively (95%) TFIID-dependent genes. Thus, while showing minimal dependence on SAGA, the high-level transcription achieved by the RPGs involves the concerted functions of all three CRs for efficient TBP recruitment.

## DISCUSSION

In this report, we have provided evidence for distinct contributions of three CRs in regulating the binding of transcriptional activator Gcn4 at its numerous target genes, and illuminated functions of these CRs in controlling TBP recruitment for PIC assembly and transcription, both at Gcn4 target genes and more generally throughout the genome. Gcn4 binds in the NDRs of most of its target genes (5’ sites), but within the CDS at a subset of activated genes (ORF sites) where it stimulates bidirectional transcription of antisense and subgenic sense transcripts as well as activating the 5’-positioned promoters (Rawal et al. 2018b). We found that depleting or eliminating the catalytic subunits of RSC (Sth1) or Ino80C (Ino80) reduces Gcn4 occupancies at both 5’ and ORF binding sites; with differential contributions of these two CRs at different 5’ sites. Surprisingly, SWI/SNF functions oppositely at 5’ sites in WT cells, generally reducing their Gcn4 occupancies, while having little effect at the ORF sites. In cells depleted of Sth1, by contrast, SWI/SNF functionally substitutes for RSC in promoting Gcn4 binding at both 5’ and ORF sites, as eliminating Snf2 exacerbates the effect of depleting Sth1 in reducing Gcn4 occupancies at the sites most affected by depleting Sth1 alone from otherwise WT cells. These opposing effects of SWI/SNF appear to cancel out at the subset of 5’ sites where SWI/SNF most strongly impedes Gcn4 binding in WT cells, such that the double mutation *snf2Δ PTET-STH1* has little effect on their Gcn4 occupancies. Ino80C also stimulates Gcn4 binding at 5’ sites that are highly dependent on RSC, but it stimulates binding at other 5’ sites that are largely independent of RSC, and where SWI/SNF impedes rather than enhances Gcn4 binding. Thus, our results have uncovered distinct contributions of these three CRs to Gcn4 binding at different 5’ sites.

The Gcn4 binding motifs of 5’ sites tend to reside in the center of NDRs, while the ORF motifs are generally located in linkers connecting genic nucleosomes, suggesting that Gcn4 binding is impeded by inclusion of its recognition sequence within nucleosomes (Rawal et al. 2018b). Consistent with this, our results suggest that RSC and Ino80C enhance Gcn4 binding by removing nucleosomes from the binding motifs in NDRs. Thus, the subset of 5’ sites showing decreased Gcn4 occupancies in the *PTET-STH1* single mutant or in the *ino80Δ* strain also tend to exhibit defective H3 eviction at these 5’ sites in those mutants (Fig. 4C-D(i) & Fig. S10B-C(i)). We observed similar inverse relationships between effects of the *snf2Δ PTET- STH1* double mutation on H3 and Gcn4 occupancies for the subsets of 5’ sites showing the largest reductions in Gcn4 occupancies in this double mutant (Sets_1-2). This correlation was not evident however for the 5’ sites that are least impaired for Gcn4 binding in the double mutant (Set_3), where increased H3 occupancies were not accompanied by appreciable reductions in Gcn4 binding (Figs. 4C-D, (iii) vs. (i)-(ii)), which we attributed to eliminating the inhibitory effect of SWI/SNF on Gcn4 binding at these sites by the *snf2Δ* mutation in the double mutant. The dual, opposing effects of SWI/SNF on Gcn4 binding can also account for the fact that the 5’ sites in Sets_1-2 show increased H3 occupancies in the *snf2Δ* single mutant but generally no reduction in Gcn4 binding (Figs. 4C-D(i)-(ii)); and also that the sites in Set_3 exhibiting the least effect of *snf2Δ* on H3 eviction show markedly increased Gcn4 binding in this mutant (Figs. 4C-D(iii)). Thus, the effect of *snf2Δ* on Gcn4 binding at any given site appears to reflect the relative importance of SWI/SNF in evicting nucleosomes (to enhance binding) versus its opposing role in diminishing Gcn4 binding.

The negative effect of SWI/SNF on Gcn4 binding in WT cells is not unprecedented. In vitro studies indicate that SWI/SNF can displace the Gal4 DNA binding domain from its DNA binding site by sliding a nucleosome across the binding site (Li et al. 2015); and human SWI/SNF was found to displace glucocorticoid receptor from its binding sequence in a reconstituted nucleosome array, contributing to the transient nature of GR interactions with the promoter in chromatin (Nagaich et al. 2004). An intriguing possibility therefore is that SWI/SNF functions similarly by sliding nucleosomes that have not been evicted from the NDR across Gcn4 binding motifs to displace bound Gcn4, offsetting the stimulatory effects of nucleosome eviction at these binding sites exerted by CRs (Fig. 7), and that the outcome of these opposing effects differs at different 5’ sites. Our finding that SWI/SNF acts frequently to enhance Gcn4 binding when RSC is depleted suggests that SWI/SNF is capable of functioning like RSC and Ino80C to enhance Gcn4 binding by evicting nucleosomes near the Gcn4 binding motifs. Moreover, even in the presence of RSC, SWI/SNF stimulates Gcn4 binding via nucleosome eviction at a small subset of 5’ sites (eg. sites at top of heat-map in Fig. S9B(ii)). It remains to be determined what features of the NDRs dictate whether the net outcome of SWI/SNF function is to stimulate or suppress Gcn4 binding.

**Figure 7.**
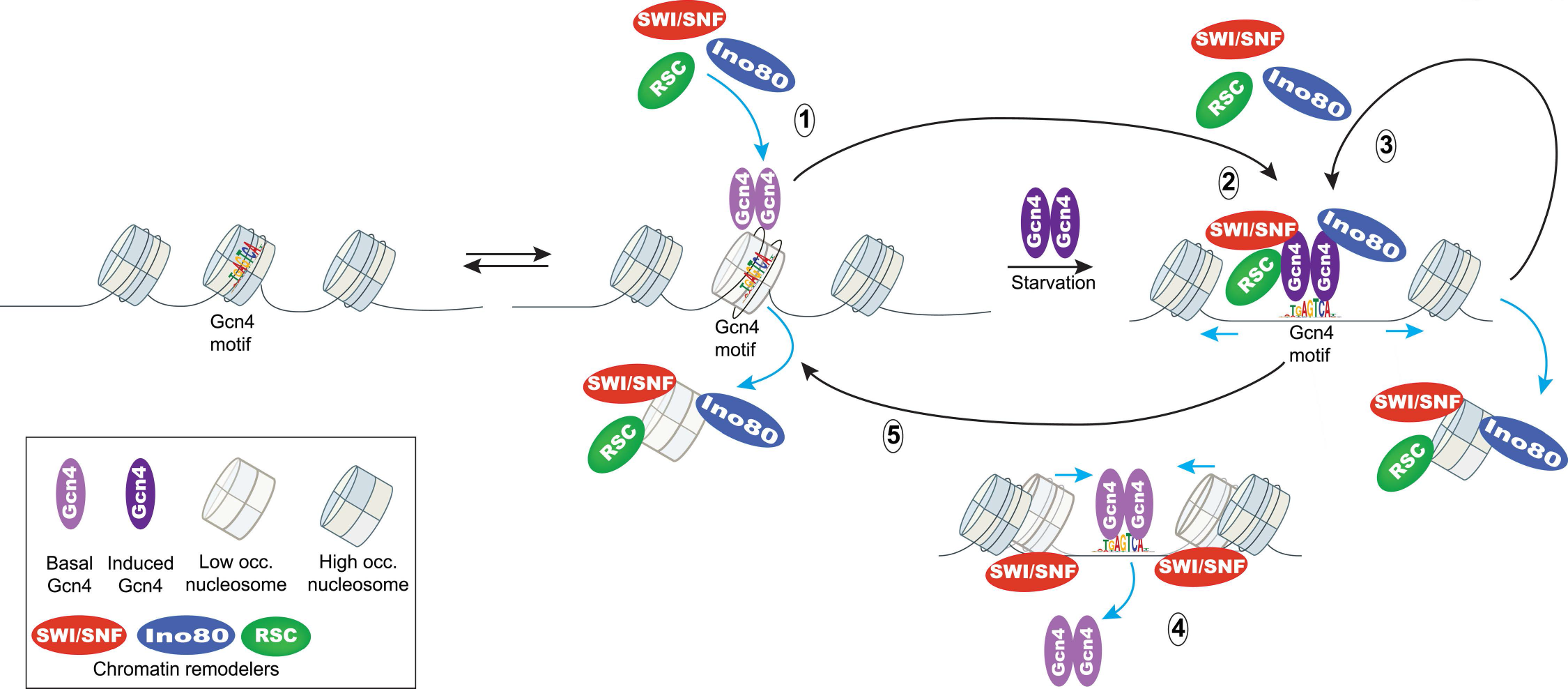
Model depicting positive and negative feedback loops involving SWI/SNF, RSC, and Ino80C in regulating Gcn4 binding to motifs in NDRs of Gcn4 target genes. (1) Basal recruitment of the three CRs reduces nucleosome occupancy of the Gcn4 binding motif in the NDR (depicted by a semi-transparent nucleosome) to facilitate binding of Gcn4 expressed at uninduced levels (light purple). (2) Induction of Gcn4 expression by amino acid starvation (dark purple) leads to increased Gcn4 binding by mass action. The increased Gcn4 occupancy leads to additional recruitment of the CRs, which in turn increases eviction and displacement of nucleosomes from the Gcn4 motif to favor additional Gcn4 binding, thus establishing a positive feedback loop that promotes Gcn4 binding (3). Increased recruitment of SWI/SNF enhances its negative effect on Gcn4 binding by sliding a nucleosome over the Gcn4 motif, leading to dissociation of Gcn4 from the NDR (4), thus establishing a negative feedback loop that prevents excessive Gcn4 binding (5).

It could be argued that if SWI/SNF impedes Gcn4 binding by sliding nucleosomes over the binding motifs, as suggested above and in Fig. 7, then nucleosome occupancies should be decreased in *snf2Δ* cells at the motifs where Snf2 most strongly opposes Gcn4 binding in WT cells. However, we observed little change in H3 occupancies near these sites in *snf2Δ* cells (cf. Fig. 4D(iii) & Fig. 4C(iii), cols. 3 vs 2; see also sites at bottom of heat-map in Fig. S9B(ii)). One way to explain this apparent discrepancy is to propose that the sliding of nucleosomes from either direction over Gcn4 motifs can dislodge Gcn4 without altering the steady-state nucleosome occupancy in the vicinity of the motifs. This might lead to “fuzzier” nucleosome positions near the motifs, which could be detected by high resolution mapping of nucleosome positions, *eg.* that achieved with a chemical cleavage technique (???Chereji et al (henikoff); and the absence of such data is a limitation of the current study. We cannot exclude the possibility that deleting *SNF2* increases Gcn4 binding by an indirect mechanism, eg. altering the expression or activity of an unknown factor that either reduces Gcn4 affinity or competes with Gcn4 for binding motifs. One finding disfavoring such an indirect mechanism, however, is that Snf2 occupancy at the Gcn4 sites is highly correlated with Snf2 function in suppressing Gcn4 binding (Fig. 3D(i) vs. (iv)). We also cannot exclude the possibility that interactions of the CR with Gcn4 and other coactivators or PIC components could favor (or disfavor) Gcn4 binding independently of the established functions of the CRs in evicting or sliding nucleosomes. Clearly, additional work is required to test the model that nucleosome sliding by SWI/SNF acts to limit Gcn4 binding at many 5’ sites.

By ChIP-seq analysis of myc-tagged versions of Sth1, Snf2, and Ino80, we obtained strong evidence that the CRs act directly to modulate Gcn4 binding at 5’ sites, finding that their peak occupancies are centered around the Gcn4 binding motifs. These occupancies increased on induction of Gcn4 by SM treatment, consistent with widespread recruitment of the CRs by Gcn4 to UAS elements. Further evidence for the latter assertion was reported previously by demonstrating reduced SWI/SNF or RSC recruitment at the Gcn4 target gene *ARG1* in a *gcn4Δ* mutant (Swanson et al. 2003; Qiu et al. 2005), and similar evidence was obtained here for Gcn4-dependent Ino80-myc recruitment for the entire population of 5’ Gcn4 binding sites. We envision that basal levels of RSC and Ino80C stimulate Gcn4 binding to the UAS elements as Gcn4 abundance begins to rise in response to starvation, and that the bound Gcn4 then recruits higher levels of the CRs to create a positive feedback loop of mutually stimulatory Gcn4 and CR binding/recruitment as induction proceeds (Fig. 7). Basal levels of SWI/SNF might similarly enhance Gcn4 binding during the early stages of induction, but the increasingly higher levels of SWI/SNF recruited as Gcn4 occupancy rises would frequently begin to impede Gcn4 binding, perhaps by nucleosome sliding, to create a negative feedback loop (Fig. 7). The basal levels of the CRs posited in this model could arise from stochastic interaction with NDRs, or from recruitment by basal levels of Gcn4 or other transcription factors bound to the UAS in non-starved cells. Presumably, the elevated occupancies of the CRs achieved at high-level Gcn4 binding enable them to diffuse laterally from the NDR to mediate the eviction and repositioning of the -1 and +1 nucleosomes we observed in starved WT cells (Qiu et al. 2016) (Rawal et al. 2018a) (Qiu et al. 2020).

Our model that Gcn4 stimulates its own binding by recruitment of CRs that evict nucleosomes from its binding sites (Fig. 7) provides a molecular mechanism that may help to explain previous observations suggesting that an activation domain can facilitate DNA binding by the activator protein. Bunker and Kingston reported that tethering polycomb group proteins to a test promoter differentially interferes with promoter activation by various activation domains that were fused to the Gal4 DNA binding domain (Bunker and Kingston 1994). It is possible that the ability of an activation domain to recruit one or more CRs was instrumental in resisting the repressive effects of polycomb group proteins.

Vashee and Kodadek found that the Gal4 activation domain could outperform the VP16 activation domain when each was linked to the Gal4 DNA binding domain for trans-activation specifically from weak Gal4 binding sites (Vashee and Kodadek 1995). It is possible that the Gal4 AD was more efficient than the VP16 AD in recruiting CRs that eliminated nucleosomes surrounding the Gal4 binding sites in a manner that preferentially facilitated Gal4 binding to non-consensus motifs.

It seems likely that the increased nucleosome occupancies in the NDRs and core promoters of Gcn4 activated genes conferred by the CR mutations observed here and elsewhere (Qiu et al. 2016) (Rawal et al. 2018a) lead to reduced transcription initiation by occluding binding sites for TBP or other general transcription factors involved in PIC assembly. Indeed, we showed recently that reductions in Pol II occupancies in *ino80Δ* cells are associated with both decreased TBP occupancies and increased promoter H3 occupancies at the SM-induced genes (Qiu et al. 2020). Here, we observed additive reductions in TBP binding at genes with 5’ Gcn4 sites on combining the *snf2Δ* and *PTET-STH1* mutations in the double mutant, which was associated with the additive effects of these mutations in elevating nucleosome occupancies at the TBP binding sites. The relatively smaller increases in H3 occupancies observed in the *PTET-STH1* or *snf2Δ* single mutants were associated with lesser reductions in TBP binding in the *PTET-STH1* cells than found in the double mutant, or even increased TBP binding in *snf2Δ* cells. The latter may reflect the elevated Gcn4 occupancies found at most 5’ sites in *snf2Δ* cells, which might enable robust TBP recruitment despite increased nucleosome occlusion of the core promoters.

The functional cooperation between SWI/SNF and RSC in TBP recruitment observed at Gcn4 target genes in SM-treated cells was also observed in non-starved cells for the group of highly expressed RP genes, for which *snf2Δ* impaired TBP recruitment in otherwise WT cells and exacerbated the recruitment defect conferred by *PTET-STH1* in the double mutant. At the large majority of other genes, however, RSC is much more important than SWI/SNF in promoting TBP recruitment in non-starved cells, where it functions *on par* with Ino80C. The greater importance of RSC and Ino80C in TBP recruitment is mirrored by their greater occupancies compared to SWI/SNF at the majority of constitutively expressed genes. It is interesting that genes capable of utilizing SAGA or TFIID as a coactivator have a greater requirement for RSC and Ino80C compared to the genes that rely primarily on TFIID. Our findings support the idea that the three CRs act directly to enhance PIC assembly throughout the genome, with RSC and Ino80C acting broadly and with particular importance at coactivator-redundant genes, whereas SWI/SNF function is concentrated at certain highly expressed genes, including RP genes in non-starved cells and Gcn4-induced genes in amino acid-starved cells.

## MATERIALS AND METHODS

### Plasmid and yeast strain constructions

Yeast strains employed in this study are listed in supplementary Table S1. Yeast strains BY4741 (F729), F731 (*gcn4Δ*) and F748 (*snf2Δ*) were purchased from Research Genetics and described previously (Qiu et al. 2004; Kim et al. 2006). Constructions of *PTET-STH1* strain HQY1632 and *snf2Δ PTET-STH1* strain HQY1660 (Rawal et al. 2018a) and *ino80Δ* strain YR092 (Qiu et al. 2020) were described previously; as were *SNF2-myc* (HQY367), *STH1-myc* (HQY459) (Swanson et al. 2003) and *INO80-myc* (HQY1687) strains (Qiu et al. 2020). Strain HQY1718 (*gcn4Δ INO80-myc*) was constructed from F731 by a PCR-based method for tagging chromosomal genes by yeast transformation (Longtine et al. 1998), using pFA6a-13Myc-His3MX6 DNA as PCR template. To deplete Sth1 from *PTET-STH1* strains, cells were cultured with 10 μg/μl doxycycline for 8-9h, and Sth1 depletion was confirmed by Western analysis.

### ChIP-seq analysis of Gcn4 occupancy

WT strain BY4741 and *snf2Δ or ino80Δ* mutant strains were cultured for at least 3 doublings in synthetic complete medium lacking isoleucine and valine (SC-Ilv) to log-phase (OD600=0.6-0.8) and SM was added at 1 µg/ml for 25 min to induce Gcn4 synthesis. To transcriptionally deplete Sth1 from *PTET-STH1* and *snf2Δ PTET-STH1* strains, cells were cultured in SC-Ilv medium with 10 μg/μl doxycycline for 8-9h to log-phase (OD600=0.4-0.6), allowing 2-3 doublings, and SM was added at 1 µg/ml for 25 min as above. ChIP-seq was conducted and DNA libraries for Illumina paired-end sequencing were prepared as described previously (Rawal et al. 2018a) except that chromatin samples containing 5 µg DNA were immunoprecipitated using Gcn4 antibodies (Zhang et al. 2008) for 3h in experiments comparing WT with *snf2Δ* and *PTET-STH1* mutants, and overnight in experiments comparing WT and *ino80Δ* strains. Biological replicates of Gcn4 ChIP-seq data (and all other ChIP-seq data) are described in Supplementary Table S2.

### ChIP-seq analysis of TBP

For analyzing TBP occupancies, strains were cultured in the presence or absence of SM treatment and subjected to ChIP-seq analysis using antibodies against native TBP, as previously described (Qiu et al. 2020).

### ChIP-seq analysis of Snf2-myc13, Sth1-myc13 and Ino80-myc13

*SNF2-myc* and *STH1-myc* strains were cultured in the presence or absence of SM treatment, and untagged WT strain BY4741 cultured with SM, were subjected to ChIP-seq analysis as described previously (Qiu et al. 2016) with a few modifications. Yeast cells (100 ml) were cross-linked for 15 min with 10 ml formaldehyde solution (50 mM HEPES KOH, pH 7.5, 1 mM EDTA, 100 mM NaCl and 11% formaldehyde) and quenched with 15 ml of 2.5 M glycine. WCEs were prepared by glass-beads lysis in 400 ml FA lysis buffer (50 mM HEPES KOH, pH 7.5, 1 mM EDTA, 150 mM NaCl, 1% TritonX-100 and 0.1% Na-deoxycholate) with protease inhibitors for 45 min at 4°C and the supernatant collected after removing the beads was pooled with 600 ml FA lysis buffer used for washing the beads. The resulting lysate was sonicated to yield DNA fragments of 300–500 bp and cleared by centrifugation. Chromatin samples containing 5.0 μg DNA were immunoprecipitated with anti-myc antibodies (Roche) for 3h. Paired-end sequencing libraries were prepared from immunoprecipitated DNA using Illumina paired-end kits from New England Biolabs (cat. #E7370 and #E7335). ChIP-seq analysis of Ino80-myc was previously described (Qiu et al. 2020). The myc occupancies at each nucleotide, normalized to the average myc occupancy per nucleotide on the chromosome, were calculated for each myc-tagged strain and for the untagged WT and *gcn4Δ* strains, and the latter values were subtracted from the former values to yield the corrected occupancies of the myc-tagged CR subunits in the tagged strains.

### ChIP-seq analysis of Rpb3 and histone H3

ChIP-seq analysis of histone H3 (Abcam, Ab 1791) and Rpb3 (Neo-Clone, W0012) using formaldehyde cross-linked chromatin sheared by sonication were conducted as described previously (Qiu et al. 2016) (Rawal et al. 2018a) (Qiu et al. 2020). Similarly, ChIP-seq analysis of histone H3 (Abcam, Ab1791) following micrococcal nuclease (MNase) digestion was carried out as described previously (Rawal et al. 2018a).

### Bioinformatics analysis of ChIP-seq data

Paired-end sequencing (50 nt from each end) was conducted by the DNA Sequencing and Genomics core facility of the NHLBI, NIH. Sequence data were aligned to the SacCer3 version of the genome sequence using Bowtie2 (Langmead and Salzberg 2012) with parameters *-X 1000 --very-sensitive*, to map sequences up to 1 kb with maximum accuracy. PCR duplicates from ChIP-seq data were removed using the *samtools rmdup* package. Numbers of aligned paired reads from each ChIP-seq experiment and correlation coefficients for genome-wide occupancy profiles of the different replicates are summarized in Table S2. Raw genome-wide occupancy profiles were obtained from the alignment (.bam) files by counting the number of DNA fragments that overlapped with every bp, using the *coverage* methods for *GRanges* objects from the GenomicRanges package in R. To allow the comparison between different samples, each profile was normalized such that the average occupancy for each chromosome was equal to one. Heat maps showing alignments of multiple loci were generated in R using custom scripts (https://github.com/rchereji/bamR). To visualize specific loci, *igvtools* was used to create tracks (*tdf* files) that were loaded in the Integrative Genomics Viewer (IGV, Broad Institute) (Robinson et al. 2011). Transcript end coordinates (TSS and TTS) were obtained from (Pelechano et al. 2013) .

For calculation of per base pair occupancies in the given intervals, generation of averaged occupancies profiles and heat maps of occupancies or occupancy differences surrounding the given genomic features, a custom R scripts package available at github.com/rchereji/bamR was employed. These R scripts, originally designed to plot ChIP-seq occupancies surrounding genic features like TSS and TTS or within coding sequences, were adapted to generate heat maps of Gcn4 occupancy differences within peak coordinates, TBP occupancy differences surrounding the peak summits, and histone H3 occupancies and nucleosome dyad densities surrounding Gcn4 motifs or TBP summits. To do so, alternative genome annotation files were generated with either Gcn4 peak start and end coordinates or 200 bp intervals surrounding the TBP peak summits replacing ORF start and end coordinates; and with Gcn4 motif centers or TBP peak summits replacing TSS coordinates. The annotation information employed for these analyses are provided in supplementary File S1. Box plots, scatter plots and line plots were generated using R package “ggpubr”.

### Identification of TBP peaks

To identify TBP peaks in the gene promoters of Gcn4 5’ target genes, MACS2 (http://liulab.dfci.harvard.edu/MACS/) analysis was employed to TBP ChIP-seq data from two replicates of WT_I cultures (Table S2), using a threshold for the p-value of 10^-3^ (supplementary File S1, sheet “MACS2_TBP_Ind”). Among 117 5’ target genes, we identified TBP peaks in the promoters of 83 5’ target genes (File S1, sheet “TBP_peak_annotations”). MACS2 analysis of WT_U TBP ChIP-seq data was conducted as above using a p-value of 10^-3^ for identifying the TBP peak within the promoter of 134 RP genes (File S1, sheet “MACS2_TBP_Unind”). TBP peaks were assigned to specific genes by using the bedtools utility “closest” to identify the TBP peak summit closest to the TSS of each gene, followed by assessment of the TBP summit by examining the TBP data in the IGV browser. As tDNA genes often reside in the 5’ non-coding regions of Pol II-transcribed genes and display large TBP peaks in both induced and uninduced conditions, the TBP occupancies within 250 bp surrounding tDNA genes (File S1, sheet “tDNA_annotations”) were subtracted from the TBP coverage in the bed file of merged replicate data and the corrected bed file was used to generate averaged profiles and calculate TBP occupancies within 200 bp intervals surrounding the TBP summits. The computation of TBP occupancies in the intervals comprised of 100 bp on either side of the TBP summits, as done in Fig 5B, was found to be unaffected if the TBP occupancies at neighboring tDNA genes were not subtracted in the manner just described.

## DATA AND SOFTWARE AVAILABILITY

File S1 contains annotations of Gcn4 and TBP occupancy peaks. Results of MACS2 analysis of TBP ChIP-seq data and alternative annotation files listing the coordinates of either Gcn4 occupancy peaks and Gcn4 motifs or TBP occupancy peaks and TBP peak occupancies, in place of ORF start and end coordinates and TSS, used in plotting heat maps of Gcn4 or TBP occupancies. A file listing the genomic coordinates of the 250 bp surrounding each tDNA gene is also provided. File S2 contains the data used to construct all plots. Raw and analyzed ChIP-seq data have been deposited in the NCBI GEO database under accession number GSE192592.

## CONFLICT OF INTEREST STATEMENT

The authors declare that they have no conflict of interest.

## ACKNOWLEDGEMENTS

We thank Razvan Chereji and David Clark for many helpful discussions, and Joseph Reese for his generous gift of TBP antibodies. This work was supported in part by the Intramural Research Program of the National Institutes of Health.

## SUPPLEMENTARY TABLE LEGEND

Table S1. **Yeast strains used in this study.** Names, genotypes and sources of each strain are given.

Table S2. **Compilation of ChIP-seq replicate experiments.** The strain and growth condition (SM-induced (I), or uninduced (U)), immunoprecipitating antiserum (IP), sample identification number, total number of sequencing reads (All PE reads), number of reads after removing duplicate reads (PE rmdup), correlation coefficient between indicated replicates, and source of data, are given for each ChIP-seq experiment.

## SUPPLEMENTARY FIGURE LEGENDS

Figure S1. **Supporting evidence that SWI/SNF and RSC have opposing effects on Gcn4 binding at 5’ sites.** Paired box pots of log2 Gcn4 occupancies comparing (A) WT_I versus *PTET-STH1_*I, (B) *PTET-STH1_*I versus WT_I and *snf211 PTET-STH1_*I and (C) WT_I versus *snf211*_I in 3 sets of Gcn4 5’ sites comprised of the (i) first (Set_1, n=30), (ii) middle two (Set_2, n=57) and (iii) last (Set_3, n=30) quartiles of the fold-changes in Gcn4 occupancy in *snf211 PTET-STH1_*I vs. WT_I cells as depicted in Fig. 2B(i). Lines connecting each data point in respective strains indicate changes in the Gcn4 occupancies of respective Gcn4 site.

Figure S2. **Gene browser profiles of Gcn4 and H3 occupancies from ChIP-seq analyses of sonicated chromatin for the indicated strains**. The Gcn4 peak numbering assigned previously (Rawal et al. 2018b) is given at the top of each profile, and the Gcn4 occupancies per nucleotide averaged over the peaks determined in this study are listed next to each peak.

Figure S3. **Differential requirements for Ino80C and RSC for Gcn4 binding at a subset of Gcn4 5’ sites. (A)** Heat maps of differences in Gcn4 occupancies averaged across the coordinates of 5’ sites between the indicated mutant and WT_I samples for (i) *ino80Δ*_I and (ii) *snf211 PTET-STH1*_I. Gcn4 5’ sites were sorted by increasing order of the ratio of Gcn4 occupancies in the double mutant *snf211 PTET-STH1*_I vs. WT_I. **(B)** Heat map depictions of Gcn4 occupancies surrounding the Gcn4 motifs of 5’ sites in (i) WT_I (for the *ino80Δ* mutant), (ii) *ino80Δ*_I, (iii) WT_I (for the *snf211 PTET-STH1*_I mutant), and (iv) *snf211 PTET-STH1*_I cells, for the same ordering of 5’ sites as in (A). The sets of Gcn4 5’ sites (Set_1, Set_2, and Set_3) defined in Fig. 2A-B are depicted in B(i). **(C & D)** Same analyses for *ino80Δ*_I and *PTET-STH1*_I as shown in (A-B) except for Gcn4 ORF sites.

Figure S4. **Identification of Gcn4 5’ sites with heightened Ino80C dependence for Gcn4 occupancy. (A)** Heat map depicting differences in Gcn4 occupancies between *ino8011*_I and WT_I cells (i); Gcn4 occupancies surrounding the motifs of 5’ sites in (ii) WT_I or (iii) *ino8011*_I cells. Gcn4 5’ sites were sorted by increasing order of the ratio of Gcn4 occupancies in *ino8011*_I vs. WT_I cells, and the first (Set_I, n=30), middle two (Set_II, n=57) and fourth (Set_III, n=30) quartiles of fold-changes are depicted in A(ii). **(B)** Heat map depictions of Gcn4 occupancies between (i) *snf211 PTET-STH1*_I and WT_I, (ii) *PTET-STH1_*I and WT_I, and (iii) *snf2*_I vs. WT_I cells. **(C)** Notched box pots of log2 Gcn4 occupancy in WT_U, WT_I, and *ino8011*_I cells in 3 sets of Gcn4 5’ sites comprised of the (i) first (Set_I, n=30), (ii) middle two (Set_II, n=57) and (iii) last (Set_III, n=30) quartiles of the fold-changes in Gcn4 occupancy in *ino8011_I* vs. WT_I cells as defined in panel A(ii). *P* values for the significance of differences in medians calculated by the Mann-Whitney-Wilcoxon test are indicated. **(D)** Scatterplots of log2 ratios of Gcn4 occupancy changes in WT_I vs. *ino8011*_I cells plotted against log2 Gcn4 occupancies in WT_I cells for 5’ (i) and ORF (ii) Gcn4 sites. Pearson correlation coefficients (*R*) and associated *p* values are indicated.

Figure S5. Supporting evidence that reduced Gcn4 binding in mutants depleted of RSC or Ino80C occurs preferentially at motifs of highest affinity or accessibility in chromatin in WT_I cells. (A & B) Histograms depicting (A) log2 Rpb3 occupancies in the *GCN4* CDS measured by Rpb3 ChIP-seq, indicating transcription levels; and (B) Gcn4 protein levels measured previously (Rawal et al. 2018a) by Western blot analysis in the indicated strains using Gcd6 signals analyzed in parallel as loading control. Band intensities for Gcn4 were normalized to those for Gcd6 in the same samples and the mean Gcn4/Gcd6 ratios determined from 3 biological replicates were plotted. Significance of differences in mean values was calculated with the student’s t test. (C & D) Scatterplots of log2 WT_I Gcn4 occupancies vs. motif FIMO scores (Rawal et al. 2018b) for (C) the 115 Gcn4 5’ sites and (D) the 62 Gcn4 ORF peaks, using the motif of highest score for peaks with multiple motifs. Pearson correlation coefficients (*R*) and associated *p* values are indicated. (E) Notched box pots of motif FIMO scores for the sets of Gcn4 5’ sites binned according to the fold-changes in occupancy in *snf211 PTET-STH1_*I vs. WT_I cells, as depicted in Fig. 2B.

Figure S6. Gene browser profiles of Gcn4 and corrected Snf2-myc occupancies from ChIP-seq analyses of sonicated chromatin for the indicated strains. The Gcn4 peak numbering assigned previously (Rawal et al. 2018b) is given at the top of each profile, and corrected Snf2-myc occupancies per nucleotide within ±100 bp windows surrounding the Gcn4 motifs are listed next to each peak.

Figure S7. **Supporting evidence for recruitment of the three CRs by Gcn4 to its 5’ sites.** (**A) (i)-(iii)** Scatterplots of corrected occupancies of the indicated myc-tagged CR subunits in WT_I cells measured within ±100 bp windows surrounding the Gcn4 motifs of 5’ Gcn4 peaks. Pearson correlation coefficients (*R*) and associated *p* values are indicated. **(B)** Heat map depictions of corrected Ino80-myc occupancies at the 5’ Gcn4 peaks, sorted by decreasing Gcn4 occupancies in WT_I cells and plotted relative to the Gcn4 motifs, for (i) uninduced *gcn4Δ* cells, (ii) SM-treated *gcn4Δ* cells, (iii) uninduced WT cells, and (iv) SM-treated WT cells. Occupancies were calculated from ChIP-seq data of mildly sonicated chromatin from 2 or 3 biological replicates each of isogenic *GCN4* or *gcn4Δ* strains, harboring *INO80-myc* or untagged *INO80,* under inducing or uninducing conditions, correcting the occupancies for *INO80-myc* cells for those measured for the untagged *INO80* cells of the same *GCN4* genotype and growth conditions. **(C)** Notched box plots of corrected Ino80-myc occupancies per nucleotide within ±100 bp windows surrounding the Gcn4 motifs of 5’ sites in uninduced (_U) or SM induced (_I) *gcn4Δ* or WT cells.

Figure S8. **Supporting evidence for defective eviction of nucleosomes surrounding 5’ Gcn4 motifs in SWI/SNF and RSC mutants.** Sectored scatterplot of the log2 ratios of Gcn4 occupancies vs log2 ratios of H3 occupancies per base pair in the ±100 bp windows surrounding the Gcn4 motifs in (A) *snf211 PTET-STH1_*I (B) *snf211_*I vs. WT_I cells. The 3 sets of Gcn4 5’ sites comprised of the (i) first (Set_1, red rectangles), (ii) middle two (Set_2, green pluses) and (iii) last (Set_3, blue stars) quartiles of the fold-changes in Gcn4 occupancy in *snf211 PTET-STH1_*I vs. WT_I cells are depicted by different color coding and point shapes.

Figure S9. Defective eviction of nucleosomes associated with reduced Gcn4 occupancies at a subset of 5’ Gcn4 peaks in *snf211*_I cells. (A) (i)-(iii) Heat maps depicting differences between *snf211*_I and WT_I cells for (i) Gcn4 occupancies measured as in Fig 2A(iii), (ii) H3 occupancies surrounding the Gcn4 motifs of 5’ sites from H3 ChIP-seq data, and (iii) Rpb3 occupancies averaged over the CDS of 5’ genes, for the Gcn4 5’ sites sorted by increasing order of fold-changes in Gcn4 occupancies in *snf211 PTET-STH1*_I vs. WT_I cells. (B) (i)-(iii) Same analyses shown in (A) except sorted by increasing order of fold-changes in Gcn4 occupancies in *snf211*_I vs. WT_I cells. (C) (i)-(iii) Same analyses shown in (A) except for *PTET-STH1*_I vs. WT_I data.

Figure S10. Defective eviction of nucleosomes associated with reduced Gcn4 occupancies at a subset of 5’ Gcn4 peaks in *ino8011*_I cells. (A) (i)-(iii) Heat maps depicting differences between *ino8011*_I and WT_I cells for (i) Gcn4 occupancies measured as in Fig S4A(i), (ii) H3 occupancies surrounding the Gcn4 motifs of 5’ sites from H3 ChIP-seq data, and (iii) Rpb3 occupancies averaged over the CDS of 5’ genes, for Gcn4 5’ sites sorted by increasing order of fold-changes in Gcn4 occupancies in *ino8011*_I vs. WT_I cells. (B-D) Notched box plots for the 3 sets of 5’ sites defined in Fig. S4A(ii), and indicated again in panel A(ii), depicting (B) H3 occupancies per base pair in the ±100 bp windows surrounding the Gcn4 motifs, (C) log2 Gcn4 occupancies taken from Fig. S4C, and (D) log2 Rpb3 occupancies averaged over the CDS of genes with 5’ sites. H3 and Rpb3 occupancies were calculated from ChIP-seq data of sonicated chromatin from at least 3 biological replicates of WT_U, WT_I and *ino8011*_I cells. *P* values from Mann-Whitney-Wilcoxon tests are indicated. The heat map of H3 occupancy changes conferred by *ino80Δ* around the 5’ motifs ordered by the Gcn4 occupancy reductions in this mutant (panel A(i)) reveals that the 5’ sites with the strongest reductions in Gcn4 binding in *ino80Δ* cells located at the top of the map (Set_I) show the strongest increases in H3 occupancies centered around the Gcn4 motifs (panel A(ii)). Moreover, the decreases in median Gcn4 occupancy for individual 5’ sites conferred by *ino80Δ* are paralleled by increased median H3 occupancies for the Set_I group of 5’ sites; whereas the sites in Set_II and III show no significant changes in median H3 or Gcn4 occupancies in *ino80Δ*_I versus WT_I cells (panels 10B-C, Sets_I-III, col. 3 vs. 2). The subset of genes with 5’ sites that are most dependent on Ino80C for Gcn4 binding generally show the greatest reductions in Rpb3 occupancies (Set_I sites in panel A (iii) vs. (i)). Moreover, the Set_I genes, but not genes in Sets_II-III, show reduced median occupancies of both Gcn4 and Rpb3 in *ino80Δ*_I versus WT_I cells (panels C-D(i)-(iii), col. 3 vs. 2). As only the Set_I genes also exhibit increased median H3 occupancies in *ino80Δ*_I cells (panel B(i)-(iii), col. 3 vs. 2), it seems likely that a defect in Gcn4 binding (and subsequent impaired recruitment of other coactivators) in combination with loss of Ino80C-mediated promoter nucleosome eviction produces the reduced transcription of Set_I genes conferred by *ino80Δ*. (E) Gene browser profiles of Gcn4 and H3 occupancies from ChIP-seq analyses of sonicated chromatin for the indicated strains, as described in Fig. S2.

Figure S11. **Decreased TBP recruitment at 5’ and ORF genes in *ino8011*_I cells. (A-B)** Averaged TBP occupancies surrounding the Gcn4 motifs in ORF peaks from ChIP-seq data of sonicated chromatin using anti-TBP antibodies from at least 2 biological replicates for the indicated mutant and WT strains. **(C)** Notched box plots of factor occupancies in the 83 5’ gene promoters for the two bins defined in Fig. 5C for (i) log2 TBP within ±100 bp of TBP peak summits and (ii) H3 within ±100 bp of the TBP peak summits. *P* values from Mann-Whitney-Wilcoxon tests are indicated. **(D-E)** Gene browser profiles of TBP, Rpb3, and H3 occupancies from ChIP-seq analyses of sonicated chromatin for the indicated strains. The TBP occupancies per nucleotide over the TBP peaks and Rpb3 occupancies per nucleotide over the CDS are listed next to the relevant peaks or CDSs. The locations of the Gcn4 motifs and TBP summits are indicated with vertical hash marks, and the positions of -1 and +1 nucleosomes with dashes, at the bottom of each profile.

Figure S12. Gene browser profiles of TBP, Rpb3, and H3 occupancies from ChIP-seq analyses of sonicated chromatin for the indicated strains. The TBP occupancies per nucleotide over the TBP peaks and Rpb3 occupancies per nucleotide over the CDS are listed next to the relevant peaks or CDSs. The locations of the Gcn4 motifs and TBP summits are indicated with vertical hash marks, and the positions of -1 and +1 nucleosomes with dashes, at the bottom of each profile.

Figure S13. Coactivator-redundant genes have a greater requirement than TFIID-dependent genes for RSC and Ino80C for TBP recruitment. Changes in TBP occupancies surrounding the TSSs in the indicated mutants versus WT under non-starvation conditions are plotted for the coactivator-redundant and TFIID-dependent genes defined by Donczew et al. (2020), along with the corresponding changes for the RPGs.

## SUPPLEMENTARY FILE LEGENDS

File S1. Annotations of the start and end coordinates of Gcn4 ChIP-seq occupancy peaks and of 200 bp intervals surrounding the peak summits of TBP ChIP-seq occupancy peaks.

File S2. Compilation of all data used to construct plots in both main and supplementary figures.

## Notes

### Competing Interest Statement

The authors have declared no competing interest.

## REFERENCES

1. Adkins MW, Williams SK, Linger J, Tyler JK. 2007. Chromatin disassembly from the PHO5 promoter is essential for the recruitment of the general transcription machinery and coactivators. Mol Cell Biol 27: 6372–6382.

2. Badis G, Chan ET, van Bakel H, Pena-Castillo L, Tillo D, Tsui K, Carlson CD, Gossett AJ, Hasinoff MJ, Warren CL et al. 2008. A library of yeast transcription factor motifs reveals a widespread function for Rsc3 in targeting nucleosome exclusion at promoters. Mol Cell 32: 878–887.

3. Barbaric S, Luckenbach T, Schmid A, Blaschke D, Horz W, Korber P. 2007. Redundancy of chromatin remodeling pathways for the induction of the yeast PHO5 promoter in vivo. J Biol Chem 282: 27610–27621.

4. Boeger H, Griesenbeck J, Strattan JS, Kornberg RD. 2003. Nucleosomes unfold completely at a transcriptionally active promoter. Mol Cell 11: 1587–1598.

5. Brahma S, Udugama MI, Kim J, Hada A, Bhardwaj SK, Hailu SG, Lee TH, Bartholomew B. 2017. INO80 exchanges H2A.Z for H2A by translocating on DNA proximal to histone dimers. Nat Commun 8: 15616.

6. Bunker CA, Kingston RE. 1994. Transcriptional repression by Drosophila and mammalian Polycomb group proteins in transfected mammalian cells. Mol Cell Biol 14: 1721–1732.

7. Cui F, Cole HA, Clark DJ, Zhurkin VB. 2012. Transcriptional activation of yeast genes disrupts intragenic nucleosome phasing. Nucleic Acids Res 40: 10753–10764.

8. Devlin C, Tice-Baldwin K, Shore D, Arndt KT. 1991. RAP1 is required for BAS1/BAS2- and GCN4- dependent transcription of the yeast }U}HIS4}u} gene. Mol Cell Biol 11: 3642–3651.

9. Donczew R, Warfield L, Pacheco D, Erijman A, Hahn S. 2020. Two roles for the yeast transcription coactivator SAGA and a set of genes redundantly regulated by TFIID and SAGA. Elife 9: e50109.

10. Ganguli D, Chereji RV, Iben JR, Cole HA, Clark DJ. 2014. RSC-dependent constructive and destructive interference between opposing arrays of phased nucleosomes in yeast. Genome Res 24: 1637–1649.

11. Gowans GJ, Schep AN, Wong KM, King DA, Greenleaf WJ, Morrison AJ. 2018. INO80 Chromatin Remodeling Coordinates Metabolic Homeostasis with Cell Division. Cell Rep 22: 611–623.

12. Hartley PD, Madhani HD. 2009. Mechanisms that specify promoter nucleosome location and identity. Cell 137: 445–458.

13. Jeronimo C, Watanabe S, Kaplan CD, Peterson CL, Robert F. 2015. The Histone Chaperones FACT and Spt6 Restrict H2A.Z from Intragenic Locations. Mol Cell 58: 1113–1123.

14. Jia MH, Larossa RA, Lee JM, Rafalski A, Derose E, Gonye G, Xue Z. 2000. Global expression profiling of yeast treated with an inhibitor of amino acid biosynthesis, sulfometuron methyl. Physiol Genomics 3: 83–92.

15. Jiang C, Pugh BF. 2009. Nucleosome positioning and gene regulation: advances through genomics. Nat Rev Genet 10: 161–172.

16. Kim Y, McLaughlin N, Lindstrom K, Tsukiyama T, Clark DJ. 2006. Activation of Saccharomyces cerevisiae HIS3 results in Gcn4p-dependent, SWI/SNF-dependent mobilization of nucleosomes over the entire gene. Mol Cell Biol 26: 8607–8622.

17. Klein-Brill A, Joseph-Strauss D, Appleboim A, Friedman N. 2019. Dynamics of Chromatin and Transcription during Transient Depletion of the RSC Chromatin Remodeling Complex. Cell Rep 26: 279–292 e275.

18. Klopf E, Schmidt HA, Clauder-Munster S, Steinmetz LM, Schuller C. 2017. INO80 represses osmostress induced gene expression by resetting promoter proximal nucleosomes. Nucleic Acids Res 45: 3752–3766.

19. Krietenstein N, Wal M, Watanabe S, Park B, Peterson CL, Pugh BF, Korber P. 2016. Genomic Nucleosome Organization Reconstituted with Pure Proteins. Cell 167: 709–721 e712.

20. Kuang Z, Cai L, Zhang X, Ji H, Tu BP, Boeke JD. 2014. High-temporal-resolution view of transcription and chromatin states across distinct metabolic states in budding yeast. Nat Struct Mol Biol 21: 854–863.

21. Kubik S, Bruzzone MJ, Challal D, Dreos R, Mattarocci S, Bucher P, Libri D, Shore D. 2019. Opposing chromatin remodelers control transcription initiation frequency and start site selection. Nat Struct Mol Biol 26: 744–754.

22. Kubik S, O’Duibhir E, de Jonge WJ, Mattarocci S, Albert B, Falcone JL, Bruzzone MJ, Holstege FCP, Shore D. 2018. Sequence-Directed Action of RSC Remodeler and General Regulatory Factors Modulates +1 Nucleosome Position to Facilitate Transcription. Mol Cell 71: 89–102 e105.

23. Langmead B, Salzberg SL. 2012. Fast gapped-read alignment with Bowtie 2. Nat Methods 9: 357–359.

24. Levo M, Avnit-Sagi T, Lotan-Pompan M, Kalma Y, Weinberger A, Yakhini Z, Segal E. 2017. Systematic Investigation of Transcription Factor Activity in the Context of Chromatin Using Massively Parallel Binding and Expression Assays. Mol Cell 65: 604–617 e606.

25. Li M, Hada A, Sen P, Olufemi L, Hall MA, Smith BY, Forth S, McKnight JN, Patel A, Bowman GD et al. 2015. Dynamic regulation of transcription factors by nucleosome remodeling. Elife 4: e06249.

26. Liu X, Lee CK, Granek JA, Clarke ND, Lieb JD. 2006. Whole-genome comparison of Leu3 binding in vitro and in vivo reveals the importance of nucleosome occupancy in target site selection. Genome Res 16: 1517–1528.

27. Longtine MS, McKenzie III A, Demarini DJ, Shah NG, Wach A, Brachat A, Philippsen P, Pringle JR. 1998. Additonal modules for versatile and economical PCR-based gene deletion and modification in *Saccharomyces cerevisiae*. Yeast 14: 953–961.

28. Mehta GD, Ball DA, Eriksson PR, Chereji RV, Clark DJ, McNally JG, Karpova TS. 2018. Single-Molecule Analysis Reveals Linked Cycles of RSC Chromatin Remodeling and Ace1p Transcription Factor Binding in Yeast. Mol Cell 72: 875–887 e879.

29. Mizuguchi G, Shen X, Landry J, Wu WH, Sen S, Wu C. 2004. ATP-driven exchange of histone H2AZ variant catalyzed by SWR1 chromatin remodeling complex. Science 303: 343–348.

30. Musladin S, Krietenstein N, Korber P, Barbaric S. 2014. The RSC chromatin remodeling complex has a crucial role in the complete remodeler set for yeast PHO5 promoter opening. Nucleic Acids Res 42: 4270–4282.

31. Nagaich AK, Walker DA, Wolford R, Hager GL. 2004. Rapid periodic binding and displacement of the glucocorticoid receptor during chromatin remodeling. Mol Cell 14: 163–174.

32. Natarajan K, Jackson BM, Zhou H, Winston F, Hinnebusch AG. 1999. Transcriptional activation by Gcn4p involves independent interactions with the SWI/SNF complex and SRB/mediator. MolCell 4: 657–664.

33. Natarajan K, Meyer MR, Jackson BM, Slade D, Roberts C, Hinnebusch AG, Marton MJ. 2001. Transcriptional profiling shows that Gcn4p is a master regulator of gene expression during amino acid starvation in yeast. Molecular and Cellular Biology 21: 4347–4368.

34. Nocetti N, Whitehouse I. 2016. Nucleosome repositioning underlies dynamic gene expression. Genes Dev 30: 660–672.

35. Parnell TJ, Huff JT, Cairns BR. 2008. RSC regulates nucleosome positioning at Pol II genes and density at Pol III genes. Embo J 27: 100–110.

36. Parnell TJ, Schlichter A, Wilson BG, Cairns BR. 2015. The chromatin remodelers RSC and ISW1 display functional and chromatin-based promoter antagonism. Elife 4: e06073.

37. Pelechano V, Wei W, Steinmetz LM. 2013. Extensive transcriptional heterogeneity revealed by isoform profiling. Nature 497: 127–131.

38. Qiu H, Biernat E, Govind CK, Rawal Y, Chereji RV, Clark DJ, Hinnebusch AG. 2020. Chromatin remodeler Ino80C acts independently of H2A.Z to evict promoter nucleosomes and stimulate transcription of highly expressed genes in yeast. Nucleic Acids Res 48: 8408–8430.

39. Qiu H, Chereji RV, Hu C, Cole HA, Rawal Y, Clark DJ, Hinnebusch AG. 2016. Genome-wide cooperation by HAT Gcn5, remodeler SWI/SNF, and chaperone Ydj1 in promoter nucleosome eviction and transcriptional activation. Genome Res 26: 211–225.

40. Qiu H, Hu C, Yoon S, Natarajan K, Swanson MJ, Hinnebusch AG. 2004. An array of coactivators is required for optimal recruitment of TATA binding protein and RNA polymerase II by promoter-bound Gcn4p. Mol Cell Biol 24: 4104–4117.

41. Qiu H, Hu C, Zhang F, Hwang GJ, Swanson MJ, Boonchird C, Hinnebusch AG. 2005. Interdependent recruitment of SAGA and Srb mediator by transcriptional activator Gcn4p. Mol Cell Biol 25: 3461–3474.

42. Rando OJ, Winston F. 2012. Chromatin and transcription in yeast. Genetics 190: 351–387.

43. Rawal Y, Chereji RV, Qiu H, Ananthakrishnan S, Govind CK, Clark DJ, Hinnebusch AG. 2018a. SWI/SNF and RSC cooperate to reposition and evict promoter nucleosomes at highly expressed genes in yeast. Genes Dev 32: 695–710.

44. Rawal Y, Chereji RV, Valabhoju V, Qiu H, Ocampo J, Clark DJ, Hinnebusch AG. 2018b. Gcn4 Binding in Coding Regions Can Activate Internal and Canonical 5’ Promoters in Yeast. Mol Cell 70: 297–311 e294.

45. Reinke H, Horz W. 2003. Histones are first hyperacetylated and then lose contact with the activated PHO5 promoter. Mol Cell 11: 1599–1607.

46. Reja R, Vinayachandran V, Ghosh S, Pugh BF. 2015. Molecular mechanisms of ribosomal protein gene coregulation. Genes Dev 29: 1942–1954.

47. Rhee HS, Pugh BF. 2012. Genome-wide structure and organization of eukaryotic pre-initiation complexes. Nature 483: 295–301.

48. Robinson JT, Thorvaldsdottir H, Winckler W, Guttman M, Lander ES, Getz G, Mesirov JP. 2011. Integrative genomics viewer. Nat Biotechnol 29: 24–26.

49. Rosonina E, Yurko N, Li W, Hoque M, Tian B, Manley JL. 2014. Threonine-4 of the budding yeast RNAP II CTD couples transcription with Htz1-mediated chromatin remodeling. Proc Natl Acad Sci U S A 111: 11924–11931.

50. Saint M, Sawhney S, Sinha I, Singh RP, Dahiya R, Thakur A, Siddharthan R, Natarajan K. 2014. The TAF9 C-terminal conserved region domain is required for SAGA and TFIID promoter occupancy to promote transcriptional activation. Mol Cell Biol 34: 1547–1563.

51. Schwabish MA, Struhl K. 2007. The Swi/Snf complex is important for histone eviction during transcriptional activation and RNA polymerase II elongation in vivo. Mol Cell Biol 27: 6987–6995.

52. Sharma VM, Li B, Reese JC. 2003. SWI/SNF-dependent chromatin remodeling of RNR3 requires TAF(II)s and the general transcription machinery. Genes Dev 17: 502–515.

53. Shivaswamy S, Iyer VR. 2008. Stress-dependent dynamics of global chromatin remodeling in yeast: dual role for SWI/SNF in the heat shock stress response. Mol Cell Biol 28: 2221–2234.

54. Swanson MJ, Qiu H, Sumibcay L, Krueger A, Kim S-J, Natarajan K, Yoon S, Hinnebusch AG. 2003. A Multiplicity of coactivators is required by Gcn4p at individual promoters in vivo. MolCellBiol 23: 2800–2820.

55. van Bakel H, Tsui K, Gebbia M, Mnaimneh S, Hughes TR, Nislow C. 2013. A compendium of nucleosome and transcript profiles reveals determinants of chromatin architecture and transcription. PLoS Genet 9: e1003479.

56. Vashee S, Kodadek T. 1995. The activation domain of GAL4 protein mediates cooperative promoter binding with general transcription factors }U}in}u} }U}vivo}u}. Proc Natl Acad Sci USA 92: 10683–10687.

57. Wang X, Bai L, Bryant GO, Ptashne M. 2011. Nucleosomes and the accessibility problem. Trends Genet 27: 487–492.

58. Xue Y, Pradhan SK, Sun F, Chronis C, Tran N, Su T, Van C, Vashisht A, Wohlschlegel J, Peterson CL et al. 2017. Mot1, Ino80C, and NC2 Function Coordinately to Regulate Pervasive Transcription in Yeast and Mammals. Mol Cell 67: 594–607 e594.

59. Yen K, Vinayachandran V, Batta K, Koerber RT, Pugh BF. 2012. Genome-wide Nucleosome Specificity and Directionality of Chromatin Remodelers. Cell 149: 1461–1473.

60. Yu L, Morse RH. 1999. Chromatin opening and transactivator potentiation by RAP1 in Saccharomyces cerevisiae. Mol Cell Biol 19: 5279–5288.

61. Zhang F, Gaur NA, Hasek J, Kim SJ, Qiu H, Swanson MJ, Hinnebusch AG. 2008. Disrupting vesicular trafficking at the endosome attenuates transcriptional activation by Gcn4. Mol Cell Biol 28: 6796–6818.

