## Supplemental Figures and Tables for "Distinct functions of three chromatin remodelers in activator binding and preinitiation complex assembly"

### Figure_S1

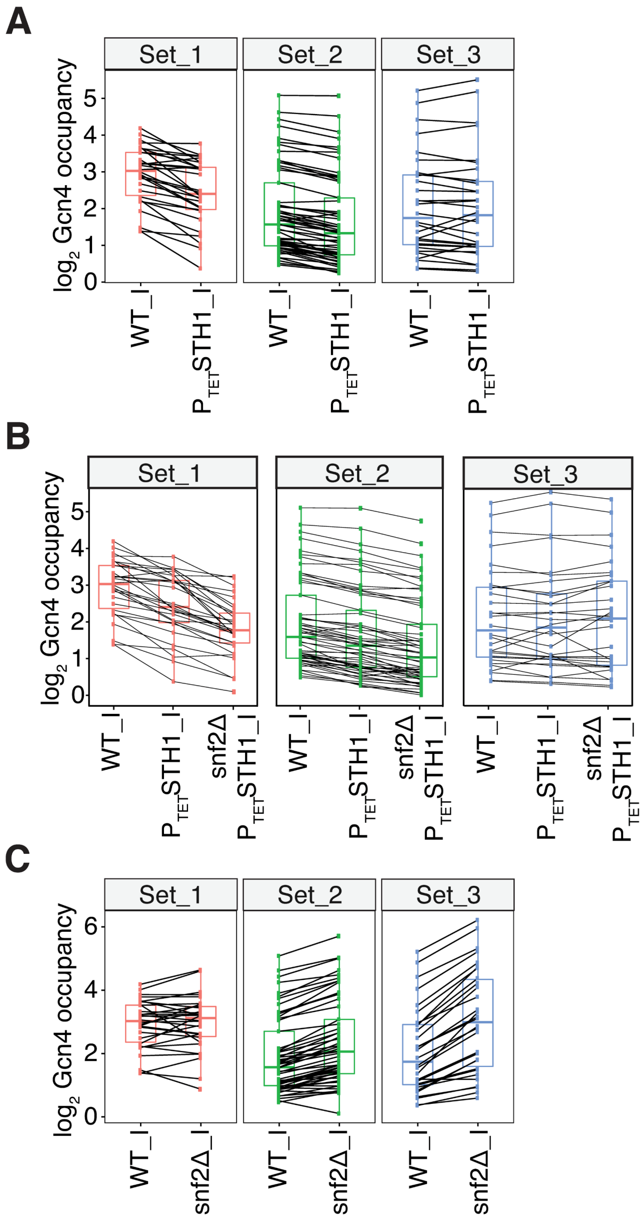


**Figure S1. Supporting evidence that SWI/SNF and RSC have opposing effects on Gcn4 binding at 5' sites.** Paired box pots of log$_2$ Gcn4 occupancies comparing (A) WT_I versus $P_{TET}$-*STH1*_I, (B) $P_{TET}$-*STH1*_I versus WT_I and *snf2Δ* $P_{TET}$-*STH1*_I and (C) WT_I versus *snf2Δ*_I in the three sets of Gcn4 5' sites defined in Fig. 2B(i). Lines connecting two data points in the respective strains indicate the change in Gcn4 occupancies at that Gcn4 site between the two strains under comparison.

#### Figure_S2

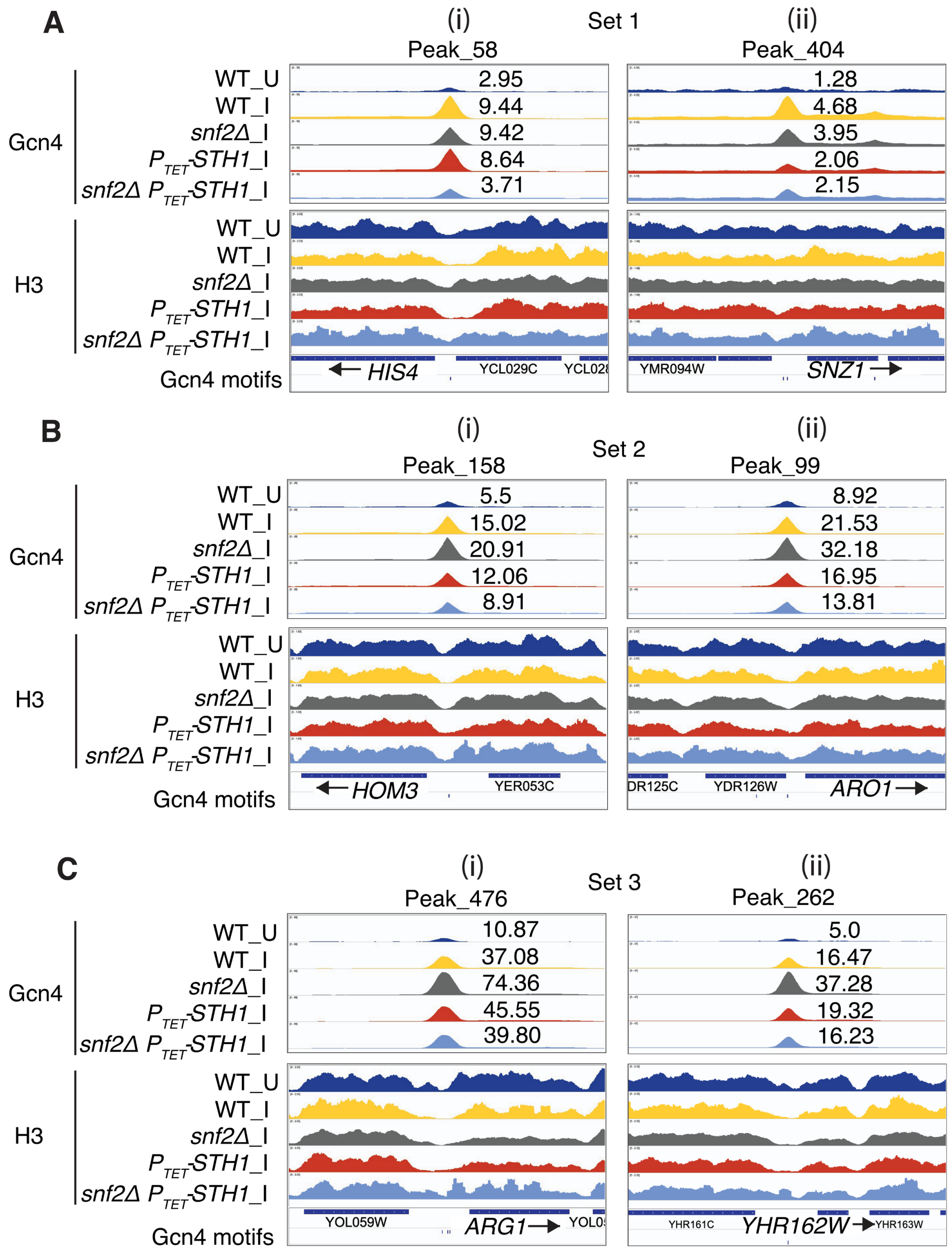
A
(i)
Set 1
(ii)
Peak_58
Peak_404
Gcn4
WT_U
WT_I
snf2Δ_I
P_TET-STH1_I
snf2Δ P_TET-STH1_I
2.95
9.44
9.42
8.64
3.71
1.28
4.68
3.95
2.06
2.15
H3
Gcn4 motifs
HIS4
YCL029C
YMR094W
SNZ1
B
Set 2
Peak_158
Peak_99
5.5
15.02
20.91
12.06
8.91
8.92
21.53
32.18
16.95
13.81
HOM3
YER053C
DR125C
YDR126W
ARO1
C
Set 3
Peak_476
Peak_262
10.87
37.08
74.36
45.55
39.80
5.0
16.47
37.28
19.32
16.23
YOL059W
ARG1
YHR161C
YHR162W
YHR163W

**Figure S2. Gene browser profiles of Gcn4 and H3 occupancies from ChIP-seq analyses of sonicated chromatin for the indicated strains**. The Gcn4 peak numbering assigned previously (Rawal et al. 2018b) is given at the top of each profile, and the Gcn4 occupancies per nucleotide averaged over the peaks determined in this study are listed next to each peak.

#### Figure_S3

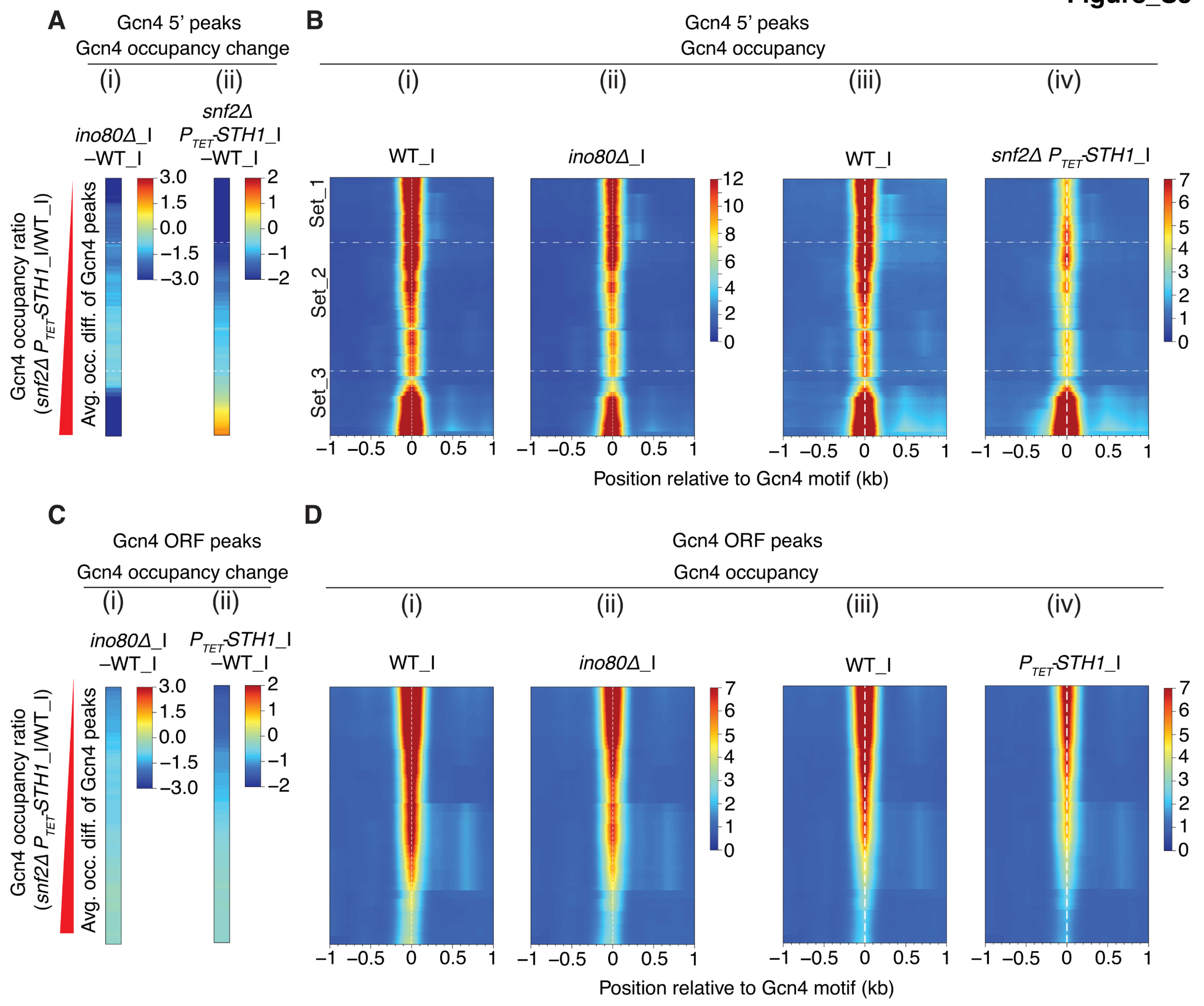
A
Gcn4 5' peaks
Gcn4 occupancy change
(i)
(ii)
ino80Δ_I
–WT_I
snf2Δ
PTET-STH1_I
–WT_I
3.0
1.5
0.0
–1.5
–3.0
2
1
0
–1
–2
Gcn4 occupancy ratio
(snf2Δ PTET-STH1_I/WT_I)
Avg. occ. diff. of Gcn4 peaks
B
Gcn4 5' peaks
Gcn4 occupancy
(i)
(ii)
(iii)
(iv)
WT_I
ino80Δ_I
WT_I
snf2Δ PTET-STH1_I
Set_1
Set_2
Set_3
12
10
8
6
4
2
0
7
6
5
4
3
2
1
0
–1 –0.5 0 0.5 1
Position relative to Gcn4 motif (kb)
C
Gcn4 ORF peaks
Gcn4 occupancy change
(i)
(ii)
ino80Δ_I
–WT_I
PTET-STH1_I
–WT_I
3.0
1.5
0.0
–1.5
–3.0
2
1
0
–1
–2
Gcn4 occupancy ratio
(snf2Δ PTET-STH1_I/WT_I)
Avg. occ. diff. of Gcn4 peaks
D
Gcn4 ORF peaks
Gcn4 occupancy
(i)
(ii)
(iii)
(iv)
WT_I
ino80Δ_I
WT_I
PTET-STH1_I
7
6
5
4
3
2
1
0
–1 –0.5 0 0.5 1
Position relative to Gcn4 motif (kb)

**Figure S3. Differential requirements for Ino80C and RSC for Gcn4 binding at a subset of Gcn4 5' sites. (A)** Heat maps of differences in Gcn4 occupancies averaged across the coordinates of 5' sites between the indicated mutant and WT_I samples for (i) *ino80Δ*_I and (ii) *snf2Δ* $P_{TET}$-*STH1*_I. Gcn4 5' sites were sorted by increasing order of the ratio of Gcn4 occupancies in the double mutant *snf2Δ* $P_{TET}$-*STH1*_I vs. WT_I. **(B)** Heat map depictions of Gcn4 occupancies surrounding the Gcn4 motifs of 5' sites in (i) WT_I (for the *ino80Δ* mutant), (ii) *ino80Δ*_I, (iii) WT_I (for the *snf2Δ* $P_{TET}$-*STH1*_I mutant), and (iv) *snf2Δ* $P_{TET}$-*STH1*_I cells, for the same ordering of 5' sites as in (A). The sets of Gcn4 5' sites (Set_1, Set_2, and Set_3) defined in Fig. 2A-B are depicted in B(i). **(C & D)** Same analyses for *ino80Δ*_I and $P_{TET}$-*STH1*_I as shown in (A-B) except for Gcn4 ORF sites.

**Figure_S4**

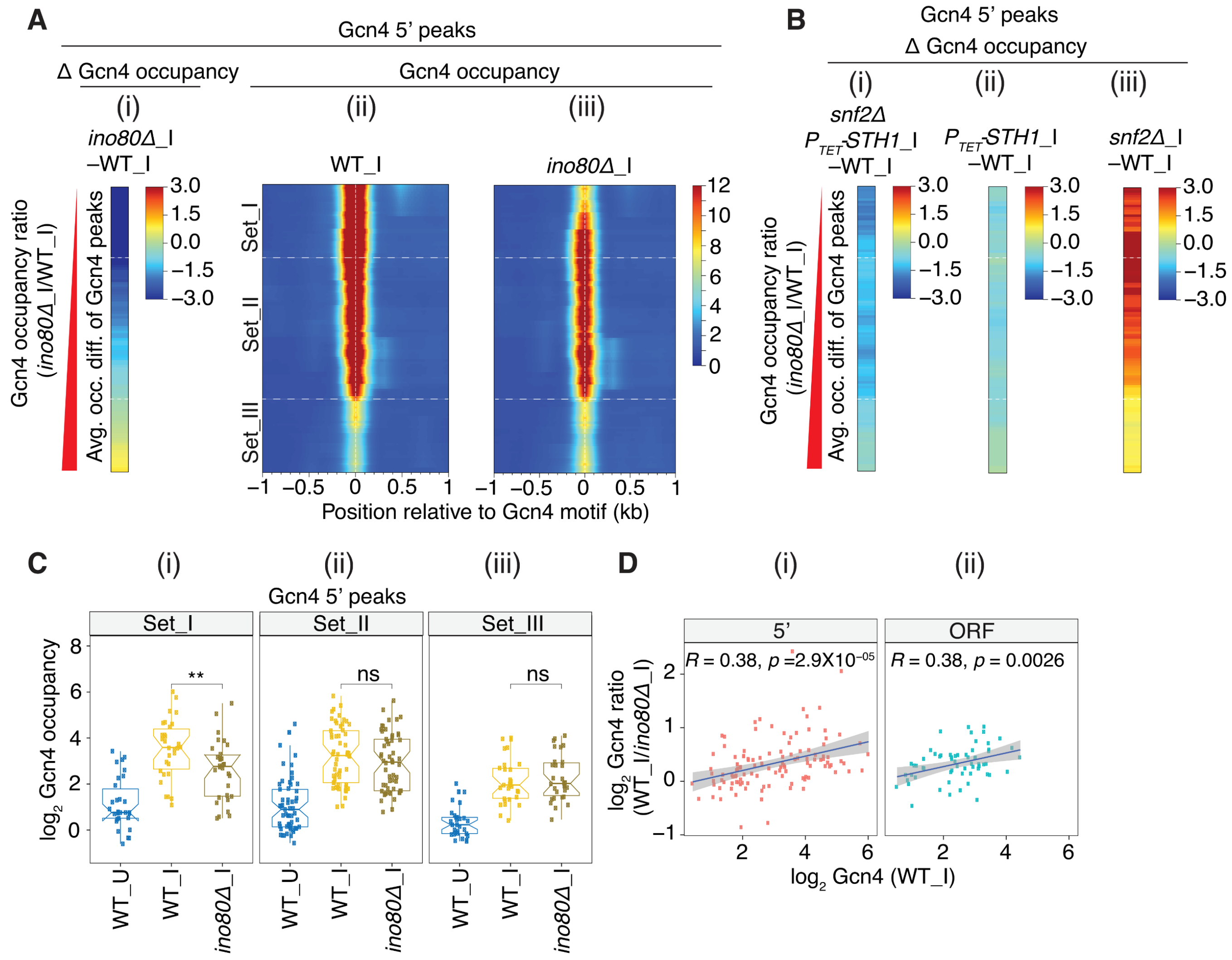

A
Gcn4 5’ peaks
Δ Gcn4 occupancy
Gcn4 occupancy
(i)
(ii)
(iii)
ino80Δ_I
–WT_I
WT_I
ino80Δ_I
Gcn4 occupancy ratio
(ino80Δ_I/WT_I)
Avg. occ. diff. of Gcn4 peaks
Set_I
Set_II
Set_III
Position relative to Gcn4 motif (kb)
B
Gcn4 5’ peaks
Δ Gcn4 occupancy
snf2Δ
P_TET-STH1_I
–WT_I
P_TET-STH1_I
–WT_I
snf2Δ_I
–WT_I
C
Gcn4 5’ peaks
Set_I
Set_II
Set_III
log2 Gcn4 occupancy
**
ns
ns
WT_U
WT_I
ino80Δ_I
D
5’
ORF
R = 0.38, p =2.9X10−05
R = 0.38, p = 0.0026
log2 Gcn4 ratio
(WT_I/ino80Δ_I)
log2 Gcn4 (WT_I)

**Figure S4. Identification of Gcn4 5' sites with heightened Ino80C dependence for Gcn4 occupancy.** (A) Heat map depicting differences in Gcn4 occupancies between *ino80Δ*_I and WT_I cells (i); Gcn4 occupancies surrounding the motifs of 5' sites in (ii) WT_I or (iii) *ino80Δ*_I cells. Gcn4 5' sites were sorted by increasing order of the ratio of Gcn4 occupancies in *ino80Δ*_I vs. WT_I cells, and the first (Set_I, n=30), middle two (Set_II, n=57) and fourth (Set_III, n=30) quartiles of fold-changes are depicted in A(ii). (B) Heat map depictions of Gcn4 occupancies between (i) *snf2Δ* $P_{TET}$*-STH1*_I and WT_I, (ii) $P_{TET}$*-STH1*_I and WT_I, and (iii) *snf2*_I vs. WT_I cells. (C) Notched box plots of $log_2$ Gcn4 occupancy in WT_U, WT_I, and *ino80Δ*_I cells in 3 sets of Gcn4 5' sites comprised of the (i) first (Set_I, n=30), (ii) middle two (Set_II, n=57) and (iii) last (Set_III, n=30) quartiles of the fold-changes in Gcn4 occupancy in *ino80Δ_I* vs. WT_I cells as defined in panel A(ii). *P* values for the significance of differences in medians calculated by the Mann-Whitney-Wilcoxon test are indicated. (D) Scatterplots of $log_2$ ratios of Gcn4 occupancy changes in WT_I vs. *ino80Δ*_I cells plotted against $log_2$ Gcn4 occupancies in WT_I cells for 5' (i) and ORF (ii) Gcn4 sites. Pearson correlation coefficients (*R*) and associated *p* values are indicated.

### Figure_S5

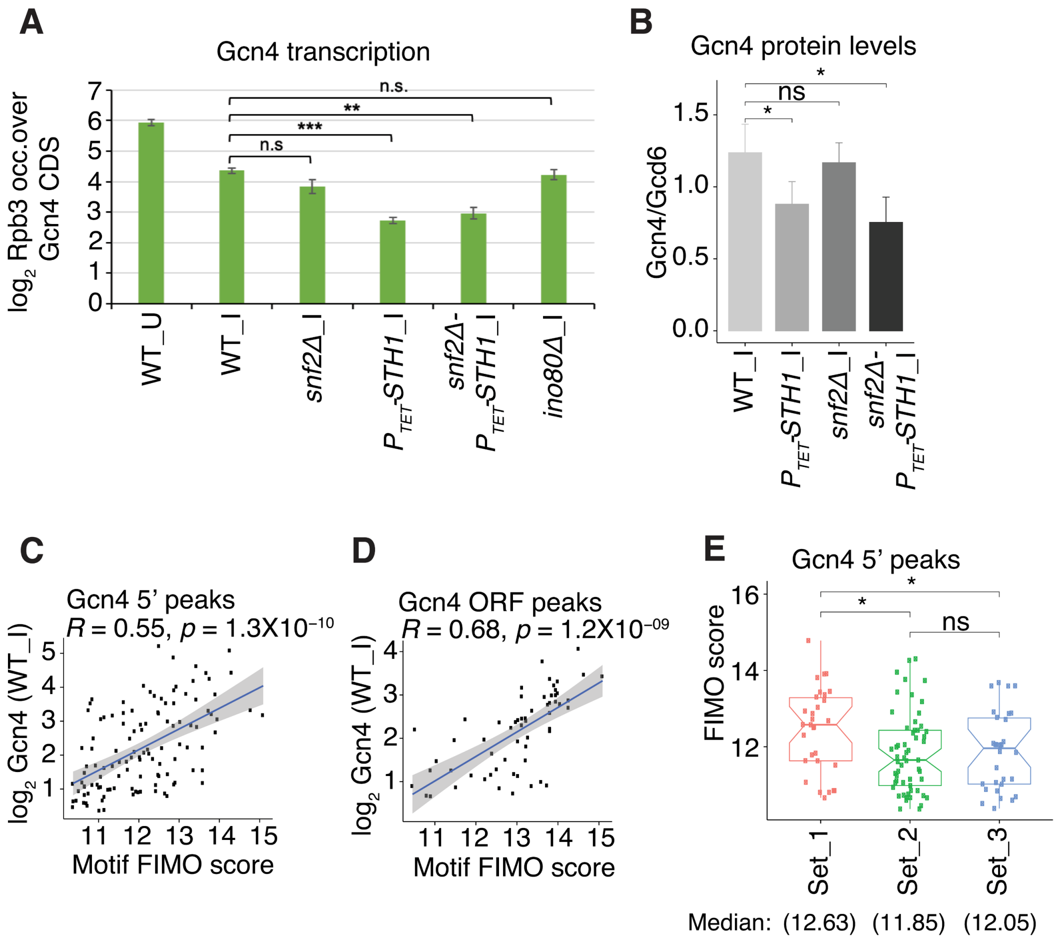

A
Gcn4 transcription
log2 Rpb3 occ.over Gcn4 CDS
n.s.
**
***
n.s
WT_U
WT_I
snf2Δ_I
PTET-STH1_I
snf2Δ-PTET-STH1_I
ino80Δ_I
B
Gcn4 protein levels
Gcn4/Gcd6
*
ns
*
WT_I
PTET-STH1_I
snf2Δ_I
snf2Δ-PTET-STH1_I
C
Gcn4 5' peaks
R = 0.55, p = 1.3X10−10
log2 Gcn4 (WT_I)
Motif FIMO score
D
Gcn4 ORF peaks
R = 0.68, p = 1.2X10−09
log2 Gcn4 (WT_I)
Motif FIMO score
E
Gcn4 5' peaks
FIMO score
*
*
ns
Set_1
Set_2
Set_3
Median: (12.63) (11.85) (12.05)

**Figure S5. Supporting evidence that reduced Gcn4 binding in mutants depleted of RSC or Ino80C occurs preferentially at motifs of highest affinity or accessibility in chromatin in WT_I cells. (A & B)** Histograms depicting (A) $log_2$ Rpb3 occupancies in the *GCN4* CDS measured by Rpb3 ChIP-seq, indicating transcription levels; and (B) Gcn4 protein levels measured previously (Rawal et al. 2018a) by Western blot analysis in the indicated strains using Gcd6 signals analyzed in parallel as loading control. Band intensities for Gcn4 were normalized to those for Gcd6 in the same samples and the mean Gcn4/Gcd6 ratios determined from 3 biological replicates were plotted. Significance of differences in mean values was calculated with the student's t test. **(C & D)** Scatterplots of $log_2$ WT_I Gcn4 occupancies vs. motif FIMO scores (Rawal et al. 2018b) for (C) the 115 Gcn4 5' sites and (D) the 62 Gcn4 ORF peaks, using the motif of highest score for peaks with multiple motifs. Pearson correlation coefficients ($R$) and associated $p$ values are indicated. **(E)** Notched box pots of motif FIMO scores for the sets of Gcn4 5' sites binned according to the fold-changes in occupancy in *snf2Δ* $P_{TET}$-*STH1*_I vs. WT_I cells, as depicted in Fig. 2B.

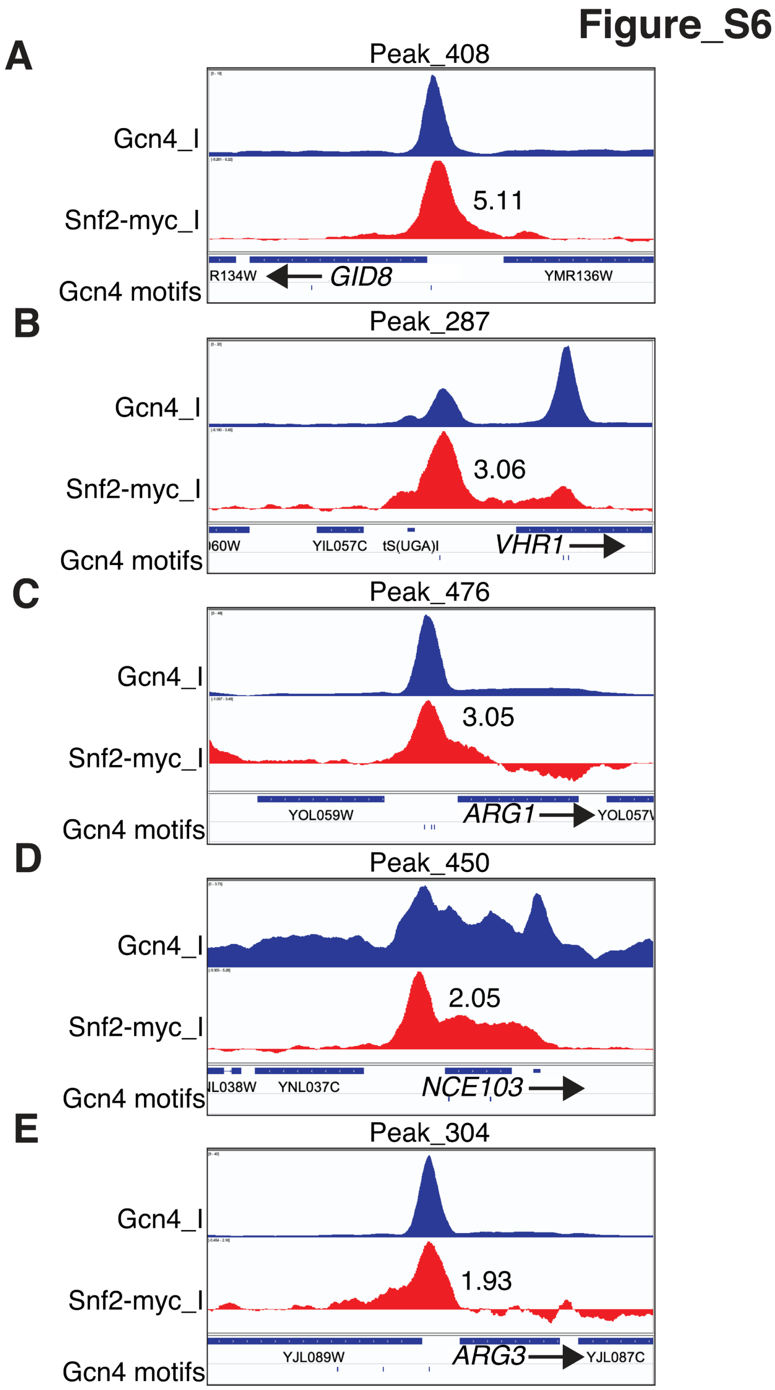


**Figure S6. Gene browser profiles of Gcn4 and corrected Snf2-myc occupancies from ChIP-seq analyses of sonicated chromatin in SM induced WT and *SNF2-myc* cells for selected genes.** The Gcn4 peak numbering assigned previously (Rawal et al. 2018b) is given at the top of each profile, and corrected Snf2-myc occupancies per nucleotide within ±100 bp windows surrounding the Gcn4 motifs are listed next to each peak.

#### Figure_S7

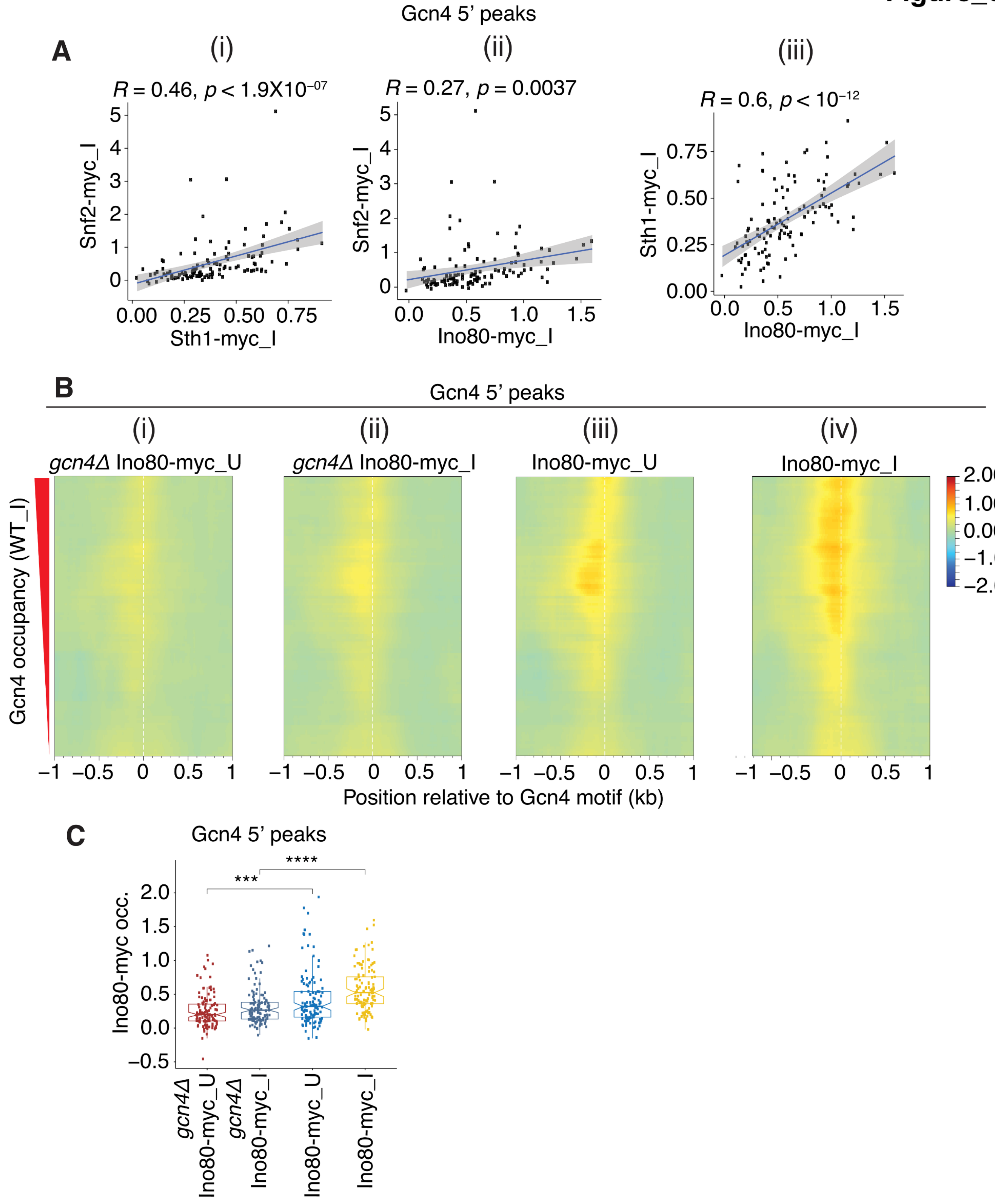
Gcn4 5' peaks
A
(i)
R = 0.46, p < 1.9X10^-07
Snf2-myc_I
Sth1-myc_I
(ii)
R = 0.27, p = 0.0037
Snf2-myc_I
Ino80-myc_I
(iii)
R = 0.6, p < 10^-12
Sth1-myc_I
Ino80-myc_I
B
Gcn4 5' peaks
(i)
gcn4Δ Ino80-myc_U
(ii)
gcn4Δ Ino80-myc_I
(iii)
Ino80-myc_U
(iv)
Ino80-myc_I
Gcn4 occupancy (WT_I)
2.00
1.00
0.00
−1.00
−2.00
Position relative to Gcn4 motif (kb)
C
Gcn4 5' peaks
Ino80-myc occ.
***
****
gcn4Δ Ino80-myc_U
gcn4Δ Ino80-myc_I
Ino80-myc_U
Ino80-myc_I

**Figure S7. Supporting evidence for recruitment of the three CRs by Gcn4 to its 5' sites.** (**A) (i)-(iii)** Scatterplots of corrected occupancies of the indicated myc-tagged CR subunits in WT_I cells measured within ±100 bp windows surrounding the Gcn4 motifs of 5' Gcn4 peaks. Pearson correlation coefficients (*R*) and associated *p* values are indicated. **(B)** Heat map depictions of corrected Ino80-myc occupancies at the 5' Gcn4 peaks, sorted by decreasing Gcn4 occupancies in WT_I cells and plotted relative to the Gcn4 motifs, for (i) uninduced *gcn4Δ* cells, (ii) SM-treated *gcn4Δ* cells, (iii) uninduced WT cells, and (iv) SM-treated WT cells. Occupancies were calculated from ChIP-seq data of mildly sonicated chromatin from 2 or 3 biological replicates each of isogenic *GCN4* or *gcn4Δ* strains, harboring *INO80-myc* or untagged *INO80,* under inducing or uninducing conditions, correcting the occupancies for *INO80-myc* cells for those measured for the untagged *INO80* cells of the same *GCN4* genotype and growth conditions. **(C)** Notched box plots of corrected Ino80-myc occupancies per nucleotide within ±100 bp windows surrounding the Gcn4 motifs of 5' sites in uninduced (_U) or SM induced (_I) *gcn4Δ* or WT cells.

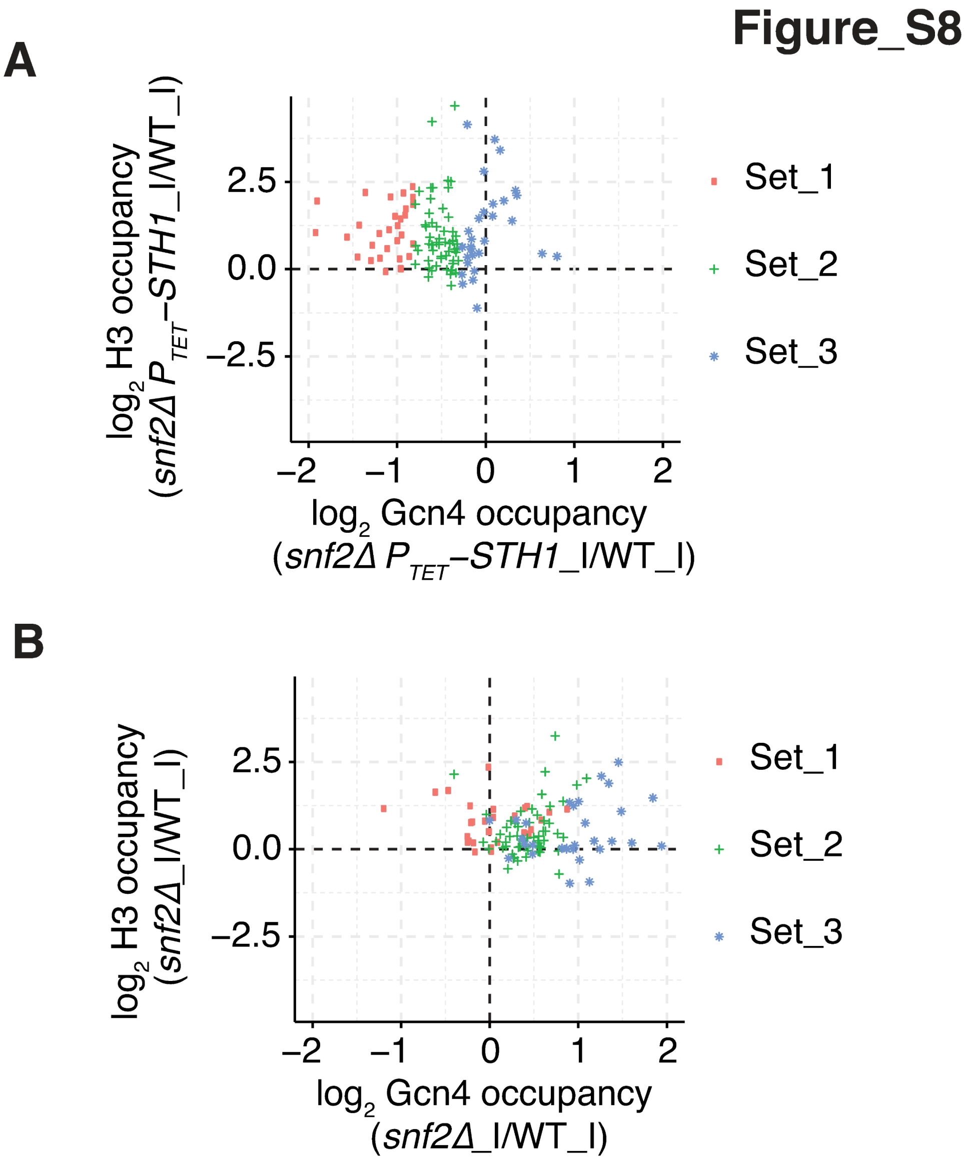


**Figure S8. Supporting evidence for defective eviction of nucleosomes surrounding 5' Gcn4 motifs in SWI/SNF and RSC mutants.** Sectored scatterplot of the $log_2$ ratios of Gcn4 occupancies plotted against the $log_2$ ratios of H3 occupancies per base pair in the ±100 bp windows surrounding the Gcn4 motifs in (A) *snf2Δ* $P_{TET}$*-STH1*_I vs. WT_I or (B) *snf2Δ*_I vs. WT_I cells. The data points for the Gcn4 5' sites defined in Fig. 2B(i) are depicted as follows: Set_1, red rectangles, Set_2, green pluses, and Set_3, blue stars.

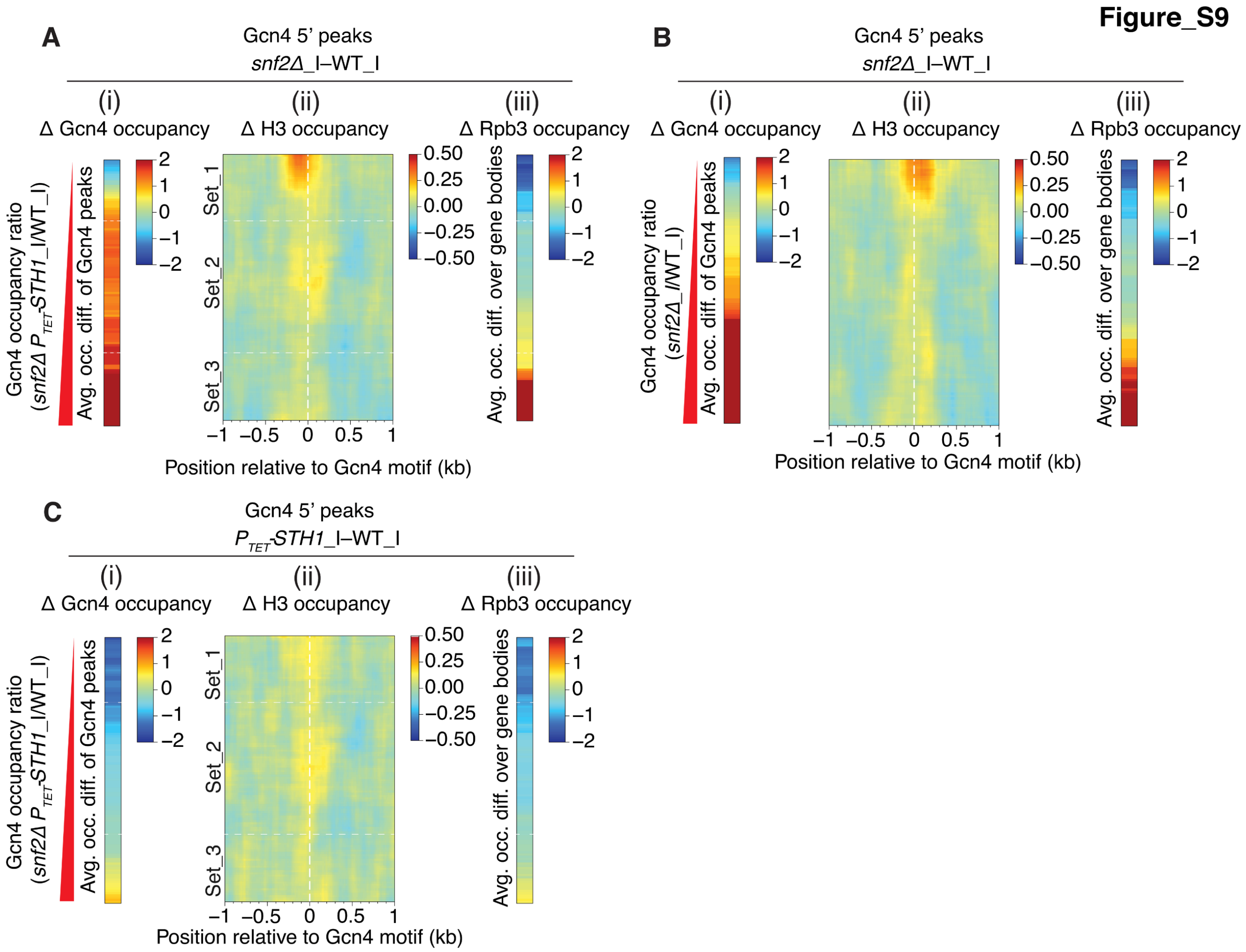


**Figure S9. Defective eviction of nucleosomes associated with reduced Gcn4 occupancies at a subset of 5' Gcn4 peaks in *snf2Δ*_I cells. (A)** (i)-(iii) Heat maps depicting differences between *snf2Δ*_I and WT_I cells for (i) Gcn4 occupancies measured as in Fig 2A(iii), (ii) H3 occupancies surrounding the Gcn4 motifs of 5' sites from H3 ChIP-seq data, and (iii) Rpb3 occupancies averaged over the CDS of 5' genes, for the Gcn4 5' sites sorted by increasing order of fold-changes in Gcn4 occupancies in *snf2Δ* $P_{TET}$-*STH1*_I vs. WT_I cells. **(B)** (i)-(iii) Same analyses shown in (A) except sorted by increasing order of fold-changes in Gcn4 occupancies in *snf2Δ*_I vs. WT_I cells. **(C)** (i)-(iii) Same analyses shown in (A) except for $P_{TET}$-*STH1*_I vs. WT_I data.

**Figure_S10**

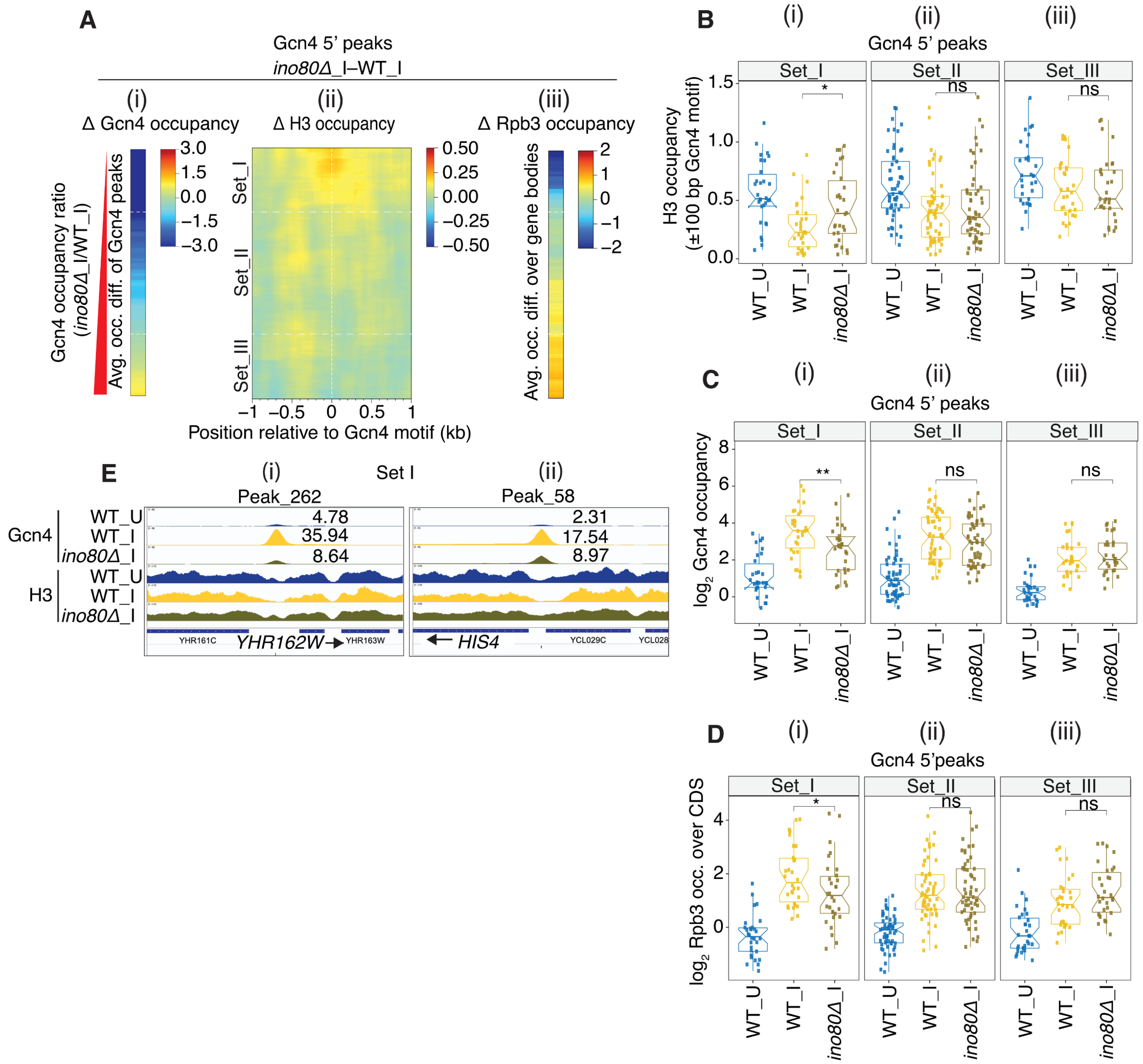
A
Gcn4 5' peaks
ino80Δ_I–WT_I
(i) Δ Gcn4 occupancy
(ii) Δ H3 occupancy
(iii) Δ Rpb3 occupancy
Gcn4 occupancy ratio (ino80Δ_I/WT_I)
Avg. occ. diff. of Gcn4 peaks
Set_I
Set_II
Set_III
Position relative to Gcn4 motif (kb)
Avg. occ. diff. over gene bodies
B
Gcn4 5' peaks
H3 occupancy (±100 bp Gcn4 motif)
C
Gcn4 5' peaks
log2 Gcn4 occupancy
D
Gcn4 5'peaks
log2 Rpb3 occ. over CDS
WT_U
WT_I
ino80Δ_I
ns
E
Set I
(i) Peak_262
(ii) Peak_58
Gcn4
H3
4.78
35.94
8.64
2.31
17.54
8.97
YHR161C
YHR162W
YHR163W
HIS4
YCL029C
YCL028

**Figure S10. Defective eviction of nucleosomes associated with reduced Gcn4 occupancies at a subset of 5' Gcn4 peaks in *ino80Δ*_I cells. (A)** (i)-(iii) Heat maps depicting differences between *ino80Δ*_I and WT_I cells for (i) Gcn4 occupancies measured as in Fig S4A(i), (ii) H3 occupancies surrounding the Gcn4 motifs of 5' sites from H3 ChIP-seq data, and (iii) Rpb3 occupancies averaged over the CDS of 5' genes, for Gcn4 5' sites sorted by increasing order of fold-changes in Gcn4 occupancies in *ino80Δ*_I vs. WT_I cells. **(B-D)** Notched box plots for the 3 sets of 5' sites defined in Fig. S4A(ii), and indicated again in panel A(ii), depicting (B) H3 occupancies per base pair in the ±100 bp windows surrounding the Gcn4 motifs, (C) $\log_2$ Gcn4 occupancies taken from Fig. S4C, and (D) $\log_2$ Rpb3 occupancies averaged over the CDS of genes with 5' sites. H3 and Rpb3 occupancies were calculated from ChIP-seq data of sonicated chromatin from at least 3 biological replicates of WT_U, WT_I and *ino80Δ*_I cells. *P* values from Mann-Whitney-Wilcoxon tests are indicated. The heat map of H3 occupancy changes conferred by *ino80Δ* around the 5' motifs ordered by the Gcn4 occupancy reductions in this mutant (panel A(i)) reveals that the 5' sites with the strongest reductions in Gcn4 binding in *ino80Δ* cells located at the top of the map (Set_I) show the strongest increases in H3 occupancies centered around the Gcn4 motifs (panel A(ii)). Moreover, the decreases in median Gcn4 occupancy for individual 5' sites conferred by *ino80Δ* are paralleled by increased median H3 occupancies for the Set_I group of 5' sites; whereas the sites in Set_II and III show no significant changes in median H3 or Gcn4 occupancies in *ino80Δ*_I versus WT_I cells (panels 10B-C, Sets_I-III, col. 3 vs. 2). The subset of genes with 5' sites that are most dependent on Ino80C for Gcn4 binding generally show the greatest reductions in Rpb3 occupancies (Set_I sites in panel A (iii) vs. (i)). Moreover, the Set_I genes, but not genes in Sets_II-III, show reduced median occupancies of both Gcn4 and Rpb3 in *ino80Δ*_I versus WT_I cells (panels C-D(i)-(iii), col. 3 vs. 2). As only the Set_I genes also exhibit increased median H3 occupancies in *ino80Δ*_I cells (panel B(i)-(iii), col. 3 vs. 2), it seems likely that a defect in Gcn4 binding (and subsequent impaired recruitment of other coactivators) in combination with loss of Ino80C-mediated promoter nucleosome eviction produces the reduced transcription of Set_I genes conferred by *ino80Δ*. **(E)** Gene browser profiles of Gcn4 and H3 occupancies from ChIP-seq analyses of sonicated chromatin for the indicated strains, as described in Fig. S2.

Figure_S11

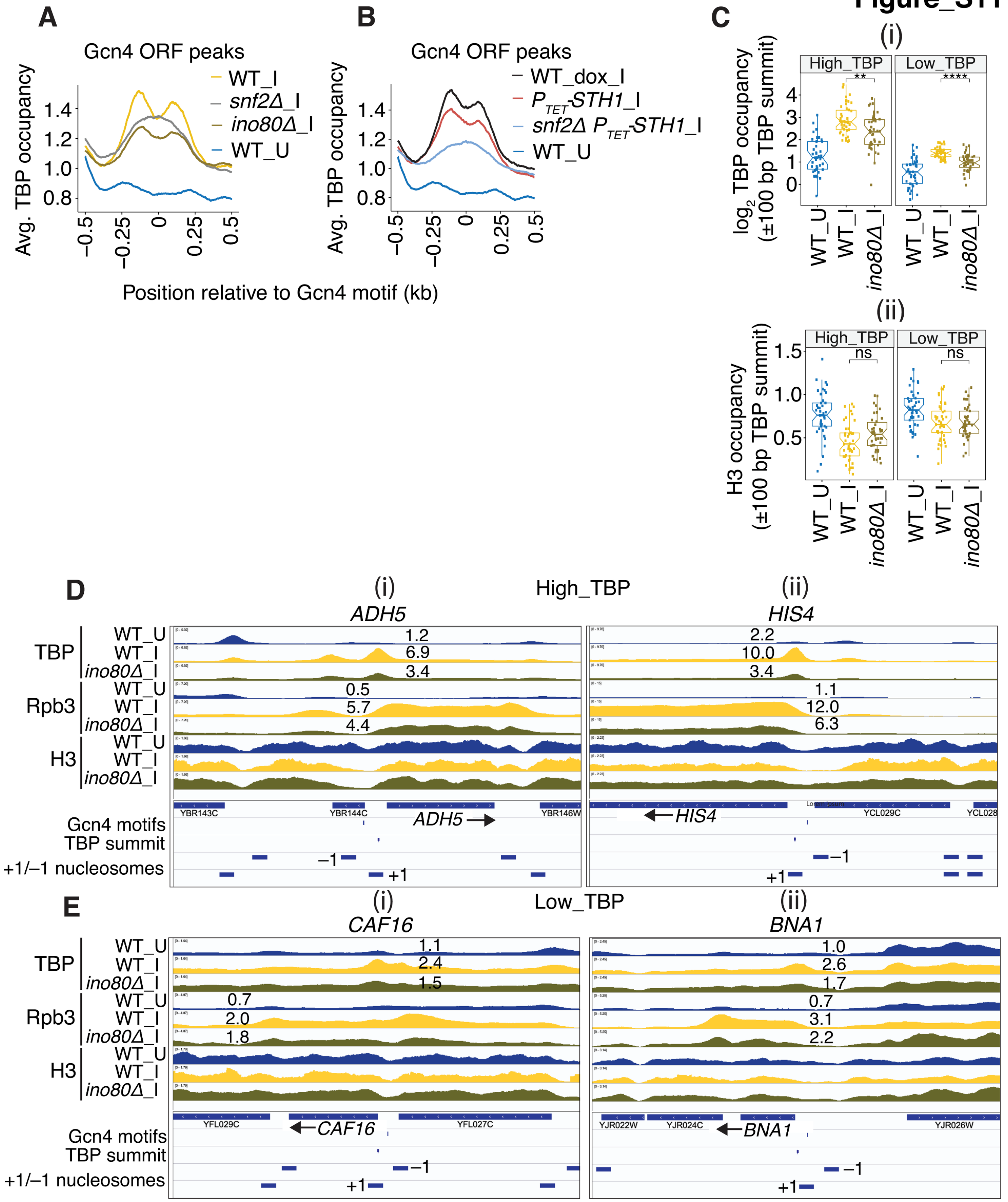
A
Gcn4 ORF peaks
WT_I
snf2Δ_I
ino80Δ_I
WT_U
Avg. TBP occupancy
B
WT_dox_I
P_TET-STH1_I
snf2Δ P_TET-STH1_I
Position relative to Gcn4 motif (kb)
C
(i)
High_TBP
Low_TBP
log2 TBP occupancy (±100 bp TBP summit)
(ii)
ns
H3 occupancy (±100 bp TBP summit)
D
High_TBP
ADH5
HIS4
TBP
Rpb3
H3
Gcn4 motifs
TBP summit
+1/−1 nucleosomes
E
Low_TBP
CAF16
BNA1

**Figure S11. Decreased TBP recruitment at 5' and ORF genes in *ino80Δ*_I cells. (A-B)** Averaged TBP occupancies surrounding the Gcn4 motifs in ORF peaks from ChIP-seq data of sonicated chromatin using anti-TBP antibodies from at least 2 biological replicates for the indicated mutant and WT strains. **(C)** Notched box plots of factor occupancies in the 83 5' gene promoters for the two bins defined in Fig. 5C for (i) $log_2$ TBP within ±100 bp of TBP peak summits and (ii) H3 within ±100 bp of the TBP peak summits. *P* values from Mann-Whitney-Wilcoxon tests are indicated. **(D-E)** Gene browser profiles of TBP, Rpb3, and H3 occupancies from ChIP-seq analyses of sonicated chromatin for the indicated strains. The TBP occupancies per nucleotide over the TBP peaks and Rpb3 occupancies per nucleotide over the CDS are listed next to the relevant peaks or CDSs. The locations of the Gcn4 motifs and TBP summits are indicated with vertical hash marks, and the positions of -1 and +1 nucleosomes with dashes, at the bottom of each profile.

**Figure_S12**

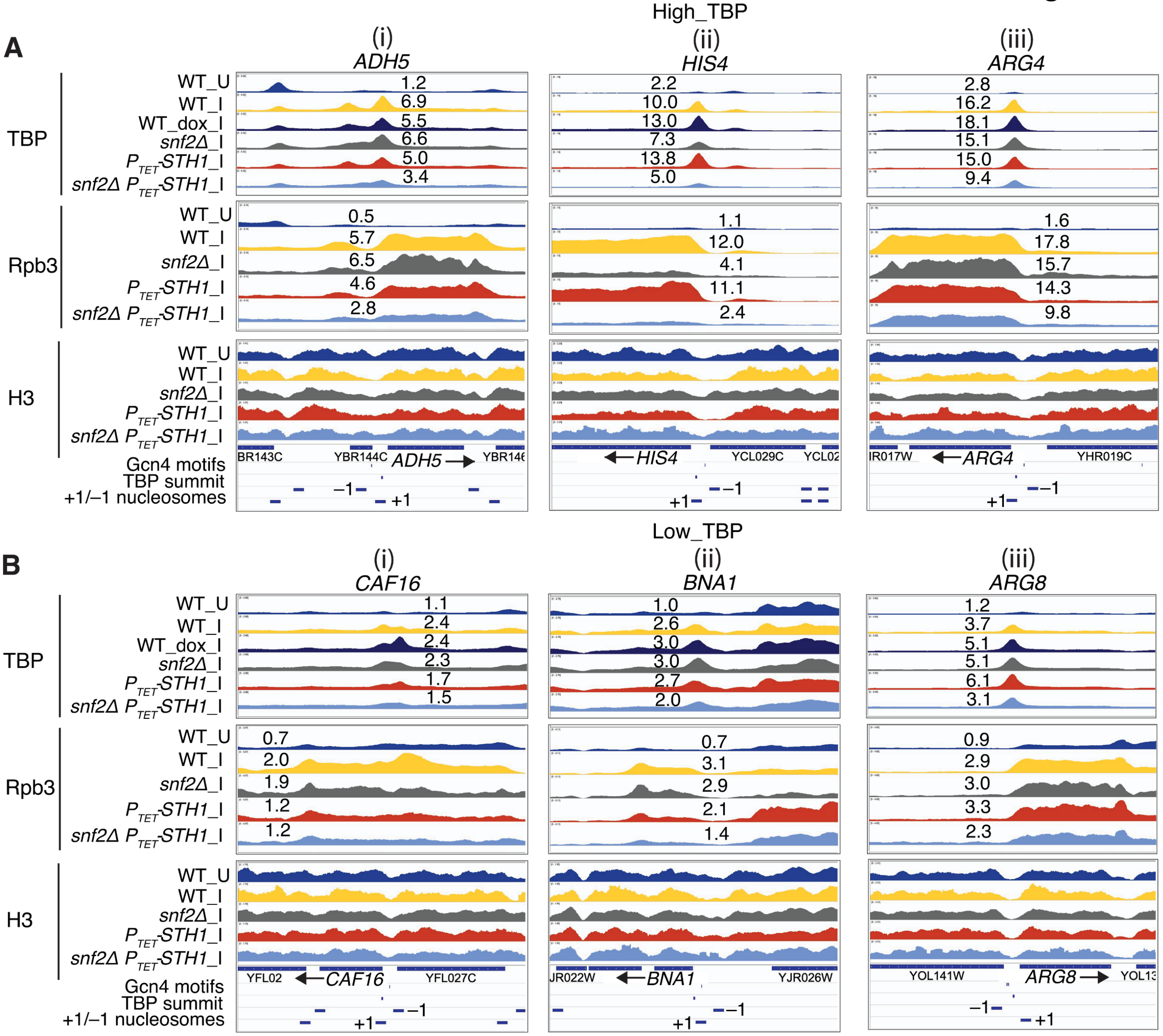


**Figure S12. Gene browser profiles of TBP, Rpb3, and H3 occupancies from ChIP-seq analyses of sonicated chromatin for the indicated strains.** The TBP occupancies per nucleotide over the TBP peaks and Rpb3 occupancies per nucleotide over the CDS are listed next to the relevant peaks or CDSs. The locations of the Gcn4 motifs and TBP summits are indicated with vertical hash marks, and the positions of -1 and +1 nucleosomes with dashes, at the bottom of each profile.

#### Figure_S13

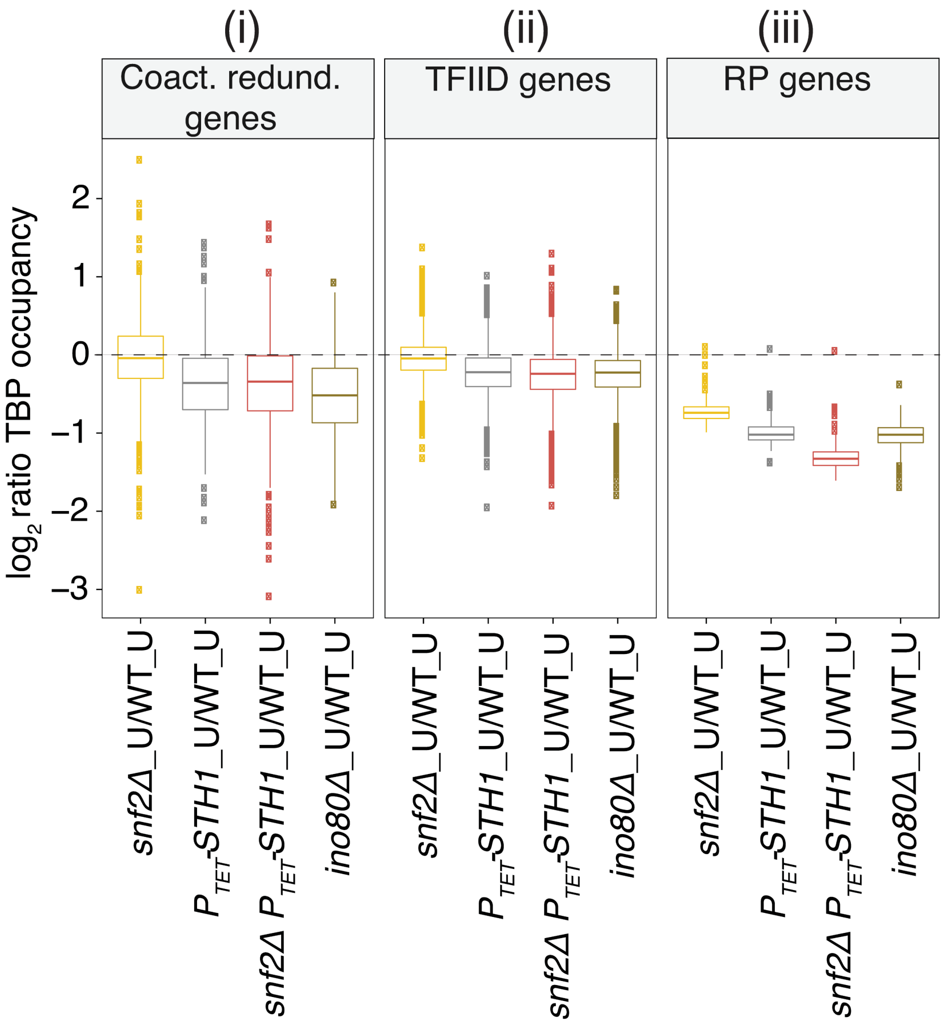


**Figure S13. Coactivator-redundant genes have a greater requirement than TFIID-dependent genes for RSC and Ino80C for TBP recruitment.** Changes in TBP occupancies surrounding the TSSs in the indicated mutants versus WT under non-starvation conditions are plotted for the coactivator-redundant and TFIID-dependent genes defined by Donczew et al. (2020), along with the corresponding changes for the RPGs.

**Table S1: Yeast strains used in this study**

| Name | Parent | Genotype | Reference |
|---|---|---|---|
| F729/BY4741 | NA | *MATa his3Δ1 leu2Δ0 met15Δ0 ura3Δ0* | Research genetics |
| F731 | BY4741 | *MATa his3Δ1 leu2Δ0 met15Δ0 ura3Δ0 gcn4Δ::kanMX4* | Research genetics |
| F748 | BY4741 | *MATa his3Δ1 leu2Δ0 met15Δ0 ura3Δ0 snf2Δ::kanMX4* | Research genetics |
| HQY1632 | BY4741 | *MATa his3Δ1 leu2Δ0 met15Δ0 ura3Δ0 HIS3MX6::*$P_{TET}$*STH1* | (Rawal et al. 2018) |
| HQY1660 | F748 | *MATa his3Δ1 leu2Δ0 met15Δ0 ura3Δ0 snf2Δ::kanMX4 HIS3MX6::*$P_{TET}$*STH1* | (Rawal et al. 2018) |
| YR092 | Y24517 | *MATa his3Δ1 leu2Δ0 met15Δ0 ura3Δ0 ino80Δ::kanMX4* | (Qiu et al. 2020) |
| HQY367 | BY4741 | *MATa his3Δ1 leu2Δ0 met15Δ0 ura3Δ0 SNF2-myc13::HIS3MX6* | (Swanson et al. 2003) |
| HQY459 | BY4741 | *MATa his3Δ1 leu2Δ0 met15Δ0 ura3Δ0 STH1-myc13::HIS3MX6* | (Swanson et al. 2003) |
| HQY1687 | BY4741 | *MATa his3Δ1 leu2Δ met15Δ ura3Δ INO80::myc13::HIS3MX6* | (Qiu et al. 2020) |
| HQY1718 | F731 | *MATa his3Δ1 leu2Δ0 met15Δ0 ura3Δ0 gcn4Δ::kanMX4 INO80::myc13::HIS3MX6* | This study |

*HIS3** designates the *HIS3* allele from *S. kluyveri*

#### Table S2: Compilation of ChIP-seq replicate experiments

##### I. Gcn4 ChIP-seq

| Strain | IP | Sample ID | All PE reads | PE rmdup | Pearson correlation between replicates | | | Source |
|---|---|---|---|---|---|---|---|---|
| | | | | | AGH32_HQ46 | AGH32_HQ47 | | |
| WT_U | Gcn4 | AGH32 _HQ45 | 9434957 | 834910 | 0.997 | 0.997 | | (Rawal, Chereji, Valabhoju, et al. 2018) |
| WT_U | Gcn4 | AGH32 _HQ46 | 8293362 | 352621 | | 0.996 | | (Rawal, Chereji, Valabhoju, et al. 2018) |
| WT_U | Gcn4 | AGH32 _HQ47 | 6312086 | 223598 | | | | (Rawal, Chereji, Valabhoju, et al. 2018) |
| | | | | | AGH95_10 | AGH66_01 | AGH66_02 | |
| WT_I | Gcn4 | AGH95_09 | 23969984 | 18446498 | 0.998 | 0.973 | 0.967 | This study |
| WT_I | Gcn4 | AGH95_10 | 34793241 | 25793908 | | 0.972 | 0.970 | This study |
| WT_I | Gcn4 | AGH66_01 | 13001067 | 3812850 | | | 0.992 | This study |
| WT_I | Gcn4 | AGH66_02 | 21568439 | 8750863 | | | | This study |
| | | | | | AGH95_12 | | | |
| *snf2Δ*_I | Gcn4 | AGH95_11 | 16513767 | 12929335 | 0.998 | | | This study |
| *snf2Δ*_I | Gcn4 | AGH95_12 | 25382449 | 19395455 | | | | This study |
| | | | | | AGH96_10 | AGH66_03 | AGH66_04 | |
| $P_{TET}$*STH1*_I | Gcn4 | AGH96_09 | 23383351 | 17959152 | 0.990 | 0.982 | 0.980 | This study |
| $P_{TET}$*STH1*_I | Gcn4 | AGH96_10 | 28009712 | 20396793 | | 0.969 | 0.969 | This study |
| $P_{TET}$*STH1*_I | Gcn4 | AGH66_03 | 13744350 | 6106482 | | | 0.993 | This study |
| $P_{TET}$*STH1*_I | Gcn4 | AGH66_04 | 17305536 | 7027543 | | | | This study |
| | | | | | AGH96_12 | AGH66_05 | AGH66_06 | |
| *snf2Δ* $P_{TET}$*STH1*_I | Gcn4 | AGH96_11 | 35827719 | 27272344 | 0.981 | 0.972 | 0.972 | This study |
| *snf2Δ* $P_{TET}$*STH1*_I | Gcn4 | AGH96_12 | 31537097 | 23712138 | | 0.949 | 0.952 | This study |
| *snf2Δ* $P_{TET}$*STH1*_I | Gcn4 | AGH66_05 | 23853729 | 8885029 | | | 0.997 | This study |
| *snf2Δ* $P_{TET}$*STH1*_I | Gcn4 | AGH66_06 | 22769050 | 9213922 | | | | This study |
| | | | | | AGH113_08 | | | |
| WT_U | Gcn4 | AGH113_07 | 10542959 | 7090399 | 0.981 | | | This study |
| WT_U | Gcn4 | AGH113_08 | 10513497 | 6863672 | | | | This study |
| | | | | | AGH113_10 | | | |
| WT_I | Gcn4 | AGH113_09 | 13698040 | 9265216 | 0.991 | | | This study |
| WT_I | Gcn4 | AGH113_10 | 41560943 | 28746522 | | | | This study |
| | | | | | AGH113_14 | | | |
| *ino80Δ*_I | Gcn4 | AGH113_13 | 19654864 | 15014306 | 0.998 | | | This study |
| *ino80Δ*_I | Gcn4 | AGH113_14 | 19210671 | 14099687 | | | | This study |

#### Table S2 (cont'd):

##### II. TBP ChIP-seq

| Strain | IP | Sample ID | All PE reads | PE rmdup | Pearson correlation between replicates | | Source |
|---|---|---|---|---|---|---|---|
| | | | | | AGH121-2 | AGH121-3 | (Qiu et al. 2020) |
| WT_U | TBP | AGH121-1 | 16,885,453 | 10,236,588 | 0.999 | 0.999 | (Qiu et al. 2020) |
| WT_U | TBP | AGH121-2 | 27,362,033 | 12,530,926 | | 0.999 | (Qiu et al. 2020) |
| WT_U | TBP | AGH121-3 | 22,937,418 | 11,812,400 | | | (Qiu et al. 2020) |
| | | | | | AGH121-5 | | (Qiu et al. 2020) |
| WT_I | TBP | AGH121-4 | 24,785,155 | 14,448,990 | 0.996 | | (Qiu et al. 2020) |
| WT_I | TBP | AGH121-5 | 4,584,274 | 3,832,277 | | | (Qiu et al. 2020) |
| | | | | | AGH121-8 | AGH121-9 | (Qiu et al. 2020) |
| *ino80Δ* _U | TBP | AGH121-7 | 26,930,283 | 18,269,275 | 0.998 | 0.999 | (Qiu et al. 2020) |
| *ino80Δ* _U | TBP | AGH121-8 | 15,095,156 | 10,094,991 | | 0.999 | (Qiu et al. 2020) |
| *ino80Δ* _U | TBP | AGH121-9 | 22,489,806 | 12,887,915 | | | (Qiu et al. 2020) |
| | | | | | AGH121-11 | AGH121-12 | (Qiu et al. 2020) |
| *ino80Δ* _I | TBP | AGH121-10 | 27,763,987 | 17,051,822 | 0.999 | 0.999 | (Qiu et al. 2020) |
| *ino80Δ* _I | TBP | AGH121-11 | 27,142,493 | 18,528,771 | | 0.999 | (Qiu et al. 2020) |
| *ino80Δ* _I | TBP | AGH121-12 | 26,288,826 | 17,279,728 | | | (Qiu et al. 2020) |
| | | | | | AGH142_14 | | |
| WT_I_dox | TBP | AGH142_13 | 27411329 | 14710509 | 0.998 | | This study |
| WT_I_dox | TBP | AGH142_14 | 31115073 | 13579121 | | | This study |
| | | | | | AGH142_6 | | |
| *snf2Δ*_U | TBP | AGH142_5 | 23221991 | 12712910 | 0.999 | | This study |
| *snf2Δ*_U | TBP | AGH142_6 | 29278698 | 14179344 | | | This study |
| | | | | | AGH142_8 | | |
| *snf2Δ*_I | TBP | AGH142_7 | 26652341 | 14607284 | 0.998 | | This study |
| *snf2Δ*_I | TBP | AGH142_8 | 25661082 | 16282120 | | | This study |
| | | | | | AGH142_10 | | |
| $P_{TET}$*STH1*_U_dox | TBP | AGH142_9 | 24829710 | 12561822 | 0.999 | | This study |
| $P_{TET}$*STH1*_U_dox | TBP | AGH142_10 | 28314060 | 12766273 | | | This study |
| | | | | | AGH142_12 | | |
| $P_{TET}$*STH1*_I_dox | TBP | AGH142_11 | 25442841 | 13057814 | 0.999 | | This study |
| $P_{TET}$*STH1*_I_dox | TBP | AGH142_12 | 25302510 | 12058412 | | | This study |
| | | | | | AGH142_2 | | |
| *snf2Δ* $P_{TET}$*STH1*_U_dox | TBP | AGH142_1 | 23,732,740 | 13,064,587 | 0.999 | | This study |
| *snf2Δ* $P_{TET}$*STH1*_U_dox | TBP | AGH142_2 | 24,739,855 | 13,218,689 | | | This study |
| | | | | | AGH142_4 | | |
| *snf2Δ* $P_{TET}$*STH1*_I_dox | TBP | AGH142_3 | 23905850 | 15330844 | 0.999 | | This study |
| *snf2Δ* $P_{TET}$*STH1*_I_dox | TBP | AGH142_4 | 28963188 | 16277896 | | | This study |

#### Table S2 (cont'd):

##### III. CR-myc ChIP-seq

| Strain | IP | Sample ID | All PE reads | PE rmdup | Pearson correlation between replicates | | Source |
|---|---|---|---|---|---|---|---|
| | | | | | AGH79_10 | | |
| *SNF2-myc*_U | myc | AGH79_09 | 14233915 | 11112228 | 0.993 | | This study |
| *SNF2-myc*_U | myc | AGH79_10 | 24867367 | 17476084 | | | This study |
| | | | | | AGH79_12 | | |
| *SNF2-myc*_I | myc | AGH79_11 | 13822144 | 10588610 | 0.967 | | This study |
| *SNF2-myc*_I | myc | AGH79_12 | 17179565 | 12644430 | | | This study |
| | | | | | AGH94_10 | | |
| *STH1-myc*_U | myc | AGH94_09 | 17209512 | 12751019 | 0.994 | | This study |
| *STH1-myc*_U | myc | AGH94_10 | 21946759 | 14900926 | | | This study |
| | | | | | AGH94_12 | | |
| *STH1-myc*_I | myc | AGH94_11 | 31459556 | 21973822 | 0.948 | | This study |
| *STH1-myc*_I | myc | AGH94_12 | 19548461 | 13093796 | | | This study |
| | | | | | AGH144_2 | | |
| WT_U | myc | AGH144_1 | 3072942 | 2120150 | 0.992 | | This study |
| WT_U | myc | AGH144_2 | 9738163 | 6130932 | | | This study |
| | | | | | AGH144_4 | | |
| WT_I | myc | AGH144_3 | 11268530 | 7234492 | 0.972 | | This study |
| WT_I | myc | AGH144_4 | 1765955 | 1301154 | | | This study |
| | | | | | AGH101_18 | | |
| WT_I | myc | AGH101_17 | 14883149 | 12054753 | 0.991 | | (Qiu et al. 2020) |
| WT_I | myc | AGH101_18 | 11930745 | 9789686 | | | (Qiu et al. 2020) |
| | | | | | AGH144_6 | AGH144_7 | |
| *INO80-myc*_U | myc | AGH144_5 | 20470589 | 14552493 | 0.990 | 0.994 | This study |
| *INO80-myc*_U | myc | AGH144_6 | 25673285 | 17246641 | | 0.992 | This study |
| *INO80-myc*_U | myc | AGH144_7 | 19606771 | 13695074 | | | This study |
| | | | | | AGH144_9 | AGH144_10 | |
| *INO80-myc*_I | myc | AGH144_8 | 13548606 | 9478743 | 0.979 | 0.979 | This study |
| *INO80-myc*_I | myc | AGH144_9 | 21052614 | 14229783 | | 0.991 | This study |
| *INO80-myc*_I | myc | AGH144_10 | 27494331 | 17575016 | | | This study |
| | | | | | AGH144_12 | | |
| *gcn4Δ*_U | myc | AGH144_11 | 16724373 | 10573134 | 0.996 | | This study |
| *gcn4Δ*_U | myc | AGH144_12 | 14598145 | 9133914 | | | This study |
| | | | | | AGH144_14 | | |
| *gcn4Δ*_I | myc | AGH144_13 | 20372363 | 11877487 | 0.987 | | This study |
| *gcn4Δ*_I | myc | AGH144_14 | 21985398 | 13319316 | | | This study |
| | | | | | AGH144_16 | AGH144_17 | |
| *gcn4Δ INO80-myc*_U | myc | AGH144_15 | 23070235 | 15948119 | 0.989 | 0.989 | This study |
| *gcn4Δ INO80-myc*_U | myc | AGH144_16 | 23382112 | 16195621 | | 0.988 | This study |
| *gcn4Δ INO80-myc*_U | myc | AGH144_17 | 25118691 | 17289728 | | | This study |
| | | | | | AGH144_19 | AGH144_20 | |

| | | | | | | | |
|---|---|---|---|---|---|---|---|
| *gcn4Δ INO80-myc*_I | myc | AGH144_18 | 22775944 | 15784565 | 0.946 | 0.980 | This study |
| *gcn4Δ INO80-myc*_I | myc | AGH144_19 | 29498726 | 18524887 | | 0.982 | This study |
| *gcn4Δ INO80-myc*_I | myc | AGH144_20 | 22367225 | 15448406 | | | This study |

#### Table S2 (cont'd):

##### IV. Rpb3 ChIP-seq

| Strain | IP | Sample ID | All PE reads | PE rmdup | Pearson correlation between replicates | | | Source |
|---|---|---|---|---|---|---|---|---|
| | | | | | AGH03_2 | AGH03_3 | | |
| WT_U | Rpb3 | AGH03_1 | 12,103,981 | 7,624,104 | 0.997 | 0.997 | | (Qiu et al. 2016) |
| WT_U | Rpb3 | AGH03_2 | 15,621,065 | 10,271,685 | | 0.994 | | (Qiu et al. 2016) |
| WT_U | Rpb3 | AGH03_3 | 14,624,526 | 8,895,936 | | | | (Qiu et al. 2016) |
| | | | | | AGH101_10 | | | |
| *snf2Δ* _U | Rpb3 | AGH101_09 | 18890173 | 15825662 | 0.999 | | | This study |
| *snf2Δ* _U | Rpb3 | AGH101_10 | 19845532 | 15822921 | | | | This study |
| | | | | | AGH101_12 | | | |
| $P_{TET}$*STH1*_U | Rpb3 | AGH101_11 | 16038655 | 13091284 | 0.999 | | | This study |
| $P_{TET}$*STH1*_U | Rpb3 | AGH101_12 | 16877236 | 13736217 | | | | This study |
| | | | | | AGH101_14 | | | |
| *snf2Δ* $P_{TET}$*STH1*_U | Rpb3 | AGH101_13 | 18258099 | 15112166 | 0.998 | | | This study |
| *snf2Δ* $P_{TET}$*STH1*_U | Rpb3 | AGH101_14 | 23241654 | 18587864 | | | | This study |
| | | | | | AGH64_8 | AGH64_9 | | |
| *ino80Δ* _U | Rpb3 | AGH64_7 | 6602236 | 3364700 | 0.993 | 0.993 | | This study |
| *ino80Δ* _U | Rpb3 | AGH64_8 | 7066940 | 3954861 | | 0.995 | | This study |
| *ino80Δ* _U | Rpb3 | AGH64_9 | 8155385 | 3999284 | | | | This study |
| | | | | | AGH03_5 | AGH03_6 | | |
| WT_I | Rpb3 | AGH03_4 | 10,978,154 | 7,859,118 | 0.996 | 0.996 | | (Qiu et al. 2016) |
| WT_I | Rpb3 | AGH03_5 | 16,862,702 | 11,526,037 | | 0.995 | | (Qiu et al. 2016) |
| WT_I | Rpb3 | AGH03_6 | 14,660,604 | 10,042,576 | | | | (Qiu et al. 2016) |
| | | | | | AGH12_8 | AGH28_1 | | |
| *snf2Δ* _I | Rpb3 | AGH12_7 | 14213867 | 10131580 | 0.995 | 0.938 | | (Qiu et al. 2016) |
| *snf2Δ* _I | Rpb3 | AGH12_8 | 12857958 | 8200570 | | 0.950 | | (Qiu et al. 2016) |
| *snf2Δ* _I | Rpb3 | AGH28_1 | 13651250 | 1859651 | | | | (Qiu et al. 2016) |
| | | | | | AGH51_05 | AGH51_06 | | |
| $P_{TET}$*STH1*_I | Rpb3 | AGH51_01 | 6836169 | 1768416 | 0.993 | 0.990 | | (Rawal, Chereji, Qiu, et al. 2018) |
| $P_{TET}$*STH1*_I | Rpb3 | AGH51_05 | 12990191 | 2395951 | | 0.992 | | (Rawal, Chereji, Qiu, et al. 2018) |
| $P_{TET}$*STH1*_I | Rpb3 | AGH51_06 | 13376423 | 2103356 | | | | (Rawal, Chereji, Qiu, et al. 2018) |
| | | | | | AGH51_09 | AGH51_10 | | |
| *snf2Δ* $P_{TET}$*STH1*_I | Rpb3 | AGH51_02 | 12853198 | 1909934 | 0.987 | 0.986 | | (Rawal, Chereji, Qiu, et al. 2018) |
| *snf2Δ* $P_{TET}$*STH1*_I | Rpb3 | AGH51_09 | 11798395 | 1996785 | | 0.990 | | (Rawal, Chereji, Qiu, et al. 2018) |
| *snf2Δ* $P_{TET}$*STH1*_I | Rpb3 | AGH51_10 | 12653664 | 2171622 | | | | (Rawal, Chereji, Qiu, et al. 2018) |
| | | | | | AGH64-11 | AGH64-12 | | |
| *ino80Δ* _I | Rpb3 | AGH64-10 | 10,776,203 | 5,019,827 | 0.9943 | 0.9937 | | (Qiu et al. 2020) |
| *ino80Δ* _I | Rpb3 | AGH64-11 | 8,847,337 | 4,762,328 | | 0.9956 | | (Qiu et al. 2020) |

| | | | | | | | | |
|---|---|---|---|---|---|---|---|---|
| *ino80Δ* _I | Rpb3 | AGH64-12 | 8,315,372 | 3,731,210 | | | | (Qiu et al. 2020) |

#### Table S2 (cont'd):

##### V. Sonication H3 (SC_H3) ChIP-seq

| Strain | IP | Sample ID | All PE reads | PE rmdup | Pearson correlation between replicates | | | | | Source |
|---|---|---|---|---|---|---|---|---|---|---|
| | | | | | AGH0220-2 | AGH0220-3 | AGH58_02 | AGH62_01 | AGH62_02 | |
| WT_U | SC_H3 | AGH0220-1 | 12,305,230 | 11,468,229 | 0.965 | 0.967 | 0.850 | 0.907 | 0.911 | (Qiu et al. 2016) |
| WT_U | SC_H3 | AGH0220-2 | 11,725,253 | 10,818,529 | | 0.964 | 0.850 | 0.904 | 0.904 | (Qiu et al. 2016) |
| WT_U | SC_H3 | AGH0220-3 | 12,511,668 | 11,629,718 | | | 0.868 | 0.916 | 0.915 | (Qiu et al. 2016) |
| WT_U | SC_H3 | AGH58_02 | 29625671 | 8011769 | | | | 0.923 | 0.908 | (Rawal, Chereji, Qiu, et al. 2018) |
| WT_U | SC_H3 | AGH62_01 | 14993257 | 8597216 | | | | | 0.966 | (Rawal, Chereji, Qiu, et al. 2018) |
| WT_U | SC_H3 | AGH62_02 | 19032734 | 9965205 | | | | | | (Rawal, Chereji, Qiu, et al. 2018) |
| | | | | | AGH0220-5 | AGH0220-6 | AGH58-04 | AGH62_03 | AGH62_08 | |
| WT_I | SC_H3 | AGH0220-4 | 9,947,964 | 7,013,803 | 0.947 | 0.939 | 0.903 | 0.937 | 0.915 | (Qiu et al. 2016) |
| WT_I | SC_H3 | AGH0220-5 | 10,810,351 | 10,096,814 | | 0.968 | 0.872 | 0.930 | 0.901 | (Qiu et al. 2016) |
| WT_I | SC_H3 | AGH0220-6 | 11,687,103 | 10,849,724 | | | 0.850 | 0.916 | 0.884 | (Qiu et al. 2016) |
| WT_I | SC_H3 | AGH58-04 | 20,620,578 | 5,466,934 | | | | 0.936 | 0.933 | (Rawal, Chereji, Qiu, et al. 2018) |
| WT_I | SC_H3 | AGH62-03 | 18,617,080 | 8,200,524 | | | | | 0.951 | (Rawal, Chereji, Qiu, et al. 2018) |
| WT_I | SC_H3 | AGH62-08 | 19,798,491 | 5,044,590 | | | | | | (Rawal, Chereji, Qiu, et al. 2018) |
| | | | | | AGH04_02 | AGH25_01 | AGH59_01 | AGH59_02 | AGH62_05 | |
| *snf2Δ* _I | SC_H3 | AGH04_01 | 15597956 | 11985536 | 0.963 | 0.881 | 0.905 | 0.897 | 0.870 | (Qiu et al. 2016) |
| *snf2Δ* _I | SC_H3 | AGH04_02 | 17071372 | 13099928 | | 0.871 | 0.916 | 0.907 | 0.883 | (Qiu et al. 2016) |
| *snf2Δ* _I | SC_H3 | AGH25_01 | 24839200 | 11548218 | | | 0.796 | 0.799 | 0.720 | (Qiu et al. 2016) |
| *snf2Δ* _I | SC_H3 | AGH59_01 | 14174694 | 7052399 | | | | 0.951 | 0.947 | (Rawal, Chereji, Qiu, et al. 2018) |
| *snf2Δ* _I | SC_H3 | AGH59_02 | 14573693 | 6456125 | | | | | 0.945 | (Rawal, Chereji, Qiu, et al. 2018) |
| *snf2Δ* _I | SC_H3 | AGH62_05 | 20877009 | 10613818 | | | | | | (Rawal, Chereji, Qiu, et al. 2018) |
| | | | | | AGH50_06 | AGH54_03 | | | | |
| $P_{TET}$*STH1*_I | SC_H3 | AGH50_01 | 23176098 | 12629620 | 0.962 | 0.970 | | | | (Rawal, Chereji, Qiu, et al. 2018) |
| $P_{TET}$*STH1*_I | SC_H3 | AGH50_06 | 26443231 | 6284621 | | 0.971 | | | | (Rawal, Chereji, Qiu, et al. 2018) |
| $P_{TET}$*STH1*_I | SC_H3 | AGH54_03 | 22858502 | 9011007 | | | | | | (Rawal, Chereji, Qiu, et al. 2018) |
| | | | | | AGH54_01 | AGH54_02 | | | | |
| *snf2Δ* $P_{TET}$*STH1*_I | SC_H3 | AGH50_10 | 30027139 | 5408247 | 0.863 | 0.881 | | | | (Rawal, Chereji, Qiu, et al. 2018) |
| *snf2Δ* $P_{TET}$*STH1*_I | SC_H3 | AGH54_01 | 23674356 | 12035478 | | 0.951 | | | | (Rawal, Chereji, Qiu, et al. 2018) |
| *snf2Δ* $P_{TET}$*STH1*_I | SC_H3 | AGH54_02 | 27937517 | 13179773 | | | | | | (Rawal, Chereji, Qiu, et al. 2018) |
| | | | | | AGH61-5 | AGH61-6 | | | | |

| | | | | | | | | | | |
|---|---|---|---|---|---|---|---|---|---|---|
| *ino80Δ* _I | SC_H3 | AGH61-4 | 16,858,717 | 8,430,284 | 0.942 | 0.942 | | | | (Qiu et al. 2020) |
| *ino80Δ* _I | SC_H3 | AGH61-5 | 18,996,243 | 9,997,136 | | 0.947 | | | | (Qiu et al. 2020) |
| *ino80Δ* _I | SC_H3 | AGH61-6 | 19,488,986 | 10,065,366 | | | | | | (Qiu et al. 2020) |

#### Table S2 (cont'd):

##### VI. MNase H3 (MN_H3) ChIP-seq

| Strain | IP | Sample ID | All PE reads | PE rmdup | Pearson correlation between replicates | | | | | Source |
|---|---|---|---|---|---|---|---|---|---|---|
| | | | | | AGH68_02 | AGH73_01 | AGH73_02 | | | |
| WT_U | MN_H3 | AGH68_01 | 21155789 | 16387173 | 0.975 | 0.939 | 0.971 | | | (Rawal, Chereji, Qiu, et al. 2018) |
| WT_U | MN_H3 | AGH68_02 | 29890860 | 21290074 | | 0.941 | 0.975 | | | (Rawal, Chereji, Qiu, et al. 2018) |
| WT_U | MN_H3 | AGH73_01 | 22748400 | 13752475 | | | 0.956 | | | (Rawal, Chereji, Qiu, et al. 2018) |
| WT_U | MN_H3 | AGH73_02 | 33931620 | 20843860 | | | | | | (Rawal, Chereji, Qiu, et al. 2018) |
| | | | | | AGH68_04 | AGH73_03 | | | | |
| WT_I | MN_H3 | AGH68_03 | 22985503 | 17204691 | 0.957 | 0.926 | | | | (Rawal, Chereji, Qiu, et al. 2018) |
| WT_I | MN_H3 | AGH68_04 | 19959153 | 15376225 | | 0.956 | | | | (Rawal, Chereji, Qiu, et al. 2018) |
| WT_I | MN_H3 | AGH73_03 | 26493831 | 17942518 | | | | | | (Rawal, Chereji, Qiu, et al. 2018) |
| | | | | | AGH68_06 | AGH73_04 | AGH73_05 | AGH73_10 | | |
| *snf2Δ* _I | MN_H3 | AGH68_05 | 20896164 | 15789365 | 0.964 | 0.961 | 0.966 | 0.949 | | (Rawal, Chereji, Qiu, et al. 2018) |
| *snf2Δ* _I | MN_H3 | AGH68_06 | 23587865 | 18055776 | | 0.972 | 0.965 | 0.965 | | (Rawal, Chereji, Qiu, et al. 2018) |
| *snf2Δ* _I | MN_H3 | AGH73_04 | 30193725 | 20155496 | | | 0.977 | 0.972 | | (Rawal, Chereji, Qiu, et al. 2018) |
| *snf2Δ* _I | MN_H3 | AGH73_05 | 28834376 | 19622426 | | | | 0.964 | | (Rawal, Chereji, Qiu, et al. 2018) |
| *snf2Δ* _I | MN_H3 | AGH73_10 | 27351420 | 18319655 | | | | | | (Rawal, Chereji, Qiu, et al. 2018) |
| | | | | | AGH68_08 | AGH73_06 | AGH73_07 | AGH73_12 | | |
| $P_{TET}$*STH1*_I | MN_H3 | AGH68_07 | 21614150 | 16939365 | 0.980 | 0.974 | 0.960 | 0.933 | | (Rawal, Chereji, Qiu, et al. 2018) |
| $P_{TET}$*STH1*_I | MN_H3 | AGH68_08 | 24081014 | 18755148 | | 0.977 | 0.964 | 0.930 | | (Rawal, Chereji, Qiu, et al. 2018) |
| $P_{TET}$*STH1*_I | MN_H3 | AGH73_06 | 29619041 | 19719008 | | | 0.979 | 0.958 | | (Rawal, Chereji, Qiu, et al. 2018) |
| $P_{TET}$*STH1*_I | MN_H3 | AGH73_07 | 26053659 | 18067975 | | | | 0.971 | | (Rawal, Chereji, Qiu, et al. 2018) |
| $P_{TET}$*STH1*_I | MN_H3 | AGH73_12 | 24618600 | 16927985 | | | | | | (Rawal, Chereji, Qiu, et al. 2018) |
| | | | | | AGH68_10 | AGH68_11 | AGH68_12 | AGH73_08 | AGH73_09 | |
| *snf2Δ* $P_{TET}$*STH1*_I | MN_H3 | AGH68_09 | 24352916 | 18823892 | 0.961 | 0.975 | 0.911 | 0.968 | 0.967 | (Rawal, Chereji, Qiu, et al. 2018) |
| *snf2Δ* $P_{TET}$*STH1*_I | MN_H3 | AGH68_10 | 35625027 | 24578682 | | 0.949 | 0.897 | 0.947 | 0.943 | (Rawal, Chereji, Qiu, et al. 2018) |
| *snf2Δ* $P_{TET}$*STH1*_I | MN_H3 | AGH68_11 | 26104427 | 19584798 | | | 0.913 | 0.966 | 0.969 | (Rawal, Chereji, Qiu, et al. 2018) |
| *snf2Δ* $P_{TET}$*STH1*_I | MN_H3 | AGH68_12 | 19106755 | 11875757 | | | | 0.883 | 0.891 | (Rawal, Chereji, |

| | | | | | | | | | | |
|---|---|---|---|---|---|---|---|---|---|---|
| | | | | | | | | | | Qiu, et al. 2018) |
| *snf2Δ* $P_{TET}$*STH1*_I | MN_H3 | AGH73_08 | 27010389 | 18495503 | | | | | 0.976 | (Rawal, Chereji, Qiu, et al. 2018) |
| *snf2Δ* $P_{TET}$*STH1*_I | MN_H3 | AGH73_09 | 25011131 | 17657515 | | | | | | (Rawal, Chereji, Qiu, et al. 2018) |
